# Seven amino acid types suffice to reconstruct the core fold of RNA polymerase

**DOI:** 10.1101/2021.02.22.432383

**Authors:** Sota Yagi, Aditya K. Padhi, Jelena Vucinic, Sophie Barbe, Thomas Schiex, Reiko Nakagawa, David Simoncini, Kam Y. J. Zhang, Shunsuke Tagami

**Affiliations:** RIKEN Center for Biosystems Dynamics Research, 1-7-22 Suehiro-cho, Tsurumi-ku, Yokohama, Kanagawa 230-0045, Japan; Université Fédérale de Toulouse, ANITI, INRAE - UR 875, Toulouse, France; TBI, Université Fédérale de Toulouse, CNRS, INRAE, INSA, ANITI, Toulouse, France; Université Fédérale de Toulouse, ANITI, IRIT - UMR 5505, Toulouse, France.

## Abstract

The extant complex proteins must have evolved from ancient short and simple ancestors. Nevertheless, how such prototype proteins emerged on the primitive earth remains enigmatic. The double-psi beta-barrel (DPBB) is one of the oldest protein folds and conserved in various fundamental enzymes, such as the core domain of RNA polymerase. Here, by reverse engineering a modern DPBB domain, we reconstructed its evolutionary pathway started by “interlacing homo- dimerization” of a half-size peptide, followed by gene duplication and fusion. Furthermore, by simplifying the amino acid repertoire of the peptide, we successfully created the DPBB fold with only seven amino acid types (Ala, Asp, Glu, Gly, Lys, Arg, and Val), which can be coded by only GNN and ARR (R = A or G) codons in the modern translation system. Thus, the DPBB fold could have been materialized by the early translation system and genetic code.

---

Modern proteins with large and complex structures are generally thought to have evolved from small and simple ancient proteins with “prototype folds” (e.g., Rossmann fold, ferredoxin fold, and (*β/α*)8-barrel) (*1–10*). These prototype folds must have played essential roles in the early evolution of life, as they are often conserved in fundamental biochemical pathways such as metabolism, replication, transcription, and translation (*11–14*). However, it remains elusive how such prototype folds emerged on the ancient earth, where the primitive translation system likely performed imprecise syntheses of short peptides composed of fewer amino acids as compared to modern proteins (*15–17*). Especially, the components of the earliest genetic code are still an open question, as the 9–13 amino acid types used in previous ancestral protein reconstructions are at scattered positions in the modern codon table (*18–21*).

The double-psi beta-barrel (DPBB) has one of the most important functions and complicated structures among such prototype folds. It is conserved in several enzymes in fundamental biochemical processes (Fig. S1) (*13, 22–24*). For example, the formate dehydrogenase in the oldest carbon fixation system, the Wood-Ljungdahl pathway, possesses a DPBB domain (*25*). The DPBB and a few related small *β*-barrel folds are also often conserved in essential proteins from transcription and translation systems (*26, 27*). Most notably, the active site of RNA polymerase from all cellular life is composed of two DPBB folds, and thus the original core of RNA polymerase may have emerged by the duplication of an ancestral DPBB (*14, 28–30*).

The DPBB fold is a six-stranded *β*-barrel consisting of two pseudo-symmetric *ββαβ* units with detectable structural and sequence homologies (Fig. 1A). All six *β*-strands are aligned in an interdigitated manner, giving its pseudoknot-like topology. The loop connecting *β*1 and *β*2 (*β*1’ and *β*2’) crosses the loop between *β*2’ and *β*3’ (*β*2 and *β*3), thus generating a shape similar to the Greek letter *ψ*. Although some other prototype folds with similar pseudo-symmetries apparently originated by oligomerization and gene duplication of shorter peptides (*31*), it remains uncertain if the interdigitating fold of DPBB could have formed in such a simple oligomerization process (*13, 22, 27, 29*). So far, no modern DPBB structure composed of a perfect sequence repeat or a dimer of shorter peptides has been reported, and thus the origin of the DPBB fold remains elusive. In this study, we demonstrate that the DPBB fold could have emerged by oligomerization and gene-duplication of a shorter and simpler peptide, by reconstructing the DPBB domains comprising perfect sequence repeats and then self-dimerizing peptides. Surprisingly, even chemically synthesized peptides could fold into the complicated pseudoknot-like topology. We also eliminated several amino acid types from the designs, and confirmed that only seven amino acid types (Ala, Asp, Glu, Gly, Lys, Arg, and Val) are sufficient for the DPBB fold. These amino acid types can be coded by only GNN and ARR (R = A or G) codons in the modern translation system. These results reveal the plausible ancient pathway for the emergence of the complicated prototype fold and transcription machinery, which coevolved with the early translation system and genetic code.

**Figure 1.**
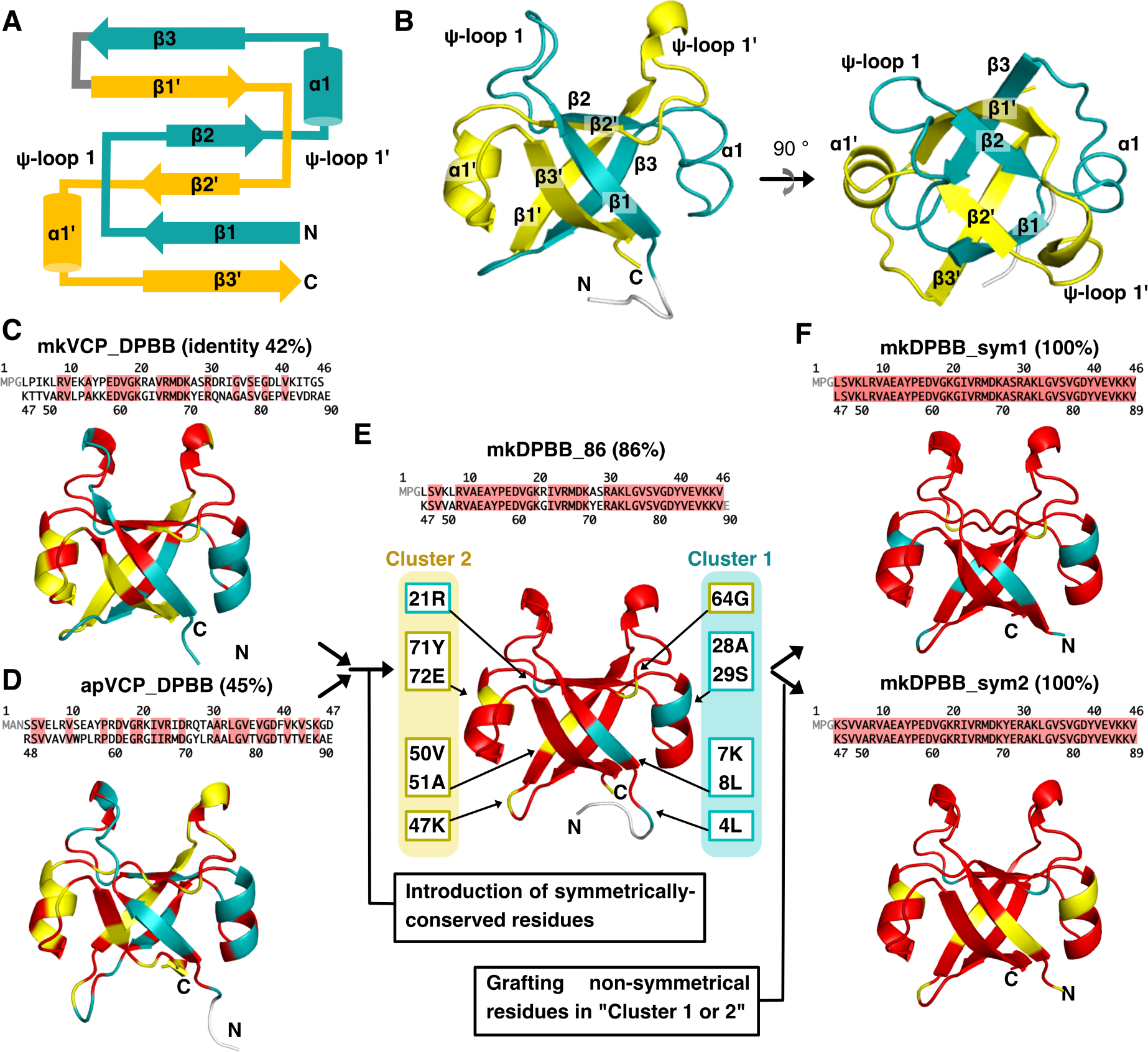
Structures and symmetrical engineering of the DPBB domains. **(A)** Topology diagram of the DPBB fold. The N- and C-terminal *ββαβ* units are colored cyan and yellow. **(B)** The crystal structure of taVCP_DPBB. Both halves have the same color as in the topology diagram of panel A. The right view shows the left view rotated by 90 degrees around the horizontal axis. **(C–F)** Symmetric-conservation design (SC-design). The amino acid sequence and crystal structure of (C) mkVCP_DPBB, (D) apVCP_DPBB, (E) mkDPBB_86, and (F) mkDPBB_sym1 and 2 are shown in each panel. The conserved residues in both halves are highlighted in red. (E) mkDPBB_86 has only a limited number of non-symmetric positions, which are grouped in two areas (clusters 1 and 2).

## Reconstruction of DPBB domains with perfect sequence symmetry

We first searched for extant DPBB domains with high internal sequence homologies, as the starting models to reconstruct a DPBB domain with perfect sequence repeats. The DPBB domain of a molecular chaperone, valosin-containing protein (VCP) from *Thermoplasma acidophilum* (taVCP_DPBB), reportedly has relatively high sequence identity (37%) between its N- and C-terminal halves (Table S1, Fig. S1F) (*22*). We determined its crystal structure in the isolated form, to confirm that it adopts the DPBB fold even without the other domains of VCP (Fig. 1B, Table S2). taVCP_DPBB also shares high structural homology with the core domain of RNA polymerase (RMSD = 1.6 Å), indicating their common origin (Fig. S1G).

**Table 1.** Summary of experimental tests for three different design strategies.

| | Tested designs | Express <sup>a</sup> | Soluble <sup>a</sup> | $\alpha/\beta$ structure | Monomeric <sup>b</sup> | Crystal structure |
| --- | --- | --- | --- | --- | --- | --- |
| SC-design | 4 | 4 | 3 | 3 | 3 | 2 |
| RE-design | 7 | 5 | 5 | 5 | 5 | 3 |
| MS-design | 2 | 2 | 2 | 2 | 2 | 1 |

<sup>a</sup> Expressions and solubilities were examined by SDS-PAGE.
<sup>b</sup> Oligomeric states were examined by size exclusion chromatography.

Using the sequence of taVCP_DPBB as a query, we searched for VCP_DPBB domains with higher internal sequence homology from other organisms and found that the VCP_DPBBs from *Methanopyrus kandleri* (mkVCP_DPBB) and *Aeropyrum pernix* (apVCP_DPBB) have 42% and 45% internal sequence identities, respectively (Tables S1, S3). We determined their crystal structures (Fig. 1C and D, Table S2) and confirmed that they also exhibit more precise structural symmetries than taVCP_DPBB (Fig. S2). Thus, these two VCP_DPBBs were estimated to have similar sequences and structures to the ancestral DPBB with perfect sequence repeats. Furthermore, mkVCP_DPBB and apVCP_DPBB were from extreme thermophiles and showed high thermostability (*Tm* > 69 °C) and refolding ability (Fig. S3 and Table S4). The coexistence of the ancestral symmetric feature and the extreme thermostability in these DBPP domains is also consistent with the widely supported hypothesis suggesting that the last universal common ancestor (LUCA) was a hyper-thermophilic organism (*32, 33*).

We then reconstructed DPBB domains with perfect internal repeats, by a method based on symmetrically-conserved positions (SC-design, Supplementary Text, Figs. S4–6, Tables S1, S2, S5). We engineered mkVCP_DPBB and apVCP_DPBB by replacing the residues in their non-symmetric positions with the symmetrically-conserved residues in VCP_DPBBs from different organisms. A few cycles of mutagenesis and structural confirmation by circular dichroism (CD), SEC, and X-ray crystallography finally resulted in two mutants, mkDPBB_sym_86 and apDPBB_sym_84, with 86% and 84% internal sequence identities, respectively (Fig. 1E and Table S1). These designs only have a limited number of non-symmetric positions clustered in two areas in their tertiary structures, defined as “cluster-1 and -2” in Fig. 1E (also see Fig. S7). By grafting the amino acid residues from each of the two clusters to the other one, we then designed four DPBBs with perfect internal sequence identities (mkDPBB_sym1, mkDPBB_sym2, apDPBB_sym1, and apDPBB_sym2, Table S1). Although one of them (apDPBB_sym2) could not be purified due to its poor stability, the other three were readily purified. They eluted as monomeric proteins in SEC analyses, exhibited similar *α/β* CD spectra with the native DPBB proteins, and retained high thermostability (*Tm* ≥ 85°C, Fig. S5, Table S5). Finally, we solved the crystal structures of mkDPBB_sym1 and mkDPBB_sym2 (Fig. 1F, Table S2), which have almost perfect structural symmetries. These results strongly indicate that the DPBB fold originally arose from a perfectly symmetric ancestor.

## Computational designs and possible sequence diversity of symmetric DPBBs

We also implemented two computational approaches for symmetric designs of DPBB to test if diverse strategies could result in different/similar sequences. The first computational approach was a modified “reverse engineering evolution” (*34*) (RE-design, Supplementary Text, Fig. 2A, Fig. S8). In this methodology, we constructed a phylogenetic tree with the respective aligned sequences, which were subsequently used as input to generate the ancestral sequences. The predicted ancestral sequences were mapped onto a manually constructed, perfectly symmetrical mkVCP structural backbone model, and were evaluated by their Rosetta energy scores (Fig. S9). The seven top-scoring designs were chosen for the experimental evaluation (Table S1).

**Figure 2.**
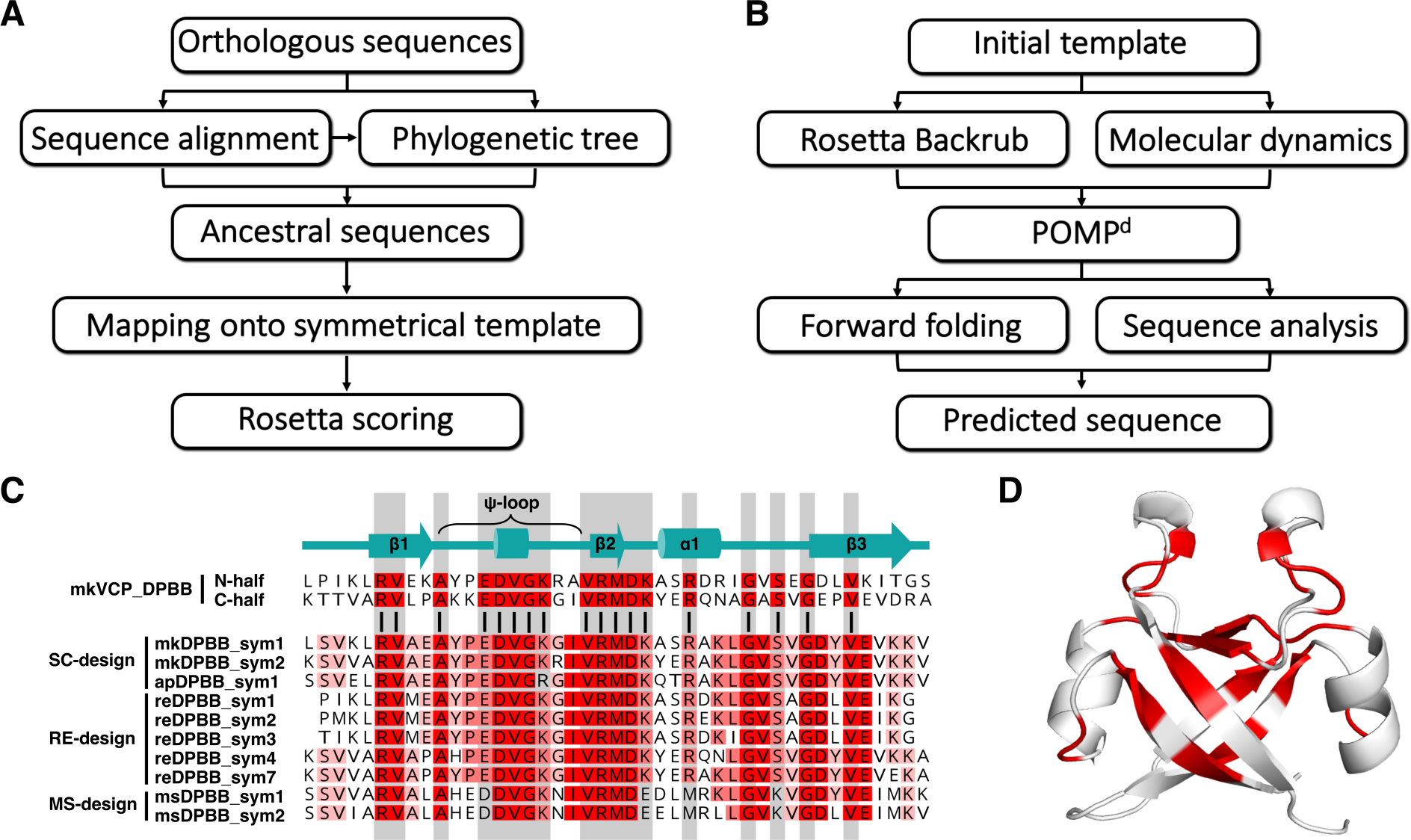
Computational design of symmetric DPBB and diversity of the symmetric DPBBs. **(A)** Flow-chart of the RE-design scheme. **(B)** Flow-chart of the MS-design scheme. **(C)** Multiple sequence alignment of one repeat unit in the symmetric designs along with the mkVCP_DPBB, the starting template DPBB. The columns corresponding to the consensus residues in mkVCP_DPBB are highlighted in gray. The perfectly identical columns in the symmetric designs are colored red, and the columns with sequence identities over 60% are colored pink. **(D)** The perfectly conserved residues among all experimentally confirmed symmetric designs are mapped on the crystal structure of reDPBB_sym1.

The second computational approach was a multi-state computational protein design (MS-design, Methods, Fig. 2B, Fig. S10). This method describes the target protein structure as an ensemble of fixed backbone conformational states, to account for protein flexibility (*35*). The sequence that minimizes the average energy over all conformational states is computed by the AI automated reasoning prover ToulBar2 (*36*). The MS-designs were less homologous to the other designs since they were obtained by energy minimization, without any knowledge extraction from other homologous DPBB domains. From the top-scoring designs, we chose the two with the highest sequence dissimilarities from the SC- or RE-designs for the experimental evaluation (Fig. S11, Table S1).

We tried to express the seven RE-designs and two MS-designs in *E. coli*. Although two RE- designs were not expressed well, the other designs were readily purified, exhibited similar properties to mkVCP_DPBB in SEC and CD analyses, and retained high thermostability (*Tm* ≥ 67 °C, Figs. S12 and S13, Table S5). Furthermore, we solved the crystal structures of three RE- designs and one MS-design and confirmed that they adopt the designed structures (Fig. 2D, Fig. S6B–E, Table S2).

Interestingly, while all ten purified designs (three SC-designs and seven computational designs) have almost identical properties (structure, thermostability), their sequences are very diverse (Fig. 2C and Fig. S11, Supplementary Text). Only 34 positions (17 positions × 2 repeats) were perfectly conserved in the ∼90 a.a. designs (Fig. 2D). The high sequence diversity and success ratios of these designs (Table 1) indicate that the symmetric DPBB fold can be adopted by a significant variety of sequences and had a high probability of emerging during the early evolution of life.

## Homo-dimerization of halved fragments

To further investigate if the DPBB structure can be formed by the homo-dimerization of halved fragments (∼46 aa) of symmetric DPBBs, we expressed the N-terminal halves of the four SC- designs (mk1h, mk2h, ap1h, and ap2h; Fig. 3A and 3B and Table S1). All four fragments were expressed as soluble peptides in *E. coli* and formed dimers with *α/β* structures (Fig. S14 and Table S6). Crystallographic analyses of mk2h and ap1h demonstrated that the halved fragments adopt the DPBB fold by interlacing homo-dimerization (Fig. 3C, Fig. S6F and Table S2). They also exhibited high thermostability and refoldability after heat denaturation (Fig. 3D, 3E, Fig. S14 and Table S6). These results strongly indicate that the DPBB fold originally emerged simply by the homo-dimerization of a short peptide, in spite of its complicated interlaced topology.

**Figure 3.**
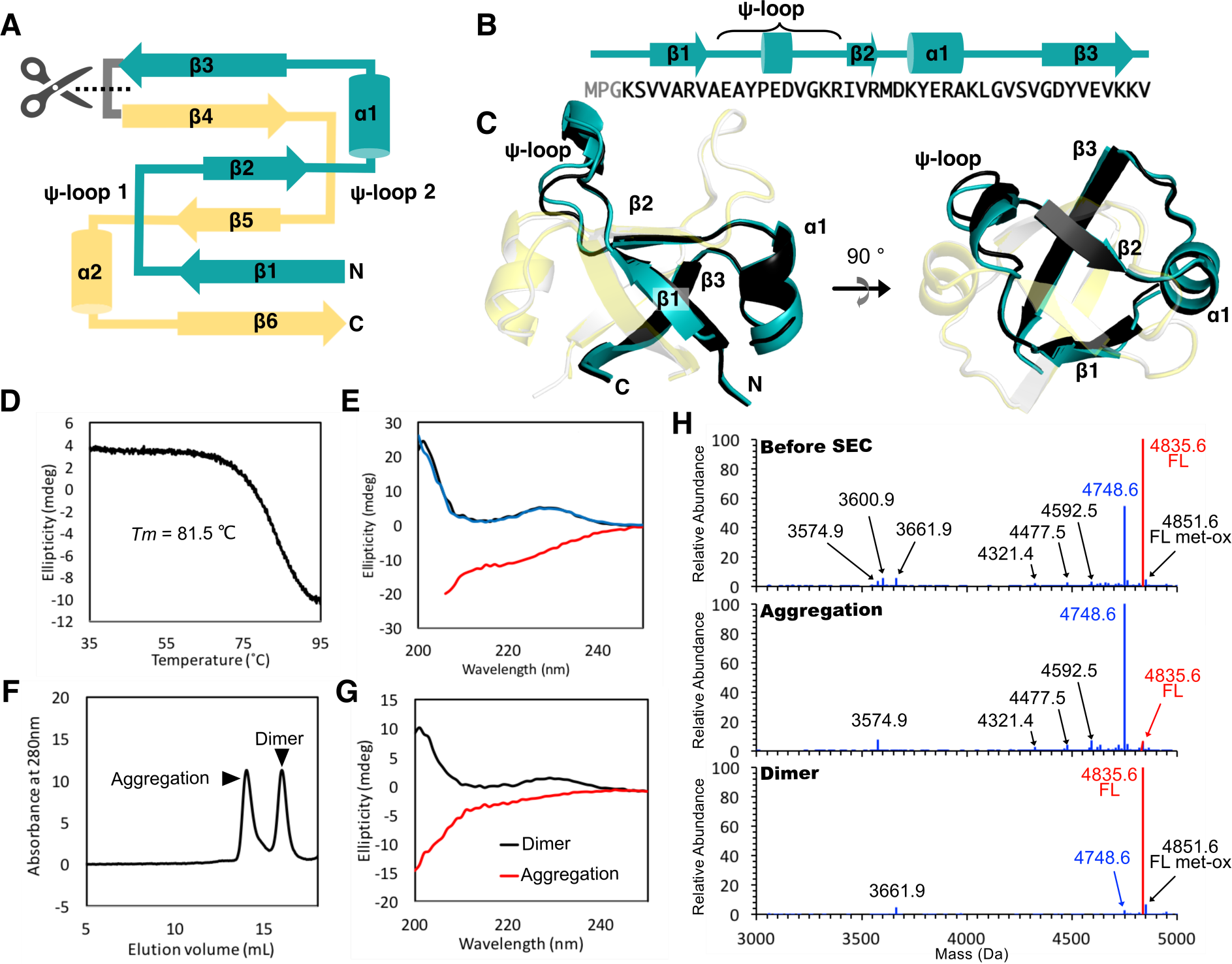
The DPBB fold formed by halved fragments. **(A)** Topology diagram of the halved fragments. **(B)** The sequence and secondary structure of mk2h. **(C)** The crystal structures of homo- dimeric mk2h. The *E. coli*-produced peptides are colored cyan and yellow. The chemically synthesized peptides are colored black and white. **(D and E)** Thermostability and refolding ability of mk2h. (D) Denaturation was monitored by measuring the CD ellipticity at 222 nm. (E) CD spectra at 35 °C (black line), after heating to 95 °C (red line), and upon cooling back to 35 °C (blue line). **(F–H)** Folding of the low-purity mk2h sample (71.16%). (F) SEC analysis showing that the dissolved mk2h peptide adopted aggregated and dimeric states. (G) CD spectra indicating that the aggregated and dimeric species, separated in Fig. 3F, adopt random-coil and *α/β* structures, respectively. (H) The peptide species in the sample before and after SEC purification were analyzed by LC/MS. The deconvoluted mass spectra are shown. The labels for the full-length mk2h peptide (4835.6 Da) and the major contaminant peptide (4748.6 Da) are highlighted in red and blue, respectively.

We also tested the foldability of the chemically-synthesized mk2h peptide. First, the dried powder of mk2h (95.14% purity) was dissolved in 20 mM Bis-tris HCl, pH 6.0, with 150 mM NaCl, and subjected to crystallization screening. We readily obtained some crystals under various conditions (Table S7) and determined its structure to confirm that it also adopts the DPBB structure (Fig. 3C, Table S2). Thus, the chemically-synthesized peptide can fold precisely into the interlacing DPBB structure without any factors/environment in the cell, demonstrating that the amino acid sequence of mk2h encodes its homo-dimerizing and folding information.

Next, to investigate whether the peptide could fold even in the presence of contaminants with similar sequences, we analyzed the foldability of a low-purity sample of mk2h containing byproducts from the chemical synthesis (71.16% purity). In the SEC analysis, two major peaks corresponding to aggregated and dimer species appeared (Fig. 3F). The dimer fraction showed the typical CD spectra for *α/β* proteins, while the aggregated fraction exhibited a disordered conformation (Fig. 3G). The LC/MS analysis revealed that most of the contaminants were enriched in the aggregated fraction (e.g., the 4748.6 Da byproduct corresponding to a serine deletion, Fig. 3H and Table S8). In contrast, the full-length peptide (4835.6 Da) was enriched in the dimer fraction. This auto-purification phenomenon during protein folding was also observed with another design, mk1h (Fig. S15 and Table S8). Therefore, the homo-dimerization and folding processes of the peptides likely worked as a purification/selection system by excluding contaminated sequences, and thus might have enabled the production of the DPBB domains by an imprecise ancient translation system or prebiotic peptide synthesis.

## Reduction of amino acid repertory

Interestingly, mk2h contained only 13 amino acid types, although we did not intend to enrich/exclude any specific amino acid species in the engineering process (Fig. 4A, Table S9). To examine how many and what kind of amino acid species are required to comprise a DPBB scaffold, we tried to further simplify the amino acid repertoire of mk2h (Fig. 4B). In mk2h, Ile, Leu, Met, Pro, and Ser were used only once or twice. Tyrosine was the only aromatic amino acid type and used just three times (Fig. 4A). Thus, we replaced each of these amino acid types with other amino acid residues conserved in different organisms or possessing similar chemical/structural properties (e.g., Ile to Val) to generate mutants containing 12 amino acid repertoires (Table S1). MD simulations estimated that none of the mutations would have a devastating effect on the domain structure (Fig. S16). All mutants were eluted as homo-dimers in SEC and their CD spectra were similar to that of mk2h (Fig. S17), although some of them exhibited lower *Tm* values than mk2h (Table S10). Furthermore, the crystal structures of mk2h_ΔP and mk2h_ΔY revealed they indeed adopt the DPBB fold (Fig. S6G and H, Table S2). Therefore, these amino acid species may have contributed to the thermostability of mk2h, but are not essential to form the DPBB fold.

**Figure 4.**
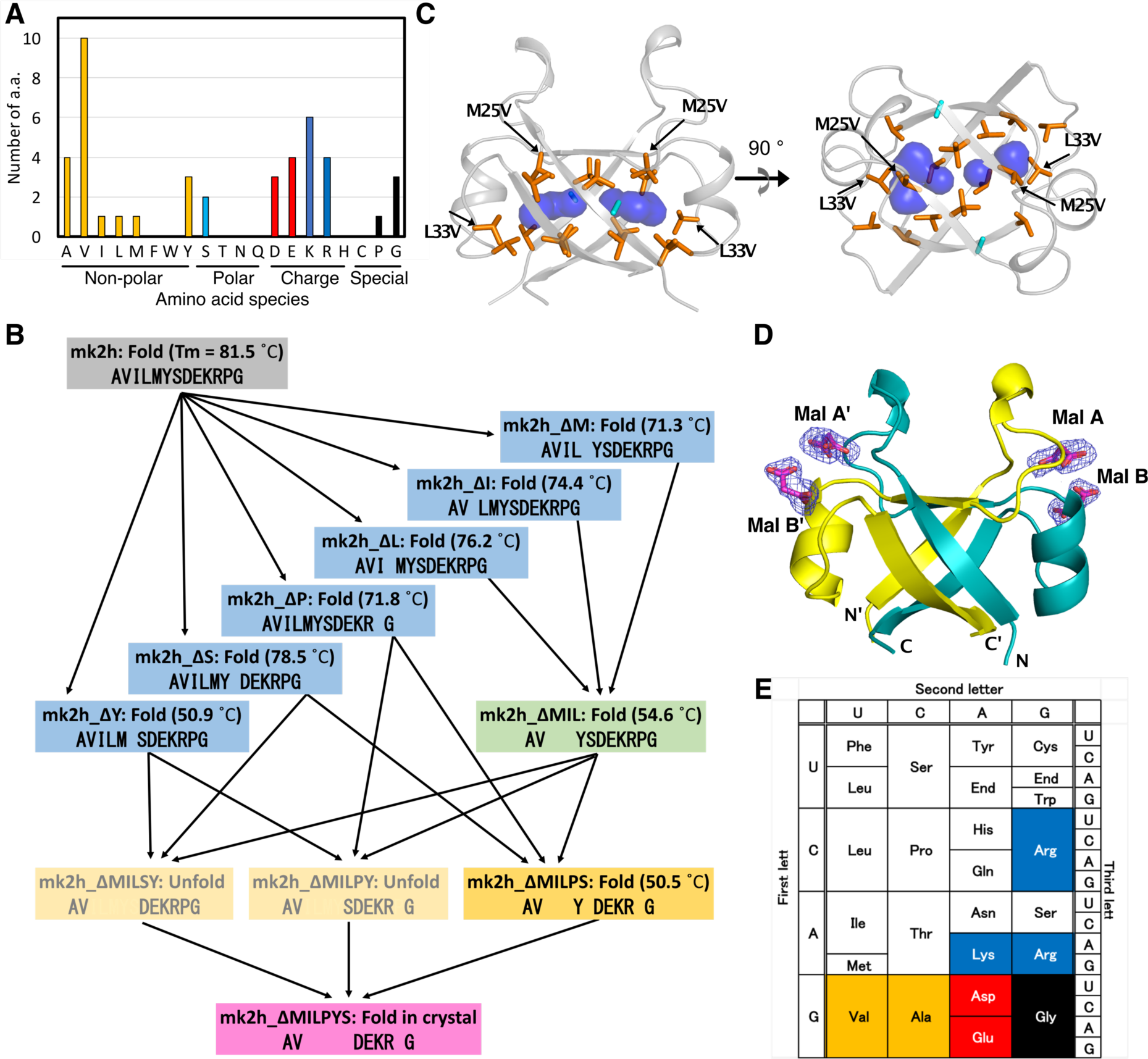
Reduction of amino acid repertory in mk2h. **(A)** The amino acid usage in mk2h. **(B)** Scheme of the design process of the simplified mk2h variants. **(C)** The hydrophobic core of mk2h_ΔMIL, composed of only Val (orange) and Ala (cyan). The substituted residues (M25V and L33V) from mk2h are indicated. The cavities in the core are shown as blue surface models. **(D)** The crystal structure of mk2h_ΔMILPYS. Four malonate ions, Mal A (A’) and B (B’) are bound to two symmetrical positions. The malonates are shown by stick models, and their Fo-Fc electron density omit maps are presented as a blue mesh (contoured at 1.0σ). **(E)** The seven amino acid species used in mk2h_ΔMILPYS are highlighted on the standard codon table.

Subsequently, we created a mutant in which Met, Ile, and Leu were eliminated simultaneously (mk2h_ΔMIL). The mutant was expressed as a homo-dimer and showed typical CD spectra for *α/β* proteins (*Tm* = 54.6 °C, Fig. S18A and Table S10). The X-ray crystallographic analysis of mk2h_ΔMIL revealed it has a very primitive hydrophobic core composed of only Val and Ala, with some unfilled cavities (Fig. 4C). Recently, a *de novo* designed protein with a hydrophobic core composed mostly of Valine residues was reported, indicating that the backbone structure is the main contributor to its thermostability (*37*). GD-box, a structural motif conserved among DPBB and various protein folds, has also been suggested to stabilize the protein tertiary structures by tethering noncontiguous segments with hydrogen bonds between the main chains (*38*). Such a stable backbone of the DPBB fold would have allowed the emergence of a globular protein with only simple and small hydrophobic amino acids during early protein evolution. Unlike hydrophilic interactions, hydrophobic interactions do not require precise angles or directions, and thus such simple hydrophobic cores might emerge relatively easily in aqueous environments without optimizing the sizes or compositions of their amino acid residues.

We also tried to exclude two amino acid types (Pro/Ser, Pro/Tyr, or Ser/Tyr) in mk2h_ΔMIL (mk2h_ΔMILPS, mk2h_ΔMILPY and mk2h_ΔMILSY; Fig. 4B and Table S1). While mk2h_ΔMILPY and mk2h_ΔMILSY were eluted as unfolded aggregates in SEC, mk2h_ΔMILPS eluted as a dimer and its CD spectra were similar to those of mk2h (Fig. S18B–D and Table S10). The crystal structures of *E. coli*-produced and chemically-synthesized mk2h_ΔMILPS were also determined (Fig. S6I and J, Table S2). Although mk2h_ΔMILPS remained thermostable (*Tm* = 50.5 °C), refoldability was not observed (Fig. S18B).

Finally, we designed an mk2h variant, mk2h_ΔMILPYS, with only seven amino acid types (Ala, Asp, Glu, Gly, Lys, Arg, and Val) by combining the mutations in mk2h_ΔMILPS and mk2h_ΔY (Fig. 4B). Although the peptides exhibited unfolded properties in the SEC and CD analyses (Fig. S18E), we obtained crystals from two conditions containing the chemically-synthesized peptide (Fig. S19). The crystal structures demonstrated that mk2h_ΔMILPYS adopts the DPBB structure through homo-dimerization (Fig. 4D, Fig. S6K, and Table S2). In the structures, the positively-charged pocket conserved in the extant VCP_DPBB is occupied by the fundamental metabolites, malonate or malic acid, contained in the crystallization conditions (Fig. 4D, Fig. S20), which might also have assisted in the folding of mk2h_ΔMILPYS. Thus, despite its limited folding propensity, the 43 a.a. peptide with only seven amino acid types can homo-dimerize and fold into the DPBB structure. Additional amino acids would have been incorporated for higher stability and foldability during protein evolution (Fig. 4B).

## Discussion

In this study, we demonstrated that one of the prototypic protein folds, DPBB, can be reconstructed by perfect sequence repeats (Figs. 1, 2), as well as by dimers of the halved fragments (Fig. 3), indicating that DPBB originally emerged by the self-dimerization of ∼40 a.a. peptides and then evolved via gene duplication and fusion. Chemically synthesized peptides were able to fold into the complicated interlaced topology, without any support from the modern biological machinery, and still retained thermostability and refoldability (Fig. 3). Furthermore, the success rate and diversity of our designs were surprisingly high (Fig. 2C and Fig. S11, Table 1). The fragmented peptides also exhibited auto-purification ability (Fig. 3H) and high amenability to engineering (Fig. 4B). Thus, the DPBB fold probably had a significant chance of emerging from the pool of primitive peptides synthesized by an immature/imprecise translation system on the early earth. Nature can find and utilize such a complicated protein fold as long as it is stable enough, even though in theory it appears unrealistic to the human eye.

By simplifying mk2h, we further constructed the DPBB fold with only seven amino acid types, Ala, Asp, Glu, Gly, Lys, Arg, and Val (mk2h_ΔMILPYS, Fig. 4). The previously reconstructed ancestral proteins with ∼10 amino acid types also contain most of them (*19–21, 39*), indicating that they were shared by various prototype proteins. Interestingly, these seven amino acid types can be coded on a clearly defined area in the standard codon table (GNN and ARR) (Fig. 4E). The five amino acids coded by GNN (Ala, Asp, Glu, Gly, and Val) were probably adopted into the earliest genetic code because they can be easily produced in the prebiotic environment (*16, 17, 40–42*). The other two amino acids (Arg and Lys) have cationic side chains and tend to interact with nucleic acid polymers. Arginine could have been abiotically synthesized by sharing some precursors to nucleotides (*42*). Recent studies have shown that peptide analogs enriched with such basic residues can be prebiotically synthesized and mutually stabilize RNA (*43, 44*). Simple peptides containing lysine residues could also enhance the activities of ribozymes (*45*). Considering the fact that the genetic code is realized by RNA-based machinery, arginine and lysine were probably readily recruited into the early genetic code during evolution. This idea is also supported by the fact that the core domain of RNA polymerase, the enzyme responsible for the synthesis of rRNA, mRNA, and tRNA, is composed of the DPBB fold. Like its modern descendants, the ancient symmetric DPBB fold might have interacted with nucleotide-related molecules, as the conserved positively-charged pockets on some symmetrized or simplified designs are occupied by negatively-charged ligands in their crystal structures (Fig. S20). Thus, the DPBB fold was likely established at an early evolutionary stage of the genetic code and supported the ancient RNA-based biosystem when only ∼7 amino acid types were available.

## Acknowledgements

This work is based on experiments performed at KEK (project number: 20G056) and SPring-8. The authors are grateful to the beamline staff scientists at KEK, SPring-8, and SLS. We thank Hideaki Niwa, Toshiaki Hosaka, and Kentaro Ihara for helping with the X-ray diffraction experiments. We also thank Shigehiro Kuraku for assistance in the LC-MS analysis. We are deeply grateful to Ryutaro Furukawa for providing an informative in-house protein database to search for natural DPBB proteins. We acknowledge RIKEN ACCC for the supercomputing resources at the Hokusai BigWaterfall supercomputer used in this study. A.K.P. acknowledges the Japan Society for the Promotion of Science (JSPS), Govt. of Japan, for the research fellowship. S.Y., K.Y.J.Z. and S.T. were supported by JSPS (20K15854, 18H02395 and 18H01328). This work was also supported by the French ANR through an ANR-19-PI3A-0004 grant. We thank the CALMIP HPC center for computational resources. We thank Satoshi Akanuma, Hiroshi Sasaki, Loren D. Williams, and Claudia Alvarez-Carreno for fruitful discussions.

## SUPPLEMENTARY MATERIALS

### Materials and Methods

#### Identification of modern DPBB sequences with high internal symmetry

Sequences of the molecular chaperone VCP were identified from the in-house protein database with reduced taxonomic bias (*1*) by searching with the BlastP program (*2*) using the taVCP_DPBB sequence as a query, and only the DPBB domains were extracted from the full-length VCP proteins. The 249 sequences thus obtained were aligned with the Muscle program. We fragmented the aligned sequences at the loop between the N-half and C-half *ββαβ* elements, by referring to the structural information of taVCP_DPBB, and re-aligned them together. Based on the obtained multiple alignments, the internal sequence identities in each organism’s DPBB were evaluated and ranked, as shown in Table S3.

#### SC-design

Perfectly symmetric DPBBs were constructed by introducing mutations in a stepwise manner, using mkVCP_DPBB and apVCP_DPBB as templates. Initially, mkDPBB_sym_65 (internal sequence identity: 65%) and apDPBB_sym_63 (63%) were constructed by introducing the symmetrically-conserved residues in the N- and C-halves of mkVCP_DPBB and apVCP_DPBB into each other. The conserved residues in the other eight DPBBs with high internal sequence identities were then introduced into mkDPBB_sym_65 and apDPBB_sym_63 to construct mkDPBB_sym_79 (79%) and apDPBB_sym_79 (79%). The engineering step was repeated one more time using the sequence information from ten additional organisms, to construct mkDPBB_sym_86 (86%) and apDPBB_sym_84 (84%). mkDPBB_sym_86 and apDPBB_sym_84 have only a limited number of non-symmetrized positions, which are clustered in two areas in their tertiary structures. By adopting the amino acid residues from each of the two non-symmetrized areas, we then designed four DPBB proteins with perfect internal sequence symmetries (mkDPBB_sym1, mkDPBB_sym2, apDPBB_sym1, and apDPBB_sym2).

#### RE-design

The design of completely symmetrical DPBB sequences was carried out using the “reverse engineering evolution” computational protein design approach (*3*). This method starts from the pseudo-symmetric sequences of each subunit, then constructs a phylogenetic tree, and subsequently generates putative ancestral sequences for further evaluation. This has been successfully applied to the design of symmetric proteins with 3-8 subunits (*4, 5*). However, DPBB is an extreme case with only 2 subunits, which makes the prediction of ancestral sequences based on the phylogenetic tree constructed from only 2 sequences challenging. To overcome this challenge, we have introduced the use of orthologous sequences instead of pseudo-symmetric sequences for the construction of phylogenetic trees. We first collected 498 VCP sequences from different organisms and then removed the redundant sequences having >75% sequence identity. The resulting sequences were aligned to generate a phylogenetic tree using a maximum-likelihood method (Jones-Taylor-Thornton (JTT) model) with 50-100 bootstrapping (*6, 7*). The aligned sequences and constructed phylogenetic tree were used together to generate putative ancestral consensus sequences by the FastML server, using a joint reconstruction substitution model (*8*). Approximately 28,000 ancestral sequences were then mapped onto a manually constructed, symmetrical mkVCP backbone structural model, and their energies were calculated using an in-house “sequence_mapping.py” program that utilizes the PyRosetta of Rosetta protein modeling suite (*9–11*). This program uses an input list of plausible sequences and maps them onto a template protein backbone structure to output a PDB model for each sequence and the associated score. The top-scored designs were then analyzed for Rosetta total score, and root mean square deviations (RMSDs) from the symmetrical mkVCP backbone model structure. Finally, shortlisted designs were selected for experimental validation based on the Rosetta scores, RMSD from the design template, predicted solubility and visual inspection.

#### MS-design

##### 1) POMP^d^ (POsitive Multistate Protein design)

POMP^d^ was used to compute minimum energy protein sequences from an ensemble of conformational states (*12*). Let us assume a rigid backbone and a pairwise decomposable energy function taking the form:

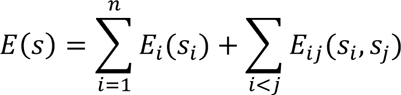

with *E*(*s*) the total energy of protein sequence *s* of length *n*, *E*_*i*_(*S*_*i*_) a unary energy term for residue *s*_*i*_ and *E*_*ij*_(*s*_*i*_, *s*_*j*_) a binary term representing energy interactions between residue pairs (*s*_*i*_, *s*_j_). In this context, POMP^d^ looks for the sequence that minimizes the energy of an ensemble of conformational states described as the sum of the energies on each state. POMP^d^ models this problem as a cost function network and solves it exactly, using the constraint programming prover toulbar2, by returning the global minimum of the energy function and proving its optimality. This deterministic approach provides optimality guarantees: given an ensemble of conformational states and an energy function, the sequence returned is the global minimum of the energy function.

##### 2) Conformational state ensemble preparation

mkVCP_sym2 crystal structure was used as the initial template. Two strategies were used in order to generate conformational states. In the first one, 100 protein models were generated using Rosetta Backrub protocol, with harmonic restraints on initial atoms coordinates. In the second one, a 100-ns molecular dynamics (MD) simulation at 300 K was performed starting from the initial template. Conformational states were extracted every 0.5 ns, generating 200 protein models. For each strategy, the resulting protein models were clustered using Durandal (*13*). The clustering radius was set to 0.15 Å for the first strategy and 0.5 Å for the second. In each case, the cluster centers of the 4 biggest clusters were selected as the ensemble of conformational states. From these two strategies, we obtained two ensembles of 4 conformational states: the backrub ensemble and the MD ensemble.

##### 3) Protein sequence predictions

POMP^d^ was used to compute optimal protein sequences from backrub and MD ensembles using two different setups, which allowed the prediction of a total of four protein sequences. For both setups, we used the ability of the toulbar2 prover to accept hard constraints in energy minimization to constrain sequences to be identical in each DPBB symmetrical subunit. In the first setup, all amino acid types were allowed at each position in a DPBB subunit. In the second setup, additional hard constraints were used in order to prevent the formation of solvent exposed hydrophobic patches.

While environmental factors such as pH, ionic strength, temperature, and the presence of various solvent additives may influence protein solubility, internal factors are defined by the amino acids present at the protein surface (*14*). Protein solubility is determined by the amount of exposed hydrophobic surface area in the protein folded state (*15, 16*). Furthermore, the rate of aggregation of proteins and peptides increases as the amount of exposed hydrophobic surface area increases (*17*). Therefore, computational protein design tools must take surface hydrophobicity into account when designing new sequences. In order to do so, we limit the formation of exposed hydrophobic surface area by adding constraints in the form of new energy function terms. This functionality has been implemented in POMP^d^ an additional feature called hpatch. The hpatch procedure is described in Supplementary Algorithm S1.

Forward folding experiments were performed on the four predicted sequences, using the protein structure prediction software EdaRose (*18*). Forward folding aims at assessing the quality of a protein design by predicting whether it will fold into the target structure or not. For each sequence, 30,000 structural models were predicted with EdaRose (Fig. S10). The number of iterations of EdaRose was set to 6, and the beta_nov16 scoring function from the Rosetta modeling software was used.

After examination of sequences and forward folding results, two sequences were selected for experimental characterization. The first one (msDPBB_sym1), from the backrub ensemble and using hpatch, was selected based on its forward folding profile. The second one (msDPBB_sym2), from the backrub ensemble and without using hpatch, was selected due to its sequence dissimilarity with other designs.

#### Molecular dynamics simulations

All simulations were performed using the Amber ff14SB force field (*19*) implemented in the AMBER 16 package (*20*). To obtain a neutral charge of the simulated systems, several counter- ions were included. Each protein with the counter-ions was solvated with TIP3P water molecules, using an octahedral box with a minimum distance of 12 Å between the solute and the simulation box edges. All systems were first subjected to 7 iterations of 1,000 minimization steps consisting of 500 steps of steepest descent minimization followed by 500 steps of conjugated gradient. A decreasing harmonic restraining potential was applied to the solute heavy atoms during the 6 first minimization iterations using a force constants of 100, 50, 20, 10, 5, and 1 kcal.mol^-1^.Å^-2^ respectively. Heating of each system (NVT simulation) up to 300 K was carried out during 100 ps under periodic boundary conditions, with positional restraints applied to the solute heavy atoms using a force constant of 25 kcal.mol^-1^.Å^-2^. NVT simulation at the target temperature (300 K) was further conducted during 300 ps in the same conditions. A 200 ps simulation at constant pressure and temperature (NPT) was later performed to equilibrate the pressure of each system around the target value of 1 bar. During this step, a weak positional restraint was applied on the solute heavy atoms, using a force constant of 5 kcal.mol^-1^.Å^-2^. An unrestrained MD simulation of 100 ns was finally performed under the same conditions. The Berendsen algorithm (*21*) was used to keep both temperature and pressure constant during simulations. A cutoff of 9 Å was used to define long-range electrostatic interactions, which were calculated using the Particle Mesh Ewald algorithm (*22*). Bonds involving hydrogen atoms were constrained using the SHAKE algorithm (*23*) to enable the use of a 2 fs time step. All trajectory analyses were carried out with the CPPTRAJ module (*24*).

#### Construction of expression vectors

The synthetic genes encoding the proteins used in this study were purchased (Thermo Fisher Scientific, MA, USA). After amplifying the genes by PCR, each product DNA fragment was cloned using In-Fusion^TM^ (TAKARA Bio, Japan) into the pET47b vector, to add a cleavable N- terminal His6-tag to the sequences. When sub-cloning the genes for a halved fragment, “XXX_half” and “Cloning_upstream” primers were used in the PCR amplification. The DNA sequences used in this study are listed in Supplementary Table S11.

#### Protein expression and purification

To produce the proteins, competent *E. coli* BL21-Gold(DE3) cells (Agilent Technologies, CA) were transformed with the respective expression vectors. The transformants were cultured at 37°C overnight in 20 mL of Luria Broth medium supplemented with 20 µg/mL kanamycin. The cells were then inoculated into 2 L of Luria Broth medium and cultured at 37°C for 2 hours. For induction, 0.5 mM isopropyl *β*-D-1-thiogalactopyranoside (IPTG) was added to the media, and the desired proteins were expressed for 4 hours under the same conditions. After harvesting the cells, the pellets were resuspended in 60 mL of 50 mM potassium phosphate buffer, pH 6.5, 150 mM NaCl, and sonicated. The bacterial lysate was fractionated into supernatant and precipitant by centrifugation at 8,000 rpm and 4 °C, for 20 min. To precipitate the contaminating *E. coli* proteins, the supernatants were incubated at 70 °C for 20 min, and then each soluble fraction was isolated by centrifugation at 8,000 rpm and 4 °C for 20 min. In the cases of mk2h_ΔMILPS, mk2h_ΔMILYS, mk2h_ΔMILPY, and mk2h_ΔMILPYS, the above heat treatment process was omitted in order to preserve their native structures. The soluble proteins were purified by HisTrap HP nickel affinity chromatography (GE Healthcare, IL). The N-terminal His6-tags were cleaved with HRV-3c protease (Funakoshi, Japan) at 4°C for 1–2 days. To remove the cleaved His6-tag and residual uncleaved proteins, the treated solutions were loaded onto HisTrap columns, and each flow-through fraction was recovered. The protein solutions were additionally loaded onto a HiLoad 16/600 Superdex75 (GE Healthcare, IL) size exclusion chromatography column, equilibrated with 50 mM potassium phosphate buffer, pH 6.0, 150 mM NaCl, and the peak fractions were collected. The purity of each protein was verified by SDS-PAGE, and the protein concentrations were determined using their A280 values, measured with an ultraviolet spectrophotometer (NanoDrop, Thermo Scientific).

For the preparation of seleno-methionine (Se-Met) substituted taVCP_DPBB, *E. coli* BL21- Gold(DE3) cells were grown in 2 L of M9 minimal medium containing 20 μg/L of kanamycin at 37 °C until they reached an absorbance at 600 nm (*A*600) of 0.4. An amino acid mixture (50 mg/L isoleucine, leucine and valine and 100 mg/l phenylalanine, threonine and lysine) and seleno-L-methionine (60 mg/mL) were then added to the culture, and the cells were grown at 37°C. After reaching an *A*600 of 0.8, the protein expression was induced with 0.5 mM IPTG, and the cells were grown further at 37°C for 5 hours.

#### Biophysical characterization

For the gel filtration analysis, the concentrations of full-length proteins and halved fragments were adjusted to 20 µM and 40 µM, respectively. A 100 µl aliquot of each purified protein was applied to a Superdex 75 Increase 10/300 size exclusion chromatography column, equilibrated with 50 mM potassium phosphate buffer, pH 6.0, 150 mM NaCl, and run on an AKTA FPLC (Amersham Biosciences) at a flow rate of 0.75 mL/min.

CD spectra were collected on a JASCO J820 circular dichroism spectrometer (JASCO, Japan). Samples containing 20 μM full-length proteins or 40 µM halved fragments, in 50 mM potassium phosphate, pH 6.0, 150 mM NaCl, were loaded into a 1 mm pathlength quartz cuvette. Spectra were recorded in the wavelength range from 200 to 250 nm at 1 nm intervals at 25°C, and each spectrum was the average of 10 scans. Spectra for mk2h mutants containing 7, 8, or 10 amino acid repertoires were recorded at 10 °C.

The melting curves were collected on a JASCO J820 CD spectrometer monitored at 222 nm, in 50 mM potassium phosphate buffer, pH 6.0, 150 mM NaCl. The temperature was increased at a rate of 1.0 °C/min. Data points were collected at 0.1 °C increments from 35 °C to 95 °C, or from 10 °C to 95 °C.

#### Crystallography

For crystallization, all purified protein solutions were dialyzed against 20 mM Bis-tris HCl, pH 6.0, 150 mM NaCl and then concentrated to 30–70 mg/ml. Crystallization screenings were carried out in 96-well sitting-drop vapor-diffusion plates. Sample solutions (200 nL) were mixed with an equal amount of reservoir solutions and incubated at 20°C. Almost all crystals were obtained after a few hours to a few days. The crystals were cryo-cooled using the reservoir solution including 13–30% glycerol as a cryo-protectant (Table S12).

Data were collected at the Photon Factory (Tsukuba, Japan)(*25, 26*), SPring 8 (Harima, Japan)(*27–32*), or Swiss Light Source (Villigen, Swiss). The beam lines and detectors are listed in Table S9. The X-ray diffraction data were processed with XDS (*33*). All structures were solved and refined with the program PHENIX (*34, 35*). The structures of taVCP_DPBB, mkVCP_DPBB, and apVCP_DPBB were solved by the SAD phasing method, using phenix.autosol. The initial structure models for other mutants were determined by the MR phasing method, using phenix.phaser-MR and the crystal structures of DPBB determined in this study as the search models. The model structures were updated manually using Coot (*36*) and iteratively refined with Phenix.refine. Statistics for diffraction data collection and refinement are summarized in Table S2.

#### Characterization of chemically-synthesized peptides

All chemically-synthesized peptides tested in this report were obtained from Japan Bio Services. The synthesized mk2h (95.14% purity) and mk2h_ΔMILPS (95.56%) peptides in powdered forms were dissolved in 20 mM Bis-tris HCl, pH 6.0, 150 mM NaCl. The concentrations were determined by the absorbance at 280 nm, using an ultraviolet spectrophotometer (NanoDrop One, Thermo Scientific). For the crystallization screenings, the mk2h_ΔMILPYS peptide (95.85%) was dissolved in 20 mM Tris-HCl, pH 8.5, 200 mM lithium sulfate. The dissolved peptide sample was separated into supernatant and precipitate fractions, and then the precipitate was resuspended in the same buffer. Both the supernatant and undissolved suspension were used for the crystallization screenings, which were carried out in 96-well sitting-drop vapor-diffusion plates. The obtained crystals were processed in the same way as the bacteria-produced proteins described above.

To examine the foldability in the presence of the contaminated sequences, the low purity peptides of mk1h (75.06%) and mk2h (71.16%) were dissolved in 20 mM Bis-tris HCl, pH 6.0, 150 mM NaCl. The aggregated and dimer species were separated on a size exclusion chromatography column (Superdex 75 Increase 10/300) equilibrated with 20 mM Bis-tris HCl, pH 6.0, 150 mM NaCl. The peptide concentrations of three samples, 1) before SEC purification, 2) aggregated fraction, and 3) dimer fraction, were adjusted to 20 µM and 10 µM for CD spectrum measurements and liquid chromatography mass spectrometry, respectively. Using a JASCO J820 circular dichroism spectrometer (JASCO, Japan), CD spectra were recorded at wavelengths from 200 to 250 nm at 1 nm intervals at 25 °C, and each spectrum was the average of 10 scans.

#### LC-MS analysis

The peptide solutions were desalted with in-house made C18 stage-tips, dried under a vacuum, and dissolved in 2% acetonitrile and 0.1% formic acid. The 10 pmol peptide mixtures were fractionated by C18 reverse-phase chromatography (1.8 μm, ID 0.075 mm x 250 mm, Aurora UHPLC Column; IonOpticks, ADVANCE UHPLC; AMR Inc.) and applied directly into a hybrid linear ion trap mass spectrometer (LTQ Orbitrap Velos Pro; Thermo Fisher Scientific). The peptides were eluted at a flow rate of 200 nL/min with a linear gradient of 5–35% solvent B over 20 min. The compositions of Solvent A and B were 0.1% TFA in water and 100% acetonitrile, respectively.

The Orbitrap mass spectrometer was programmed to carry out 4 successive scans, with the first consisting of a full MS scan from 350–2,000 m/z at a resolution of 60,000, and the second to fourth consisting of data-dependent scans of the top three most abundant ions obtained in the first scan, at a resolution of 7,500. Automatic MS/MS spectra were obtained from the highest peak in each scan, by setting the relative collision energy to 35% and the exclusion time to 90 s for molecules in the same m/z value range. Calculations of peptide masses and intensities in the time range from 15.0–35.0 min were performed with the Xtract deconvolution algorithm in FreeStyle, version 1.5 (Thermo Fisher Scientific).

### Supplementary Text

#### SC-design

Most of the amino acid residues conserved between the N- and C-halves in each VCP_DPBB have probably remained unchanged from their perfectly symmetric ancestor, as the chance of having the same amino acid residues in the symmetric positions by random mutation is low. This idea is supported by the observation that the symmetrically-conserved residues in the DPBB domain from each archaeon are often shared with other species (Fig. S4). Studies have also shown that *M. kandleri*, the organism possessing one of the highest symmetric extant DPBBs, is close to the phylogenetic root of archaea (*37–39*).

We used mkVCP_DPBB and apVCP_DPBB, with the highest symmetrical sequence identities among the DPBB domains in our database, as the starting templates to create perfectly symmetrical DPBBs. Initially, the symmetrically-conserved residues in mkVCP_DPBB or apVCP_DPBB were introduced into each other to construct the chimeric DPBBs, mkDPBB_sym_67 (internal sequence identity: 67%) and apDPBB_sym_63 (63%) (Table S1). We confirmed that both proteins were monomers with an *α/β* structure (Fig. S5A and C). We then introduced the symmetrically- conserved residues in the other eight VCP_DPBB sequences top-ranked by internal sequence identities (Fig. S4 and Table S3) into mkDPBB_sym_67 and apDPBB_sym_63, resulting in mkDPBB_sym_81 (81%) and apDPBB_sym_79 (79%). Additionally, the symmetrically- conserved residues in ten more VCP_DPBB sequences were introduced, to construct mkDPBB_sym_86 (86%) and apDPBB_sym_84 (84%). Three proteins, except for mkDPBB_sym_81, were verified to be folded (Fig. S5B, D, and E). X-ray-crystallography confirmed that mkDPBB_sym_86 and apDPBB_sym_79 adopt the DPBB fold (Figs. 1E, S6A). The mkDPBB_sym_86 and apDPBB_sym_84 proteins have only a limited number of non-symmetrized positions, which are clustered in two areas. In mkDPBB_sym_86, cluster-1 comprises 4L, 7K, 8L, 28A, 29S, and 64G around *α*-helix 1 and *β*-strand 1’, and cluster-2 comprises 21R, 47K, 50V, 51A, 71Y, and 72E around *α*-helix 1’ and *β*-strand 1 (Fig. 1E). Looking at the overall structure of the original full-length VCP containing the other domains, these symmetrical faces in the DPBB domain are in different environments: the residues at cluster-1 are exposed to the solvent, while the residues at cluster-2 contact another domain (Fig. S7). This difference in the molecular environments probably led to a breakdown of the symmetry in this area during the evolutionary process.

Subsequently, we designed four DPBB domains with perfect internal sequence symmetries (mkDPBB_sym1, mkDPBB_sym2, apDPBB_sym1, and apDPBB_sym2) by adopting the amino acid residues from either cluster-1 or -2. While apDPBB_sym2 could not be purified due to its poor stability, mkDPBB_sym1, mkDPBB_sym2, and apDPBB_sym1 were purified as stable proteins (Fig. S5F–H). We determined the crystal structures of mkDPBB_sym1 and mkDPBB_sym2 to confirm that they adopt the DPBB fold as designed (Fig. 1F, Table S2). Therefore, we succeeded in reconstructing the perfectly symmetrical DPBB structures, using the sequence information of VCP-like chaperones from only twenty archaeal species. This result led us to anticipate that the archaeal common ancestor possessed a nearly perfect symmetric DPBB sequence in the VCP-like chaperone gene. Furthermore, all of the perfectly symmetrical DPBB and intermediate mutants constructed by the SC-design exhibited high stability. Except for mkDPBB_sym_67 (*Tm*=69.2 °C), they did not completely unfold even at 95 °C (Fig. S5). These results support the hypothesis that the common ancestor of archaea is a thermophile (*40*).

#### RE-design

We utilized a modified “reverse engineering evolution” protein design approach to design the symmetrical DPBB sequences. In this design methodology, orthologous sequences of the target protein are used instead of pseudo-symmetric sequences for the construction of phylogenetic trees. Specifically, we used VCP_DPBB sequences from different organisms to construct a phylogenetic tree with respective aligned sequences, which were subsequently used as input to generate ancestral sequences (Figure S8A and B). The predicted ancestral sequences were mapped onto a manually constructed, perfectly symmetrical mkVCP_DPBB structural backbone model using an in-house program that utilizes PyRosetta, and each sequence was ranked by the Rosetta score (Figure S8C and D). The top-scored designs were analyzed for Rosetta total score, RMSD from the manually generated, symmetrical mkVCP_DPBB model and through visual inspection. First, our analysis of the Rosetta total score revealed that many output models showed significantly lower energy and converged well (Figure S9A). When the backbone RMSDs of the designs were computed and plotted against the Rosetta total score, a broad spread of total score/RMSD scores was obtained. However, the majority of the top-scored output models exhibited RMSDs <1 Å, suggesting even with diverse sequences, they did not deviate much from the starting structure (Figure S9B). Interestingly, several of the top-scored models exhibited RMSDs <0.7 Å. The reDPBB_sym1 and reDPBB_sym2 exhibited 0.65 Å and 0.64 Å RMSD from the manually generated symmetrical mkVCP_DPBB model, respectively. Next, an analysis of Rosetta total score versus percentage sequence identity revealed that the top-scored structures tend to have higher percentage sequence identity (Figure S9C). However, the shortlisted reDPBB_sym1 and reDPBB_sym2 designs shared a moderate 78% and 74% sequence identity with the manually generated, symmetrical mkVCP_DPBB model respectively.

We compared the Rosetta generated structural models to that of the crystal structures of reDPBB_sym1 and reDPBB_sym2. First, we found that in both reDPBB_sym1 and reDPBB_sym2 crystal structures, each half of the proteins exhibit 0.41 Å RMSD with each other (Figures S9D–G). Moreover, while the crystal structure of reDPBB_sym1 and the Rosetta generated model of reDPBB_sym1 share 0.66 Å RMSD, the reDPBB_sym2 crystal structure and Rosetta generated model of reDPBB_sym2 share 0.58 Å RMSD (Figures S9H and S9I). This indicated that the Rosetta- generated structural models are in close agreement with the crystal structures. This result demonstrated that our computational symmetric design approach can be successfully applied to the design of DPBB fold as well.

Our designed reDPBB_sym1 and reDPBB_sym2 sequences share 67% and 65% sequence identity with mkVCP_DPBB, and 44% identity with taVCP_DPBB respectively. This further confirms that diverse sequences can fold into DPBB structures, as predicted from our computational symmetric protein design approach. Our computational strategy, of using orthologous sequences to construct phylogenetic trees for the design of symmetric proteins, has advantages over that of using pseudo-symmetric sequences from each subunit of the target protein. It circumvents the challenges when the number of subunits in the target protein is very small, such as the two subunits in DPBB. The successful computational design of symmetric DPBB proteins has verified the applicability of our design strategy under this circumstance.

---

**Figure S1.**
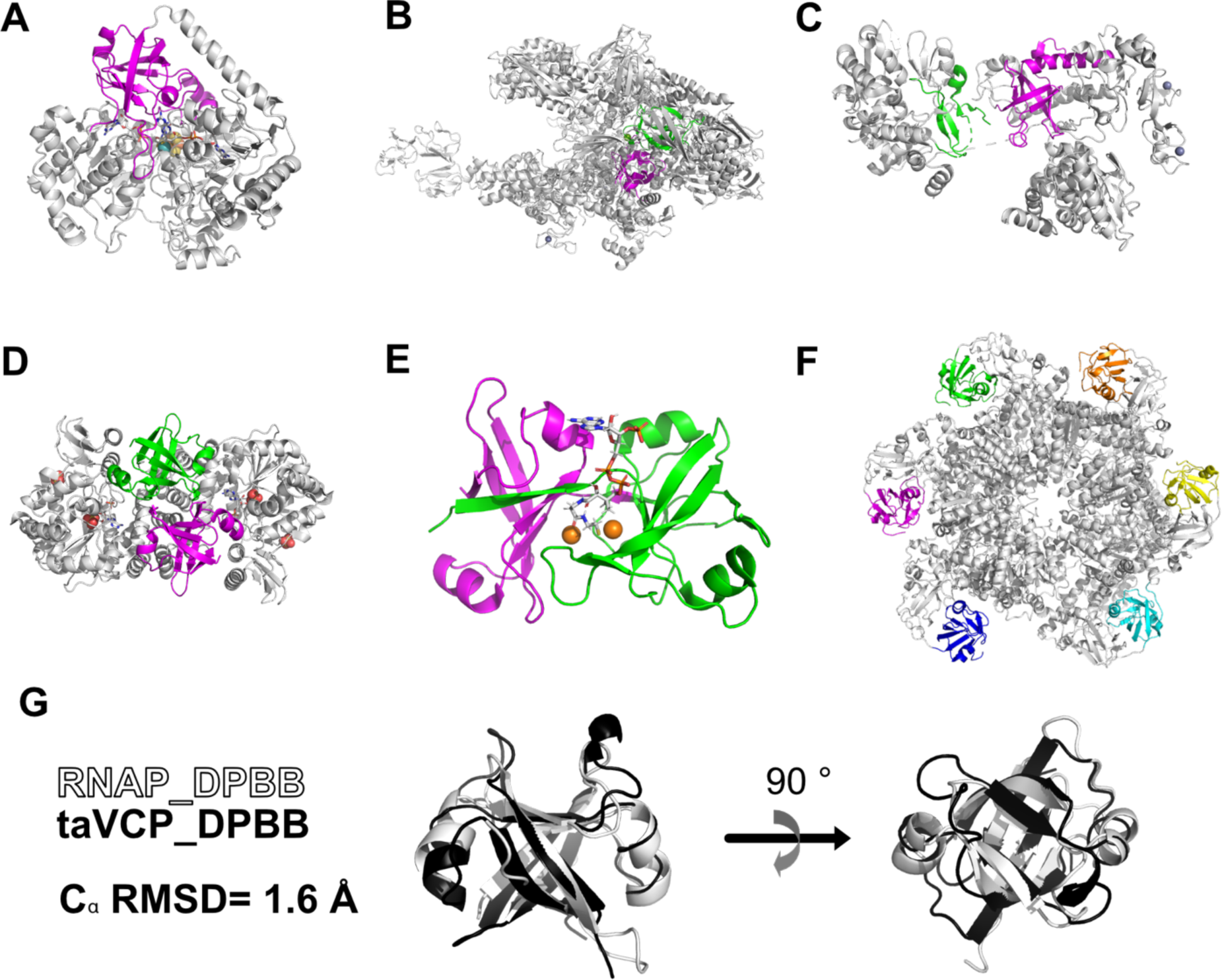
DPBB domains in natural proteins. Structures of (A) formate dehydrogenase (PDB ID 1FDO), (B) RNA polymerase (PDB ID 6ASG), (C) D-type DNA polymerase (PDB ID 5IJL), (D) AMP phosphorylase (PDB ID 4GA6), (E) phosphate propanoyltransferase (PDB ID 5CUO), and (F) molecular chaperone VCP (PDB ID 5G4F) are shown. Each DPBB domain is colored differently. (G) The superimposed structures of the DPBB domain-2 from RNA polymerase (colored pink in Fig. S1B) and the isolated DPBB domain of molecular chaperone VCP (Fig. 1B). The C*α* RMSD was calculated by CRICK (*41, 42*) (http://cospi.iiserpune.ac.in/click/).

**Figure S2.**
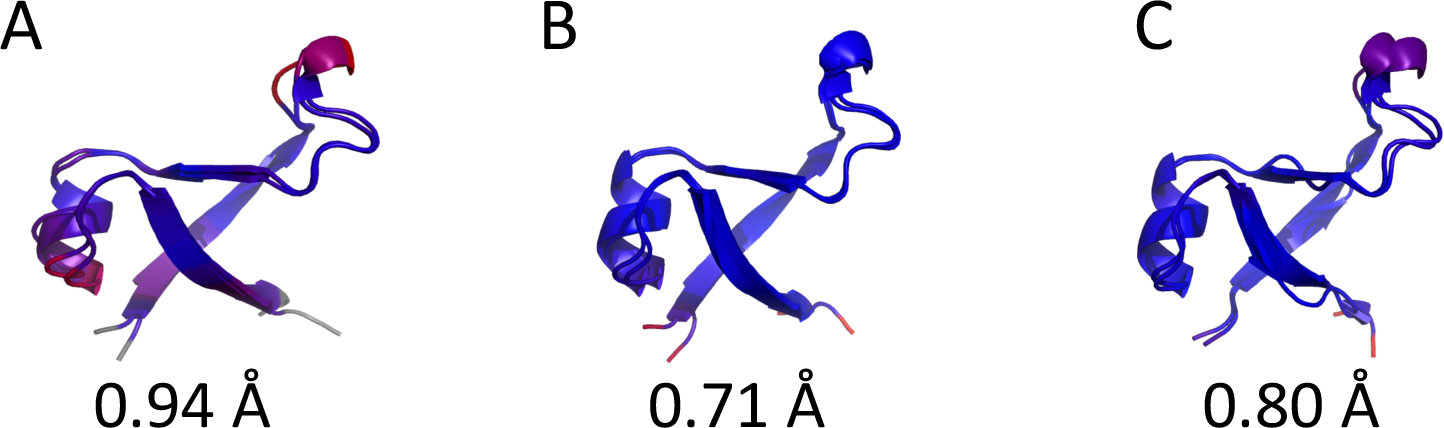
Superimposed structures of the N- and C-terminal halves of extant VCP_DPBBs. Superimposition of the N- and C-terminal halves of the crystal structures of (A) taVCP_DPBB, (B) mkVCP_DPBB, and (C) apVCP_DPBB. Darker blue indicates better alignments in the structures. The values are the C*α* RMSDs calculated by CRICK (*41, 42*) (http://cospi.iiserpune.ac.in/click/).

**Figure S3.**
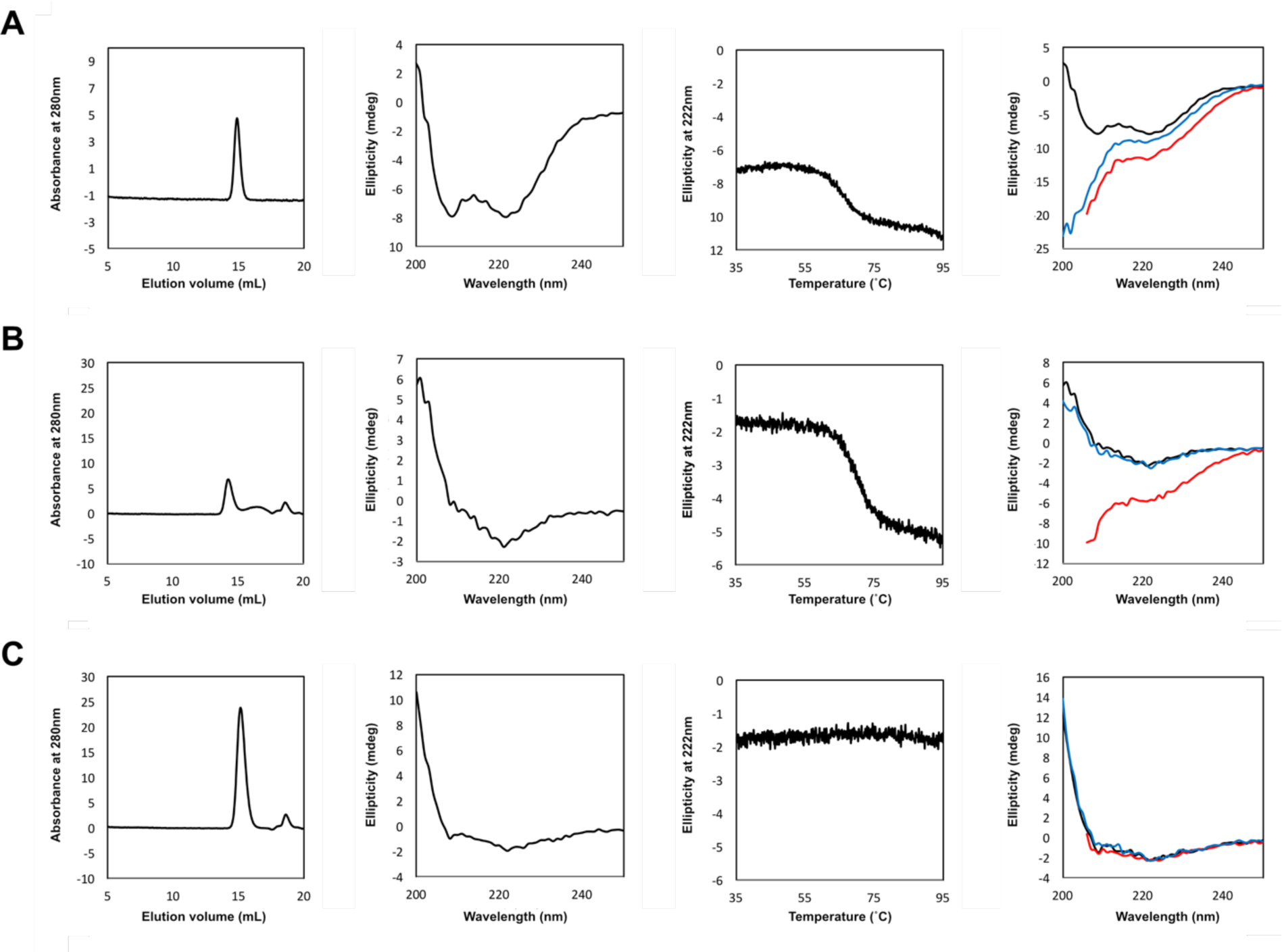
Experimental characterization of extant VCP_DPBBs. Size exclusion chromatography, CD spectra, denaturation curves, and comparisons of CD spectra at different temperatures (black: 35°C; red: 95°C; blue (refolding): 95°C à 35°C) for (A) taVCP_DPBB, (B) mkVCP_DPBB, and (C) apVCP_DPBB are shown in the panels from left to right.

**Figure S4.**
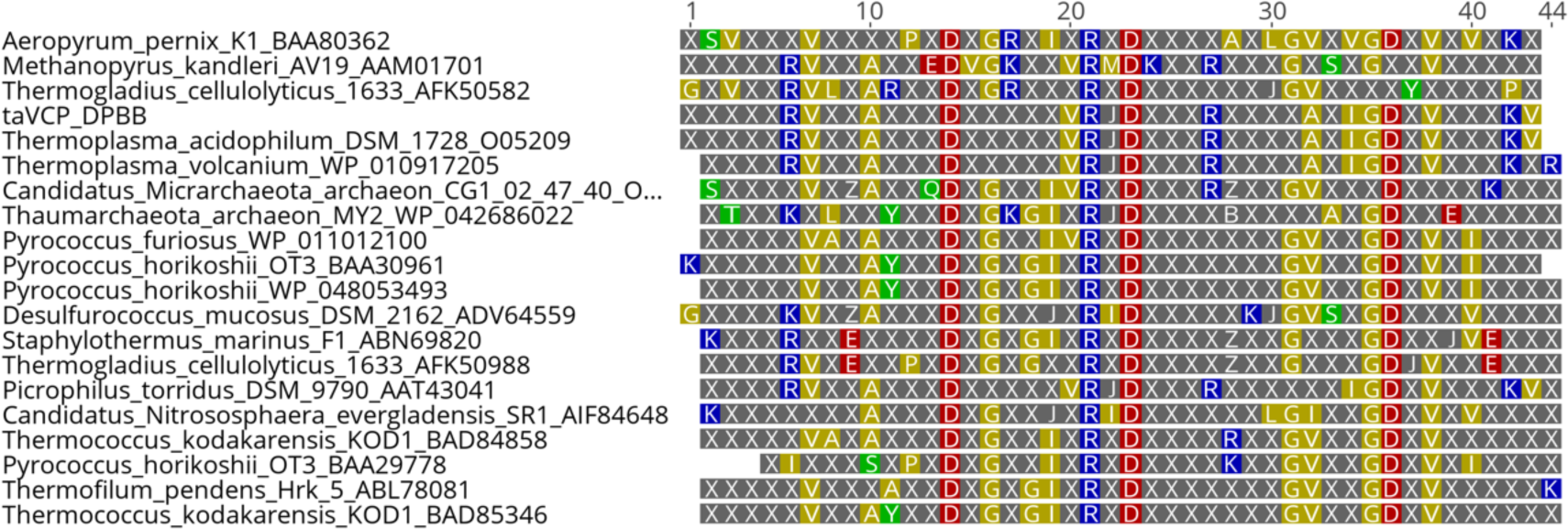
The symmetrically-conserved residues in natural VCP_DPBB domains. The symmetrically-conserved residues between the N- and C-halves in each 20 top-ranked VCP_DPBB with high internal sequence identity are highlighted. The non-symmetrically- conserved residues are represented as X (gray). The sequences are ordered by the internal sequence identity shown in Table S9. To create mkDPBB_sym_81 and apDPBB_sym_79, the symmetrically-conserved residues in the top ten sequences were considered. To create mkDPBB_sym_86 and apDPBB_sym_84, the symmetrically-conserved residues of all sequences were considered.

**Figure S5.**
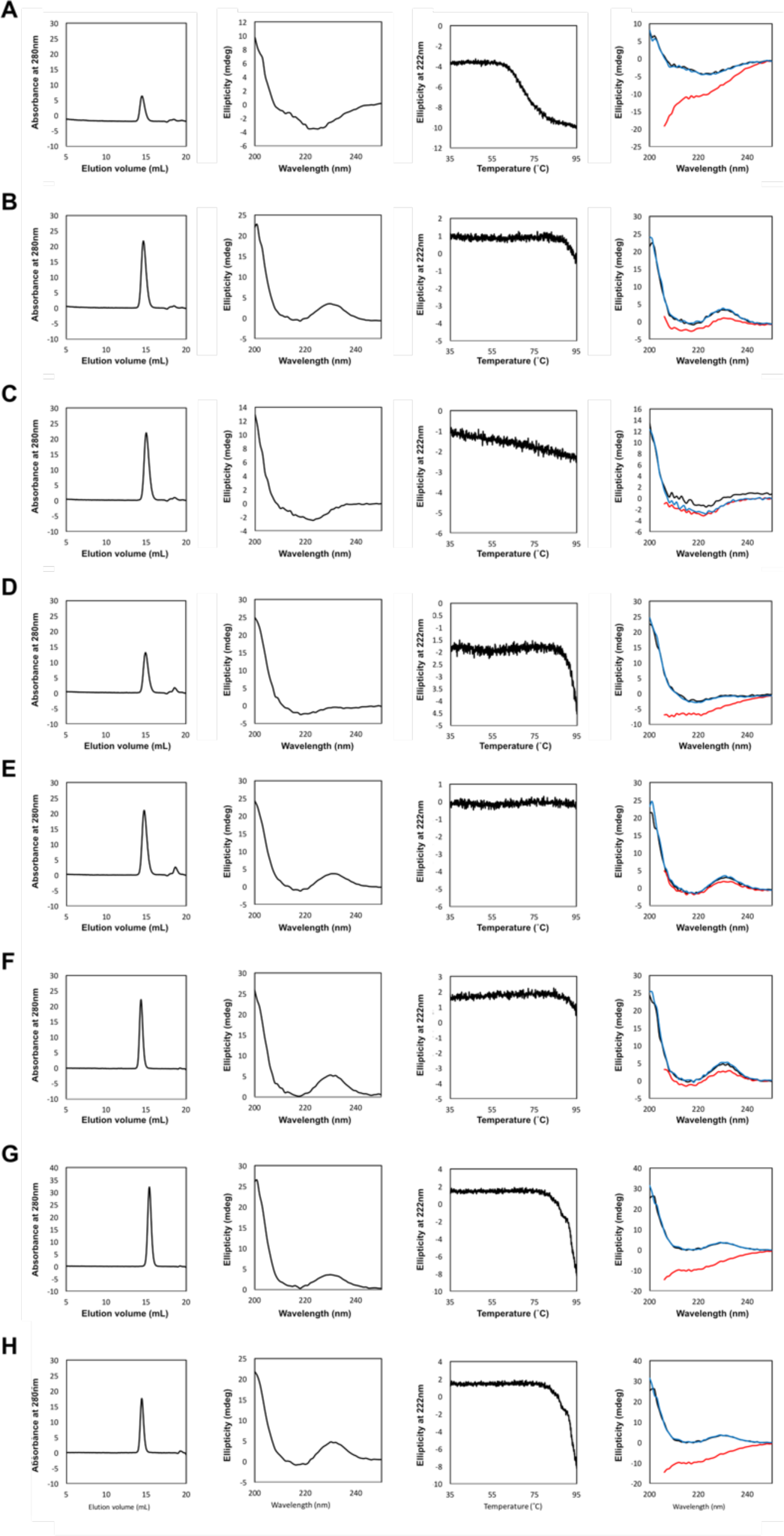
Experimental characterization of symmetrical DPBBs constructed by the SC- design method. Size exclusion chromatography, CD spectra, denaturation curves, and comparisons of CD spectra at different temperatures (black: 35°C; red: 95°C; blue (refolding): 95°C à 35°C) for (A) mkDPBB_sym_67, (B) mkDPBB_sym_86, (C) apDPBB_sym_63, (D) apDPBB_sym79, (E) apDPBB_sym_84, (F) mkDPBB_sym1, (G) mkDPBB_sym2, and (H) apDPBB_sym1 are shown in the panels from left to right.

**Figure S6.**
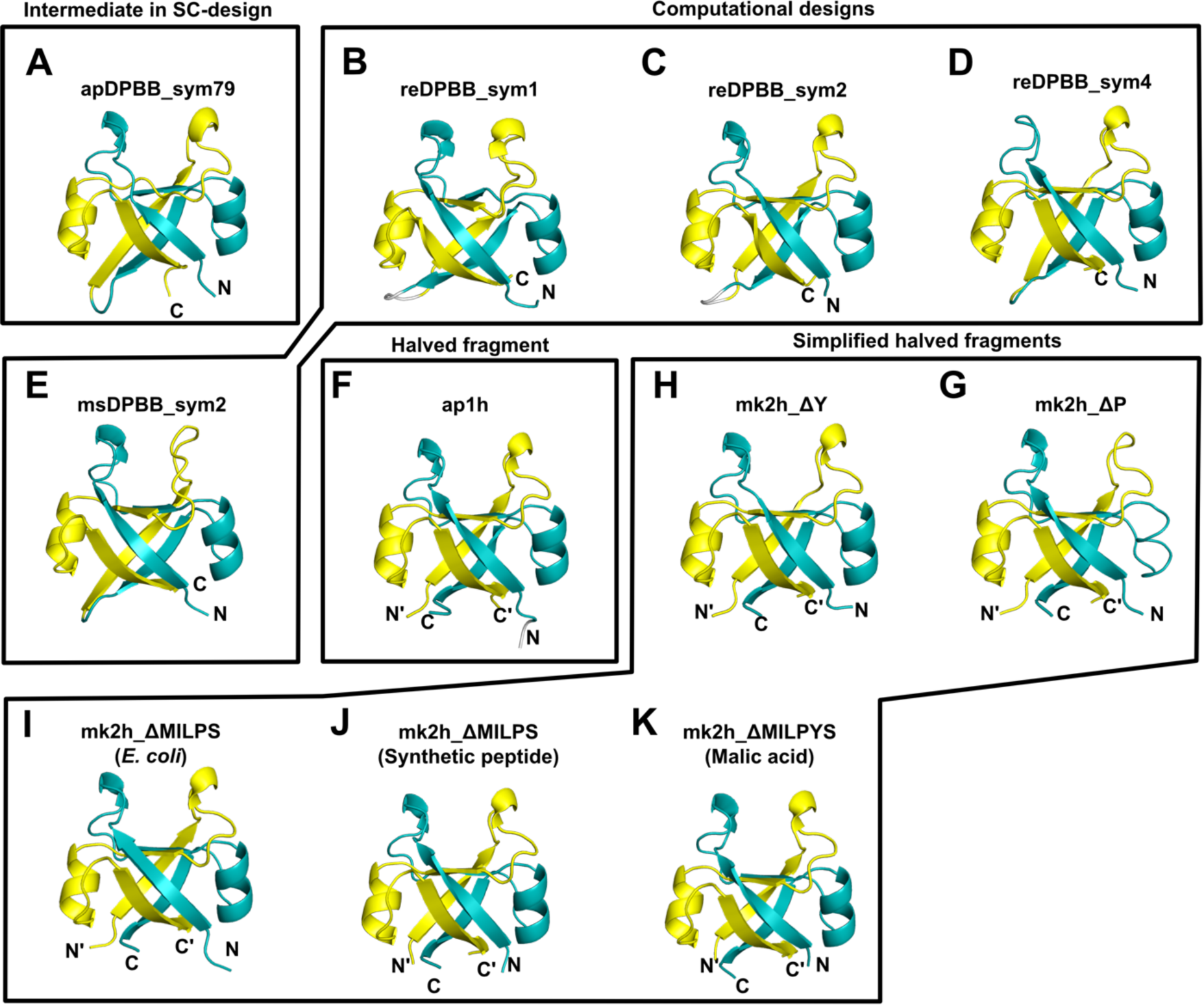
Crystal structures of the designed DPBB domains. (A) apDPBB_sym_79 (PDB ID 7DI0), (B) reDPBB_sym1 (7DVC), (C) reDPBB_sym2 (7DVF), (D), reDPBB_sym4 (7DVH), (E) msDPBB_sym2 (7DWW), (F) ap1h (7DXS), (G) mk2h_ΔP (7DXU), (H) mk2h_ΔY (7DXV), (I) *E. coli*-produced mk2h_ΔMILPS (7DXX), (J) chemically-synthesized mk2h_ΔMILPS (7DXY), and (K) chemically-synthesized mk2h_ΔMILPYS in the presence of DL-malic acid (7DYC).

**Figure S7.**
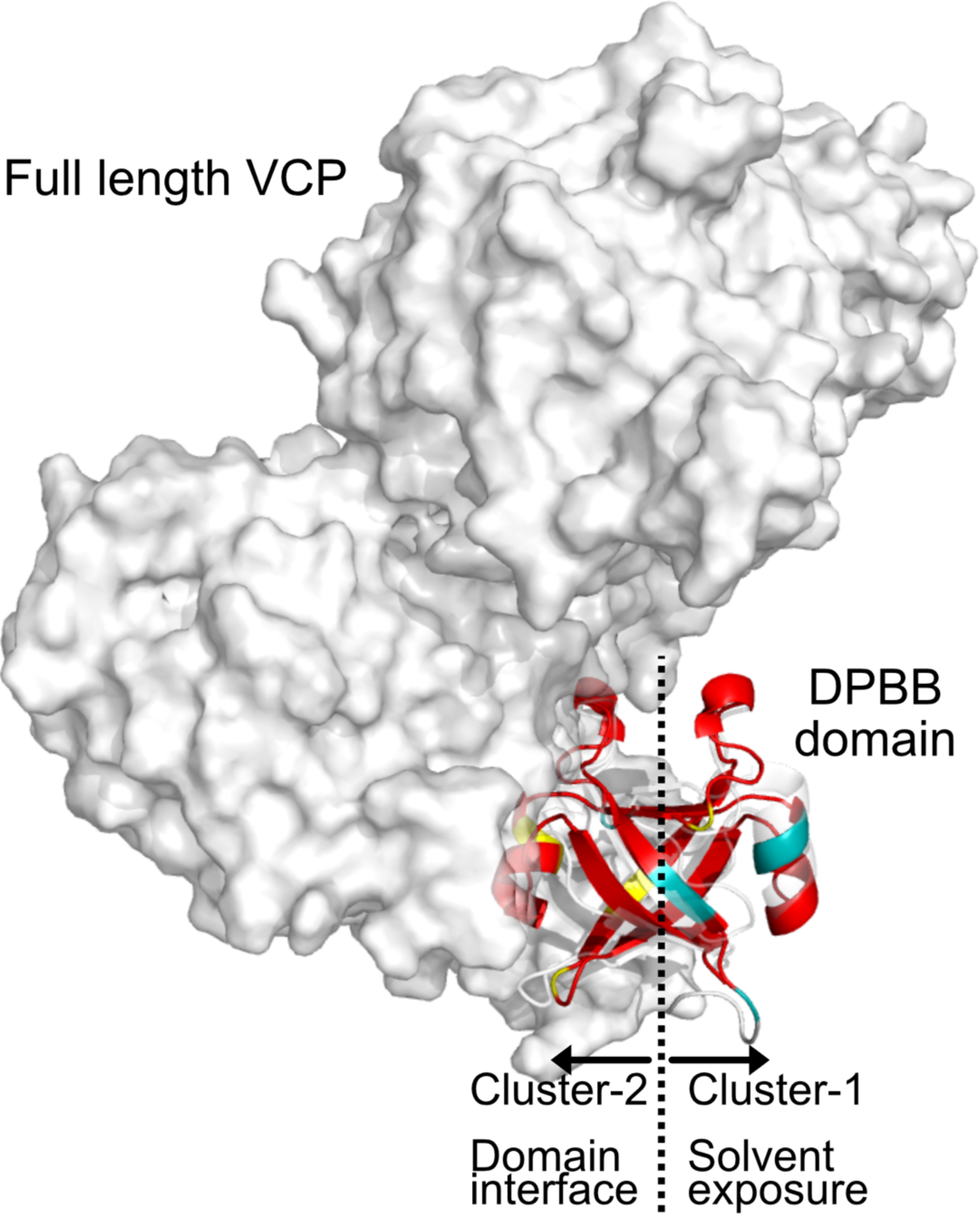
Different environments of symmetrical faces in the full-length VCP protein. The DPBB domain and other domains in the full-length VCP from *Thermoplasma acidophilum* (PDBID 5G4F) are represented by the white cartoon model and surface model, respectively. The crystal structure of mkDPBB_84 is superimposed with the DPBB domain and colored as in Fig. 1E. Cluster-1 is exposed to the solvent and cluster-2 is an interface to the other domain.

**Figure S8.**
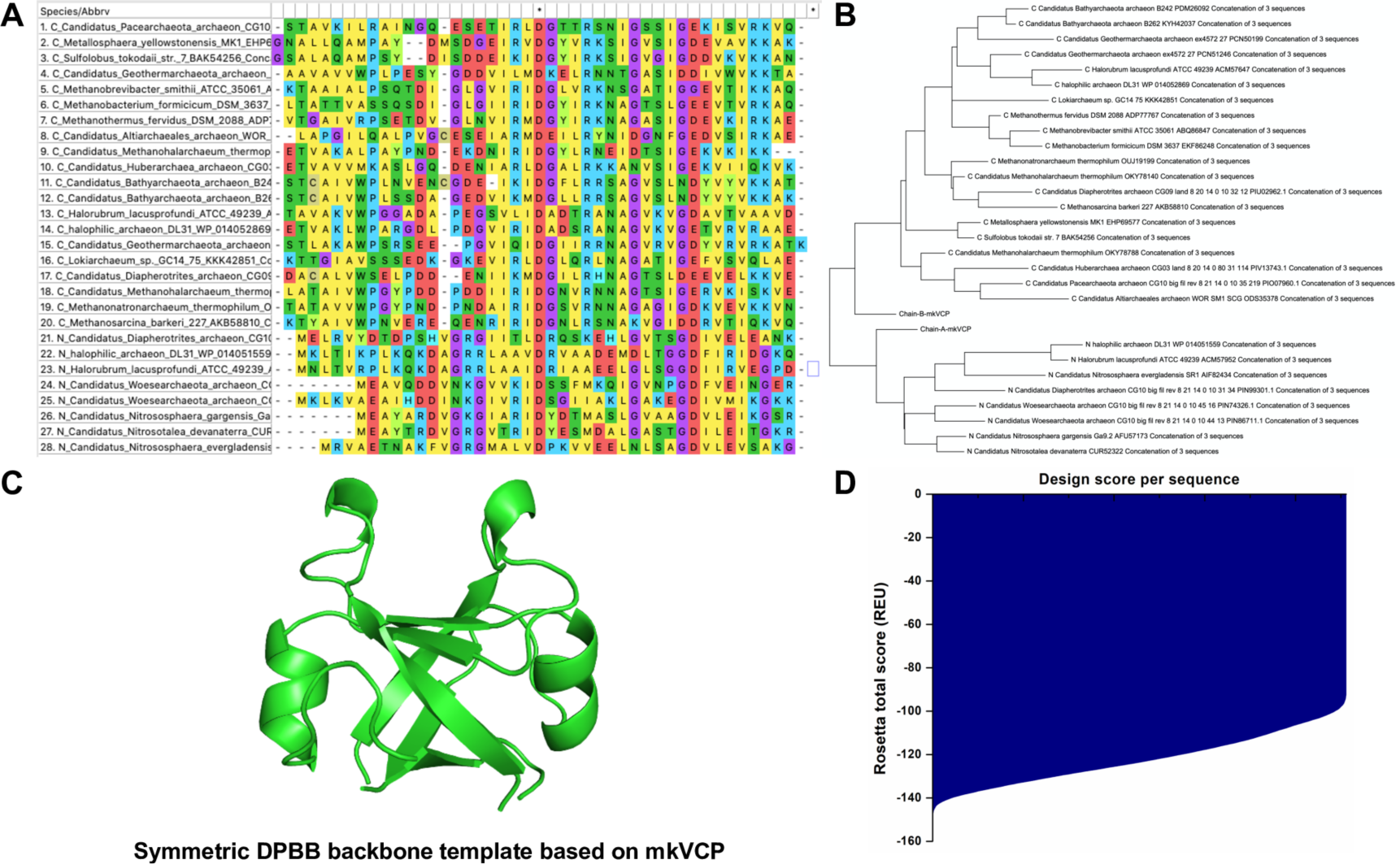
Reverse engineering evolution strategy to design a fully symmetric DPBB domain. (A) DPBB sequences from different organisms were aligned (B) to produce a phylogenetic tree, for use as input to generate possible ancestral sequences. (C) The ancestral sequences were mapped onto the symmetric DPBB backbone template based on mkVCP_DPBB and (D) scored using pyRosetta.

**Figure S9.**
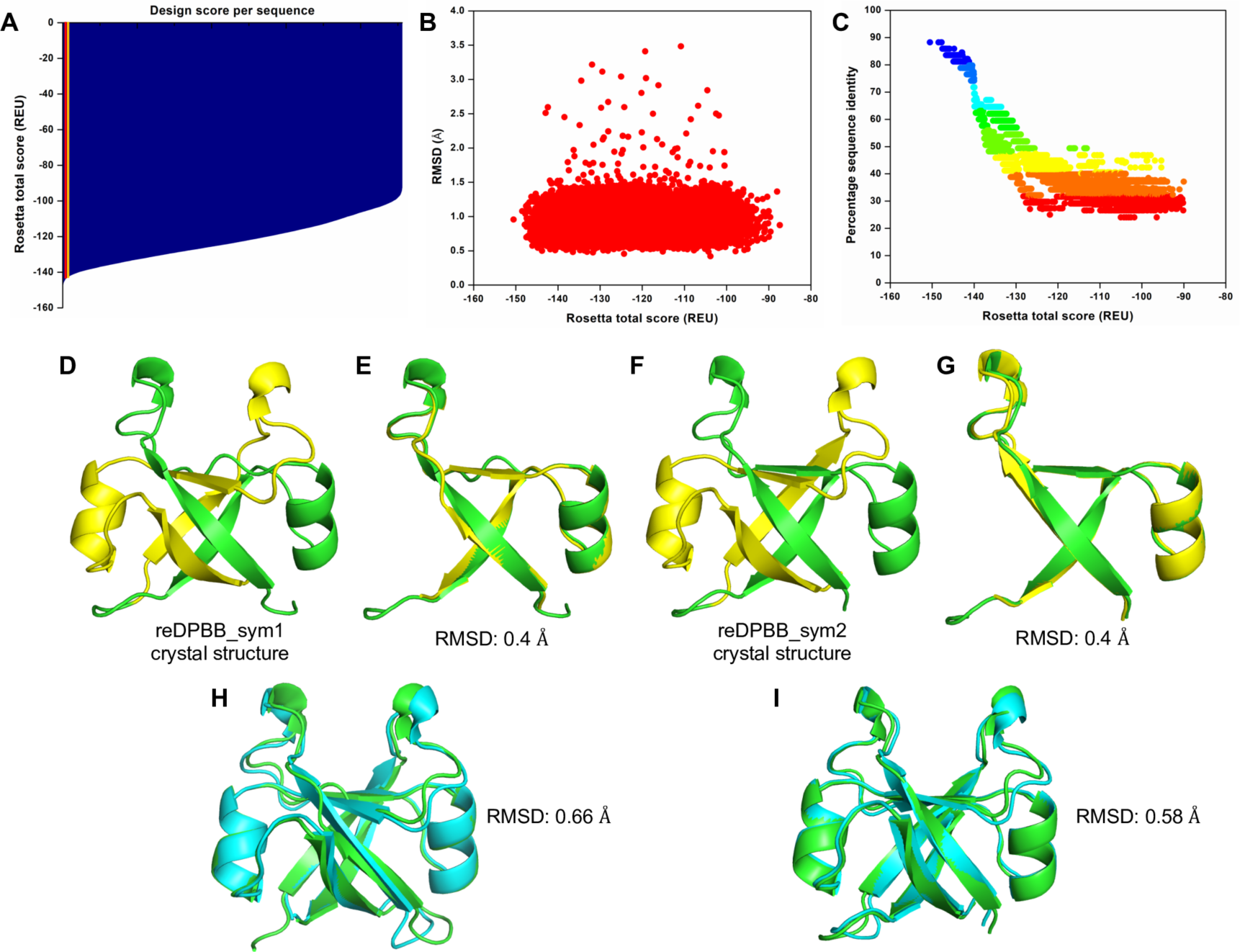
Structural and physicochemical parameters from Rosetta-design calculations, and structural differences between crystal structures and computationally modeled structures of symmetrical DPBBs. (A) Rosetta computed design score per sequence after the ancestral sequences were mapped onto the fully symmetrical DPBB structural model, where the red and yellow lines represent the top-scored reDPBB_sym1 and reDPBB_sym2 designs, respectively. (B) Rosetta scores versus the RMSD of the designs showing well-spread RMSDs from the template. (C) Rosetta total score versus the percentage sequence identity of the designs, which had sequence identities ranging from 25-90%. (D) Crystal structure of reDPBB_sym1. (E) Superimposition and computed RMSD between each half-barrel. (F) Crystal structure of reDPBB_sym2. (G) Superimposition and computed RMSD of each half-barrel, shown in green and blue colors, respectively. Structural superimpositions of (H) the reDPBB_sym1 crystal structure (green) with the Rosetta generated model (cyan) and (I) the reDPBB_sym2 crystal structure (green) with the Rosetta generated model (cyan), with RMSDs of 0.66 Å and 0.58 Å, respectively.

**Figure S10.**
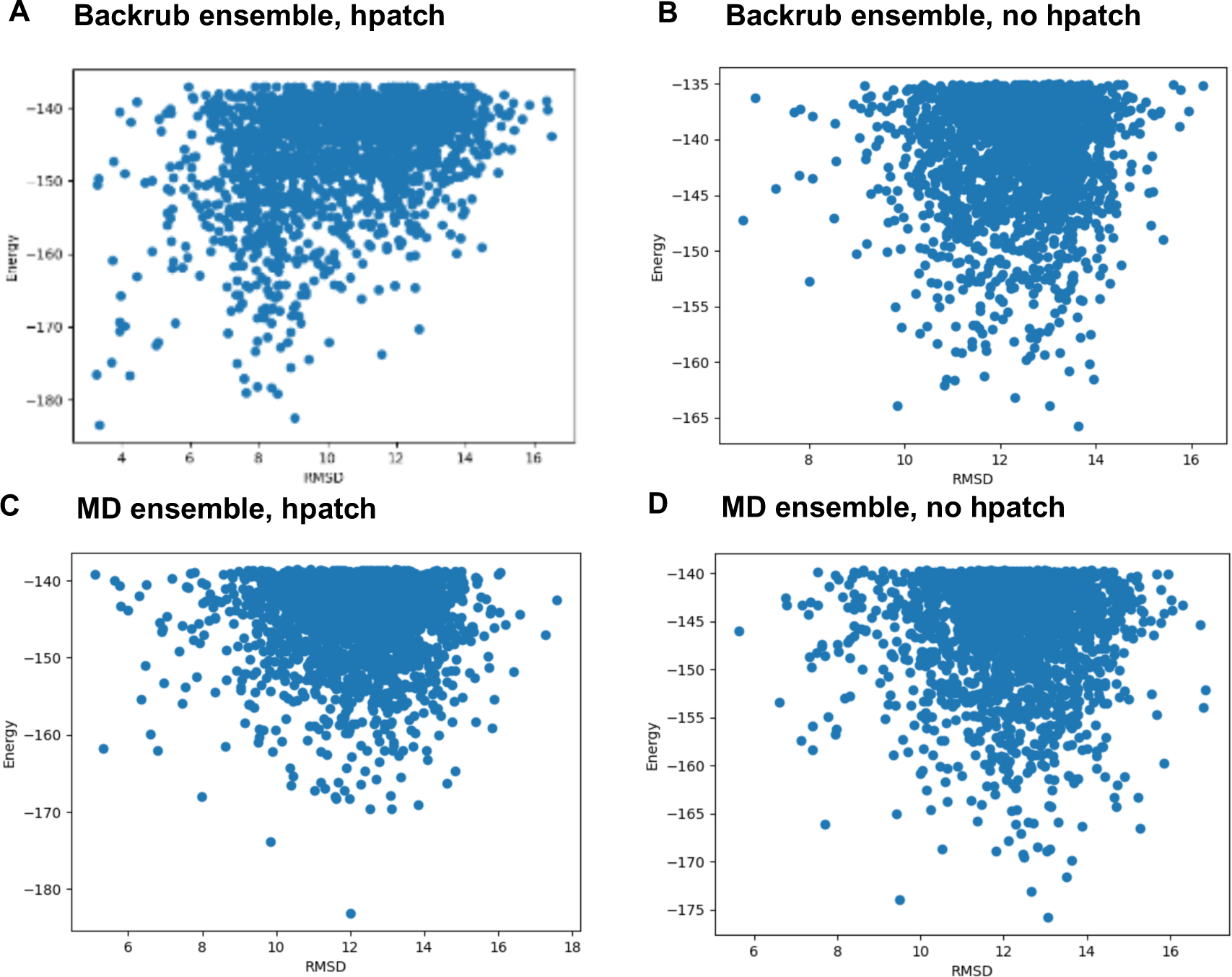
Forward folding profiles of MS designs. For each design, 30,000 protein models were predicted. The RMSD to the template structure was computed for the 1,000 top scoring models. The presence of dots in the bottom left corner of the Backrub ensemble with hpatch plot (A) indicates that EdaRose predicted low energy models within 4 Å from the target structure. For comparison, the Backrub ensemble without hpatch (B) and the MD ensembles with (C) and without (D) hpatch are shown.

**Fig. S11.**
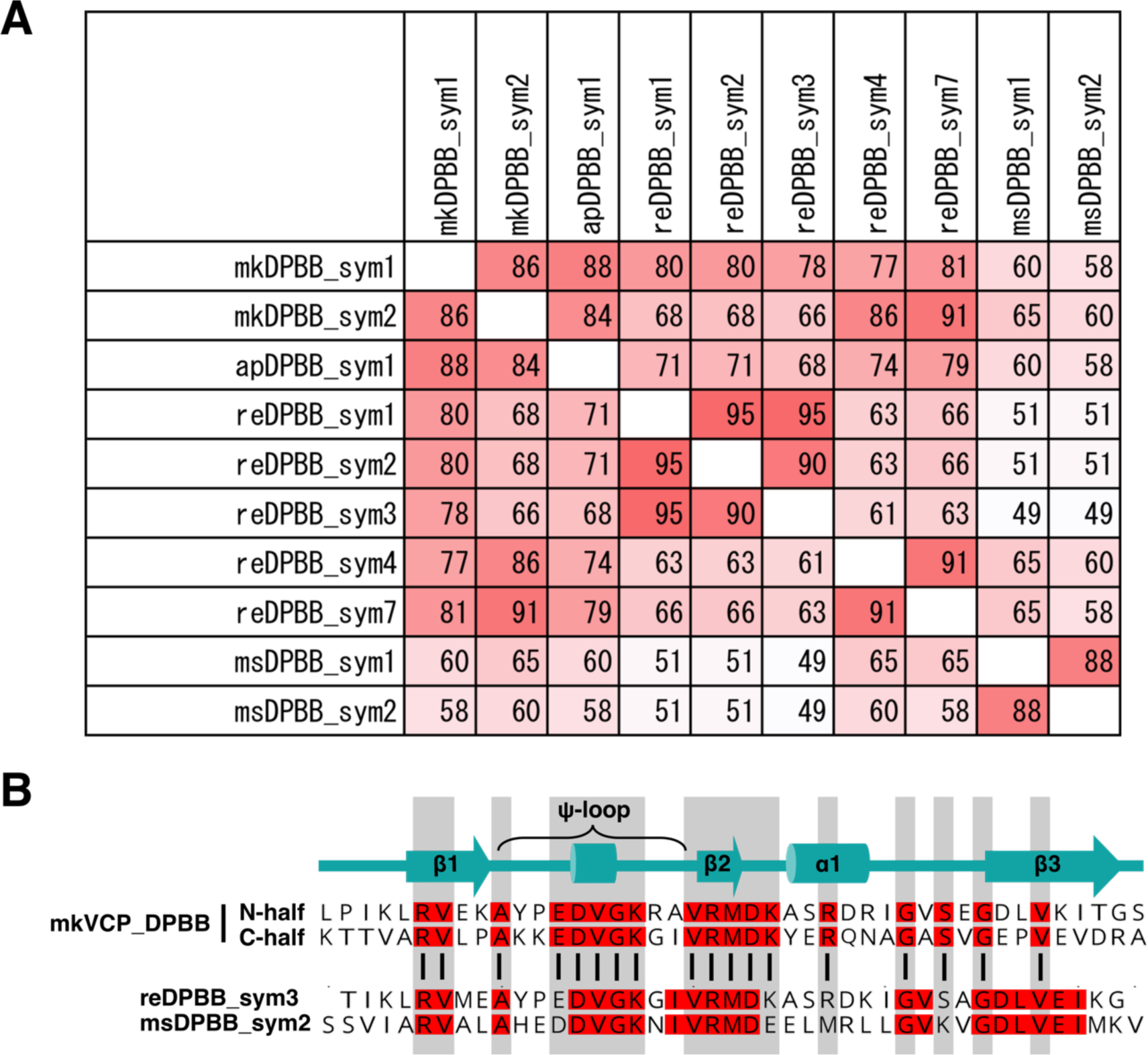
Sequence diversity in symmetrically designed DPBBs. (A) Sequence identities (%) between each pair of repeat units in the symmetrically designed DPBBs. msDPBB_sym1 and 2 share only 49% sequence identity with reDPBB_sym3. (B) Pairwise sequence alignment of the repeat units in reDPBB_sym3 and msDPBB_sym2 along with the starting template, mkVCP_DPBB. The symmetrically-conserved positions in the mkVCP_DPBB are highlighted in gray. Considering that both designs were created from the same starting model and should be biased toward being highly homologous to the original sequence, the potential diversity of the possible DPBB sequences could still be underestimated. If we compare the residues at the non-symmetric positions in the starting model, then reDPBB_sym3 and msDPBB_sym2 share only 26% sequence identity.

**Figure S12.**
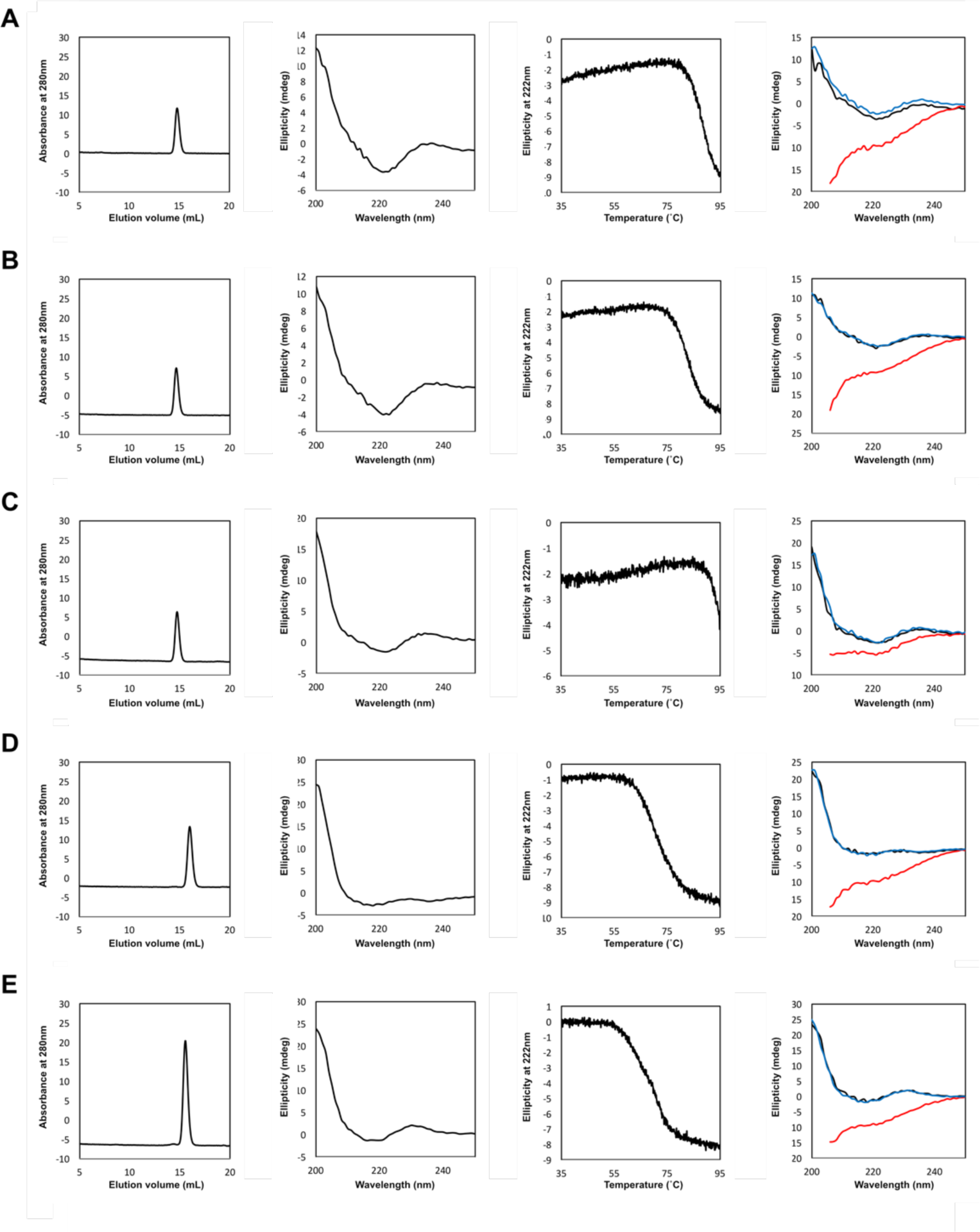
Experimental characterization of symmetrical DPBBs designed by the RE- design method. Size exclusion chromatography, CD spectra, denaturation curves, and comparisons of CD spectra at different temperatures (black: 35°C; red: 95°C; blue (refolding): 95°C à 35°C) for (A) reDPBB_sym1, (B) reDPBB_sym2, (C) reDPBB_sym3, (D) reDPBB_sym4, and (E) reDPBB_sym7 are shown in the panels from left to right.

**Figure S13.**
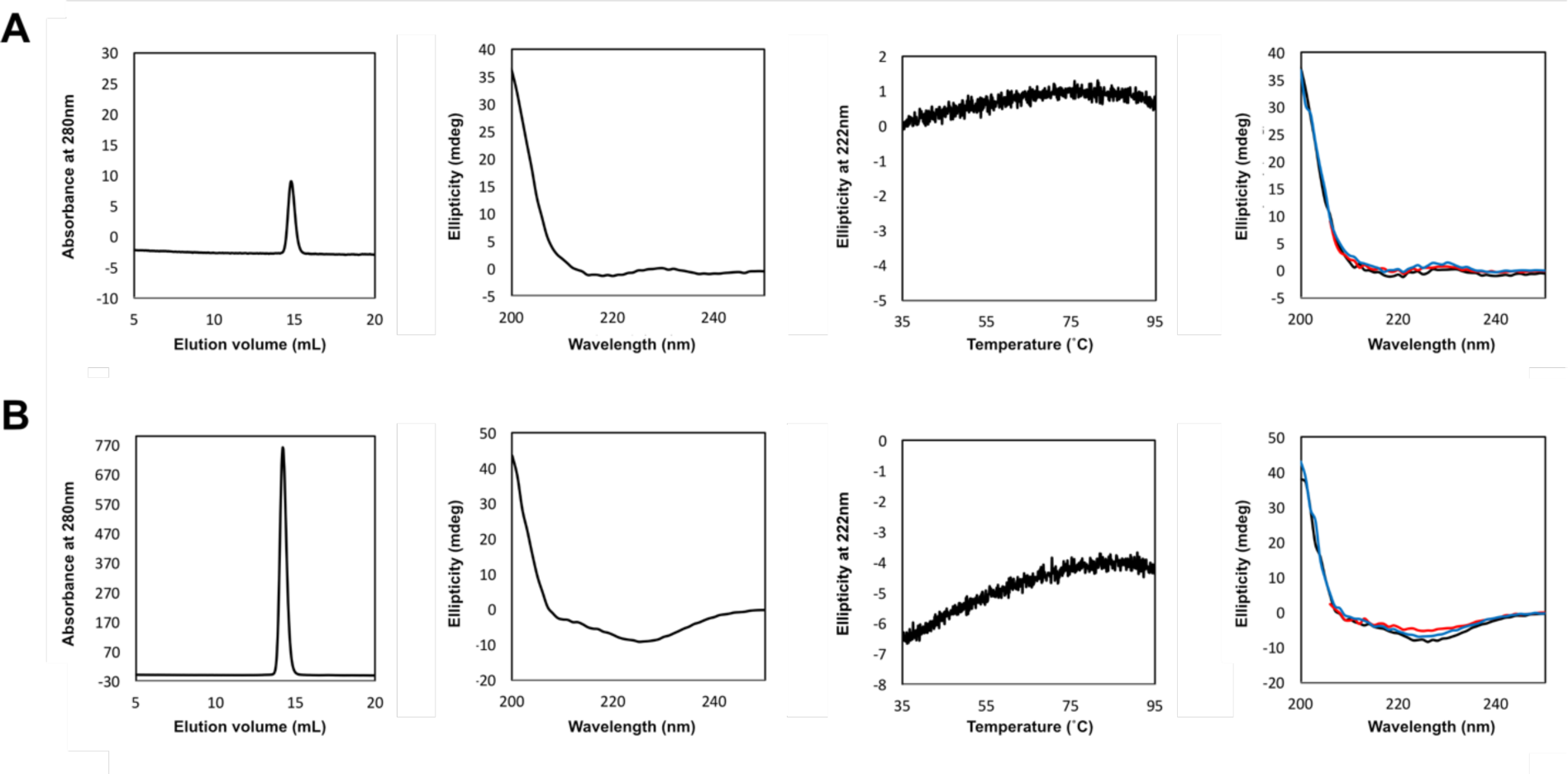
Experimental characterization of symmetrical DPBBs designed by the MS- design method. Size exclusion chromatography, CD spectra, denaturation curves, and comparisons of CD spectra at different temperatures (black: 35°C; red: 95°C; blue (refolding): 95°C à 35°C) for (A) msDPBB_sym1 and (B) msDPBB_sym2 are shown in the panels from left to right.

**Figure S14.**
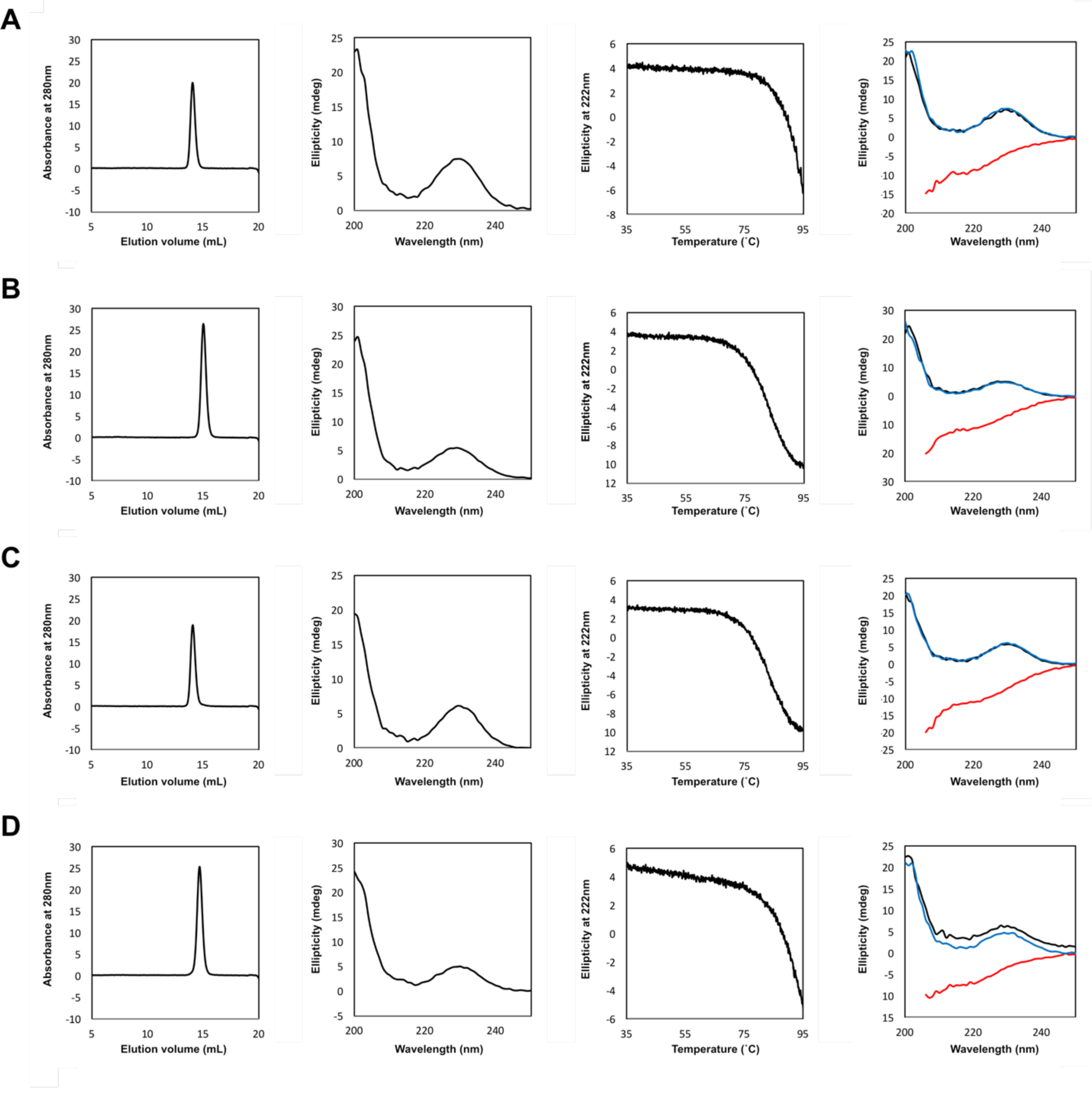
Experimental characterization of the halved fragments. Size exclusion chromatography, CD spectra, denaturation curves, and comparisons of CD spectra at different temperatures (black: 35°C; red: 95°C; blue (refolding): 95°C à 35°C) for (A) mk1h, (B) mk2h,(C) ap1h, and (D) ap2h are shown in the panels from left to right.

**Figure S15.**
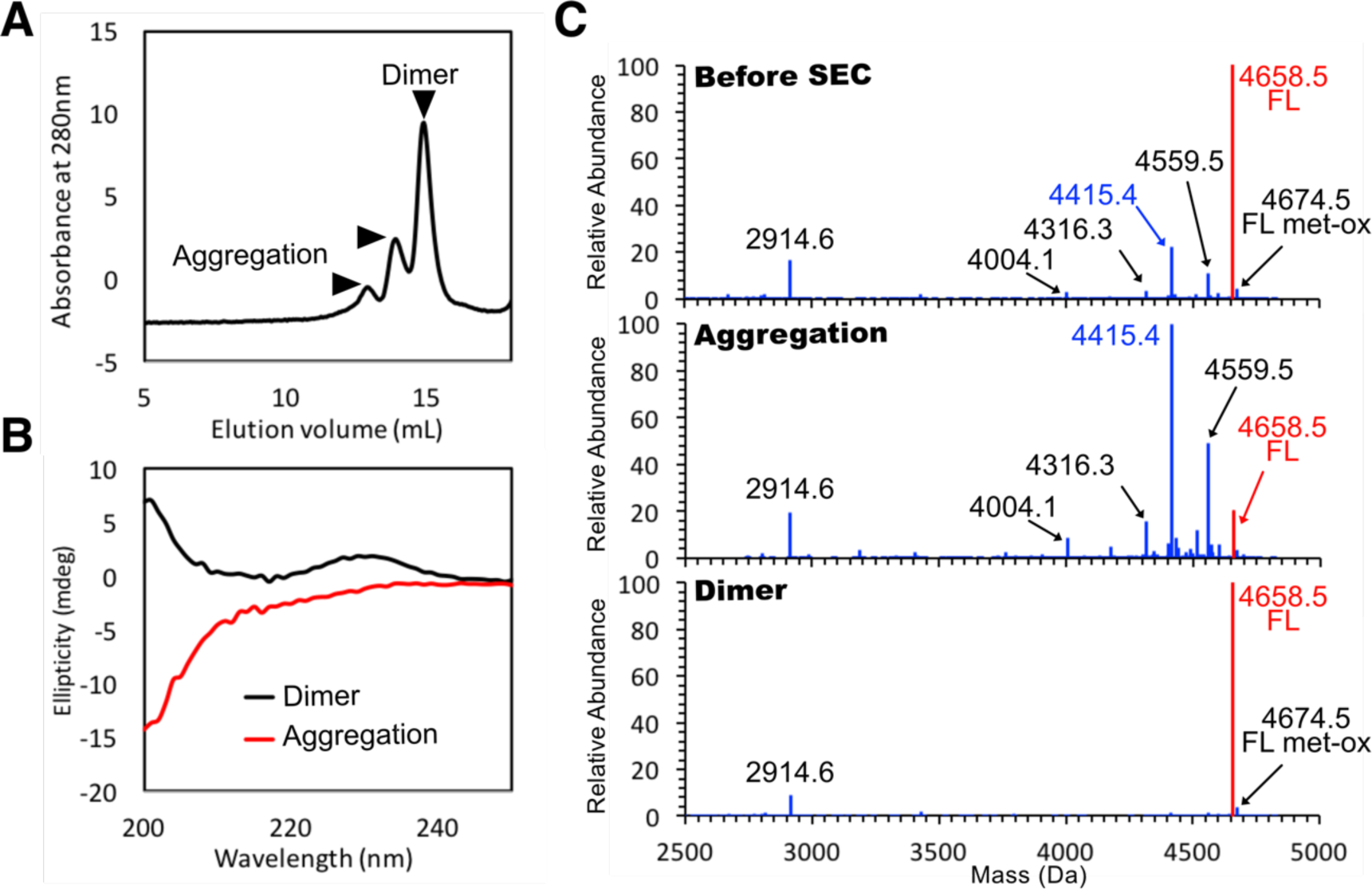
Characterization of the low-purity chemically-synthesized mk1h peptide (75.06%). (A) SEC analysis showing the aggregated and dimeric states of the dissolved mk1h peptide. (B) CD spectra indicating that the aggregated and dimer species separated in Fig. S12B, adopt random-coil and *α/β* structures, respectively. (C) The peptide species in the sample before and after the SEC purification were analyzed by LC/MS, and the deconvoluted mass spectra are shown. The labels for the full-length mk1h peptide (4658.5 Da) and the major contaminant peptide (4415.4 Da) are highlighted in red and blue, respectively. While most of the contaminant peptides were enriched in the aggregation fraction, the full-length mk1h was enriched in the dimer fraction.

**Figure S16.**
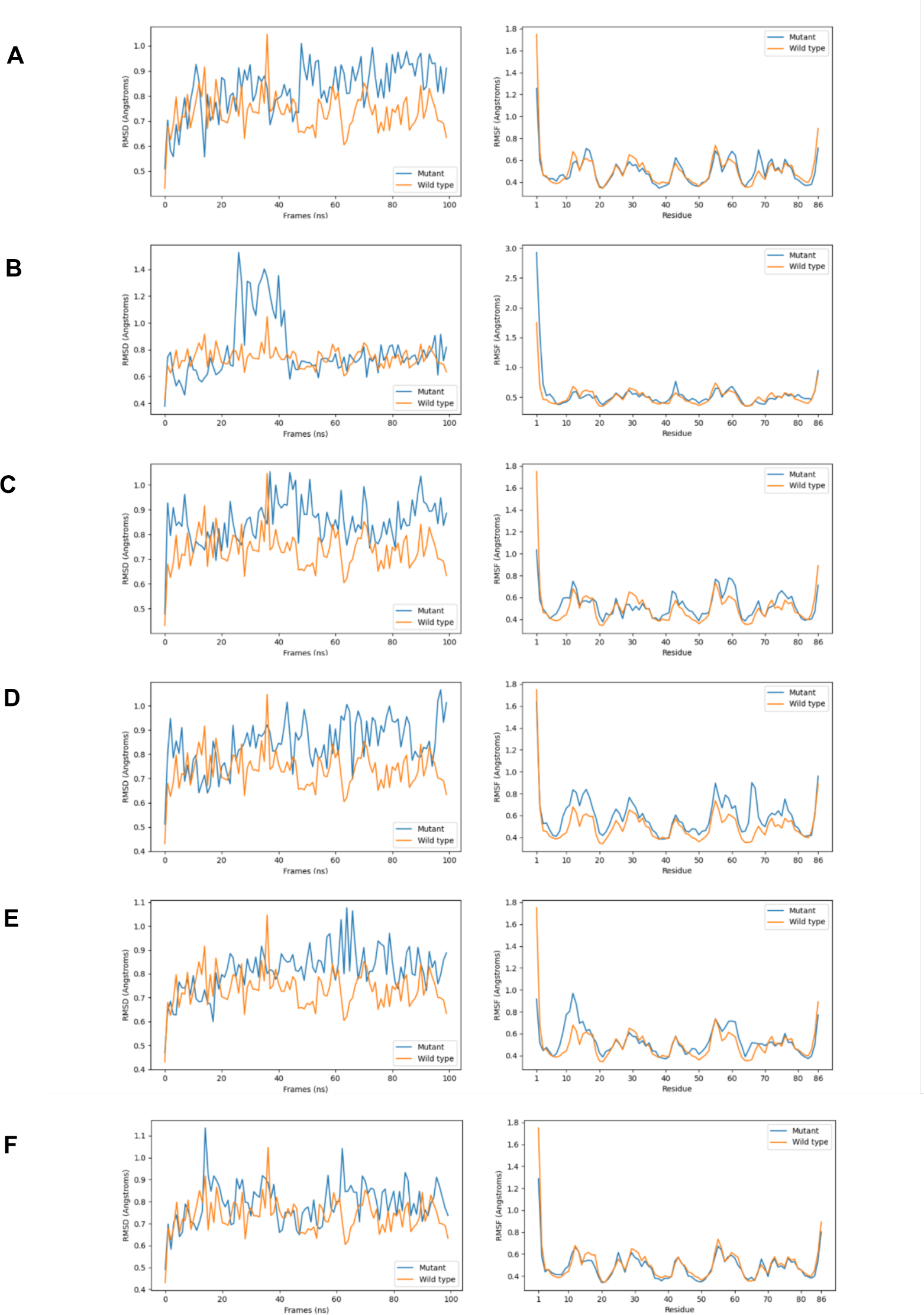
Molecular dynamics simulations of the mk2h_ΔI. (A), mk2h_ΔL (B), mk2h_ΔM (C), mk2h_ΔP (D), mk2h_ΔS (E), and mk2h_ΔY (F) mutants. The simulations were performed starting from structural models in the form of the linked repeats with 86 amino acid residues. The backbone RMSD to the reference structure along 100 ns of simulation is plotted on the left, and the backbone RMSF by residue averaged over 100 ns of simulation is plotted on the right. No large conformational change, except around the N-terminus, was observed in both the original (mk2h) and mutant designs.

**Figure S17.**
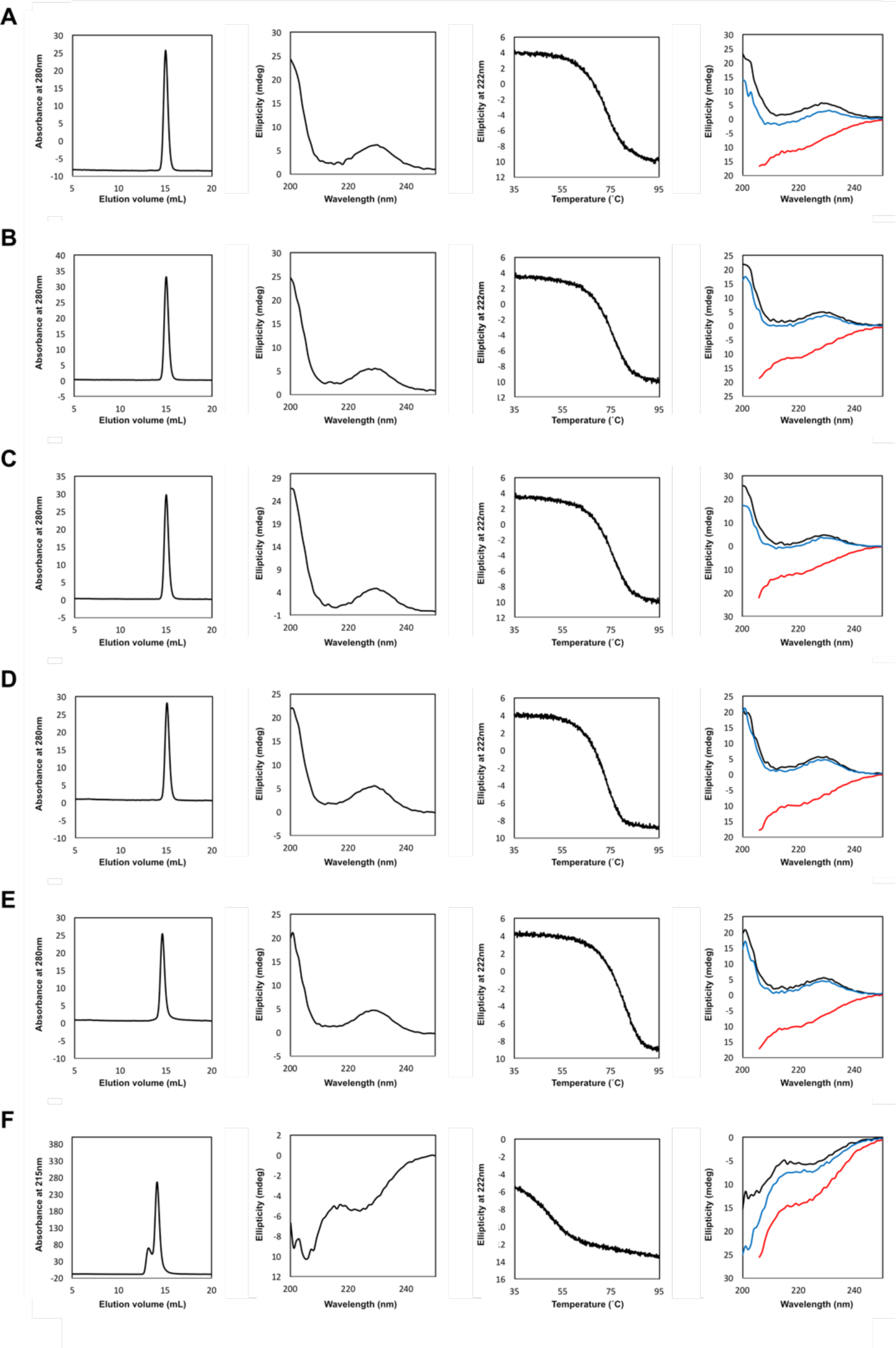
Experimental characterization of mk2h mutants containing 12 amino acid repertoires. Size exclusion chromatography, CD spectra, denaturation curves, and comparisons of CD spectra at different temperatures (black: 35°C; red: 95°C; blue (refolding): 95°C à 35°C) for (A) mk2h_ΔM, (B) mk2h_ΔI, (C) mk2h_ΔL, (D) mk2h_ΔP, (E) mk2h_ΔS, and (F) mk2h_ΔY are shown in the panels from left to right.

**Figure S18.**
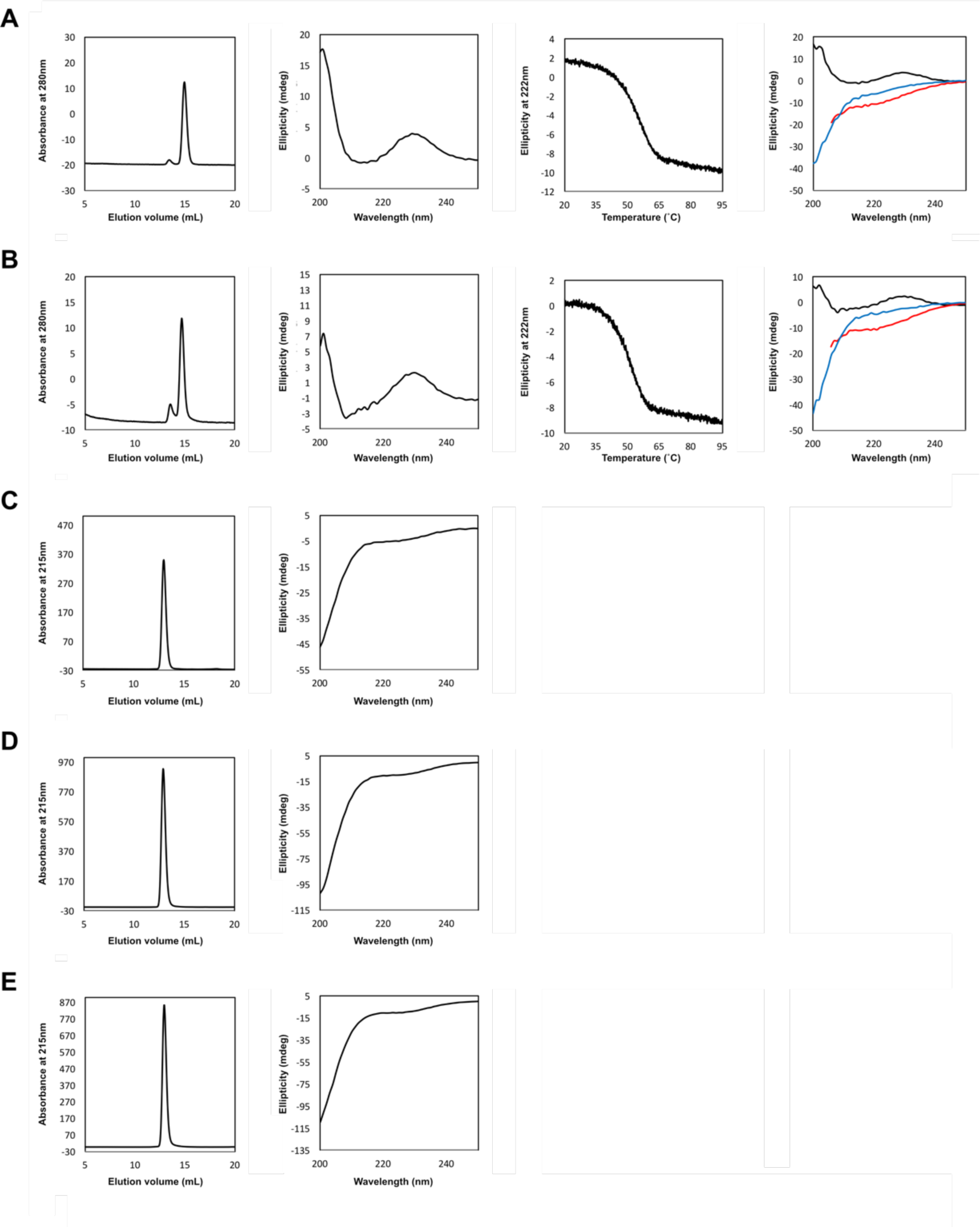
Experimental characterization of mk2h mutants containing 7, 8, or 10 amino acid repertoires. Size exclusion chromatography, CD spectra, denaturation curves, and comparisons of CD spectra at different temperatures (black: 10°C; red: 95°C; blue (refolding): 95°C à 10°C) for (A) mk2h_ΔMIL, (B) mk2h_ΔMILPS, (C) mk2h_ΔMILPY, (D) mk2h_ΔMILSY, and (E) mk2h_ΔMILPYS are shown in the panels from left to right.

**Fig. S19.**
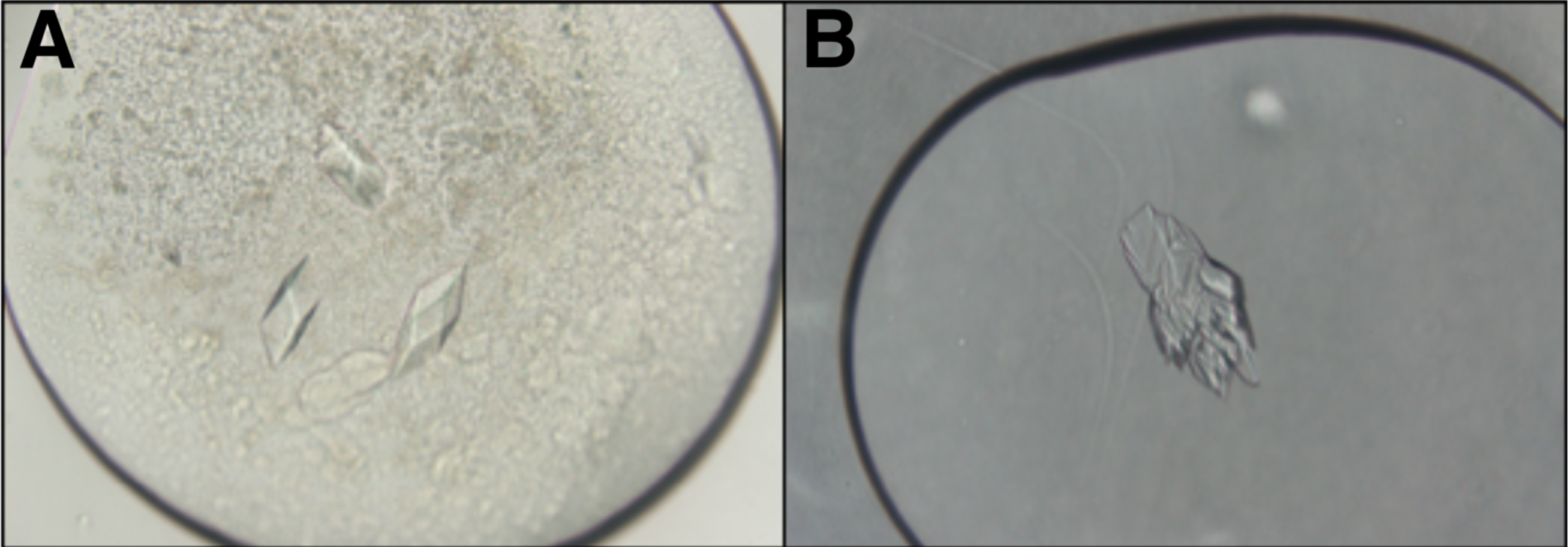
Crystals of mk2h_ΔMILPYS. The crystals were obtained under two different conditions containing (A) an undissolved suspension of the chemically-synthesized peptide in 3 M sodium malonate and (B) the dissolved peptide in 2.1 M DL-malic acid, pH 7.0. The determined crystal structures are shown in Fig. 4D and Fig. S6K, respectively.

**Figure S20.**
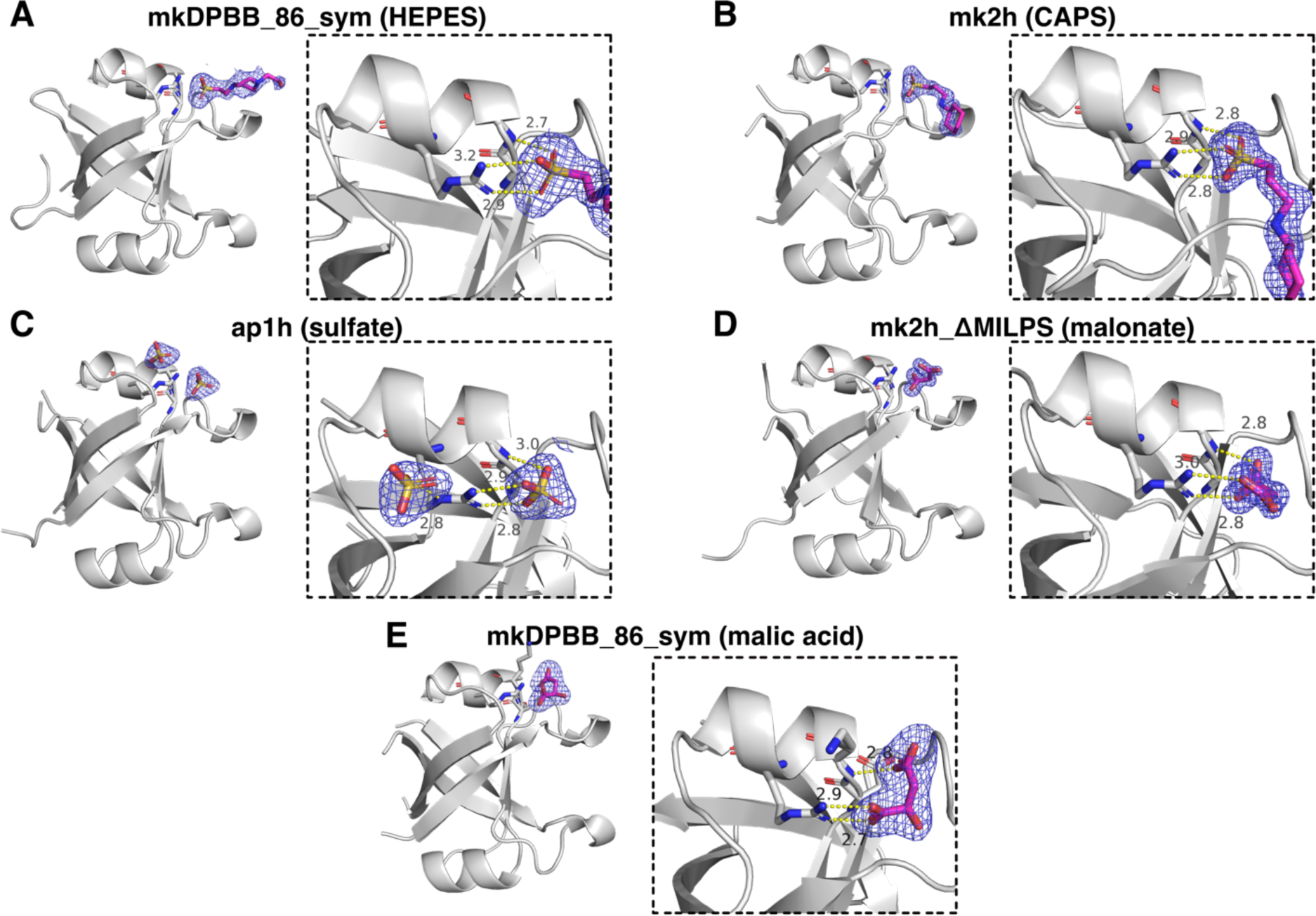
Positively-charged pockets in designed DPBBs occupied by negatively-charged ligands. (A–E) Crystal structures and close-up views of the conserved positively-charged pockets of designed DPBB domains. (A) mkDPBB_sym_84, (B) mk2h, (C) ap1h, (D) mk2h_ΔMILPS, and (E) mk2h_ΔMILPYS interactions with HEPES, CAPS, sulfate ion, malonate ion, and malic acid, respectively, at the positively charged pocket around their *α*-helices. The N-H group at the N-terminal peptide bond of the *α*1 helix and the arginine residue positioned in the middle of the *α*1 helix (Arg30 in mk2h_ΔMILPYS) form salt-bridges with the negatively-charged ligands. This observation supports the idea that the ancestral DPBB proteins composed of the repeated sequence or halved fragments could have functioned as cofactor- or nucleic acid-binding proteins, like their extant descendants (*43, 44*).

**Algorithm S1.**
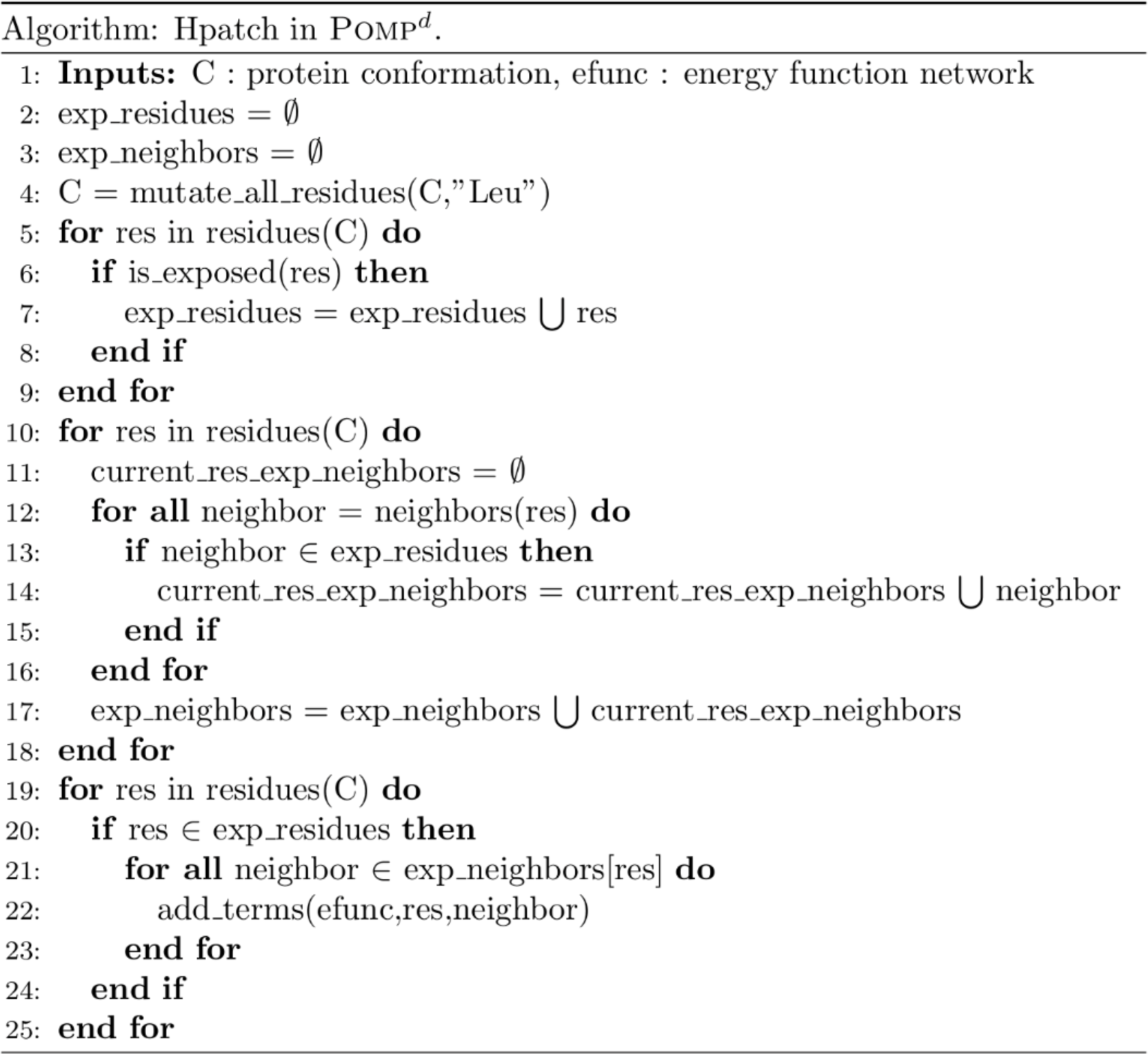
Hpatch procedure. Exposed surface residues are calculated with pyRosetta, after mutating the whole protein to LEU in order to ensure the equal treatment of all residue positions. For each residue, the list of all its exposed neighbors is computed. Finally, the neighboring hydrophobic pairs at the surface are forbidden with new energy terms.

**Table S1.**
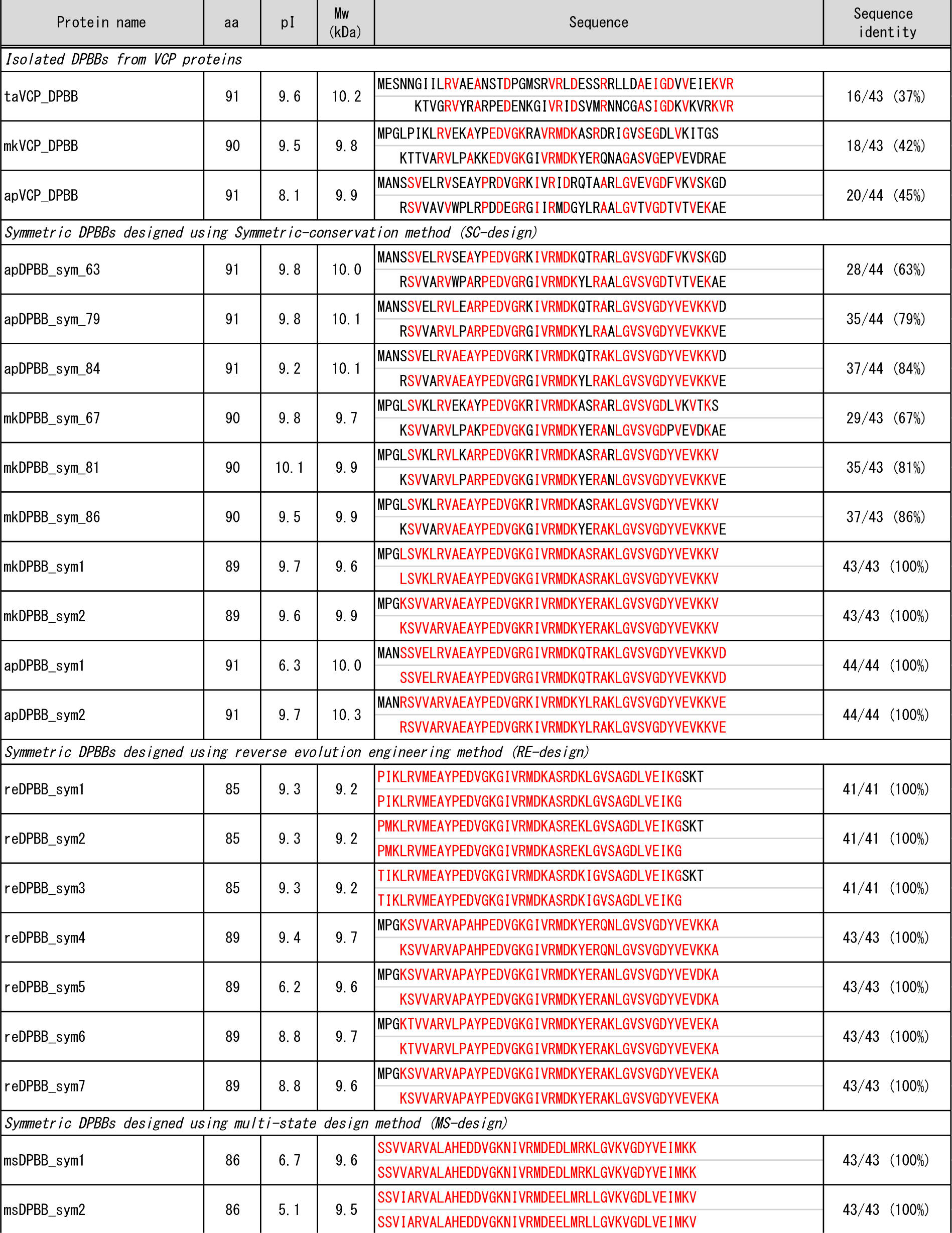
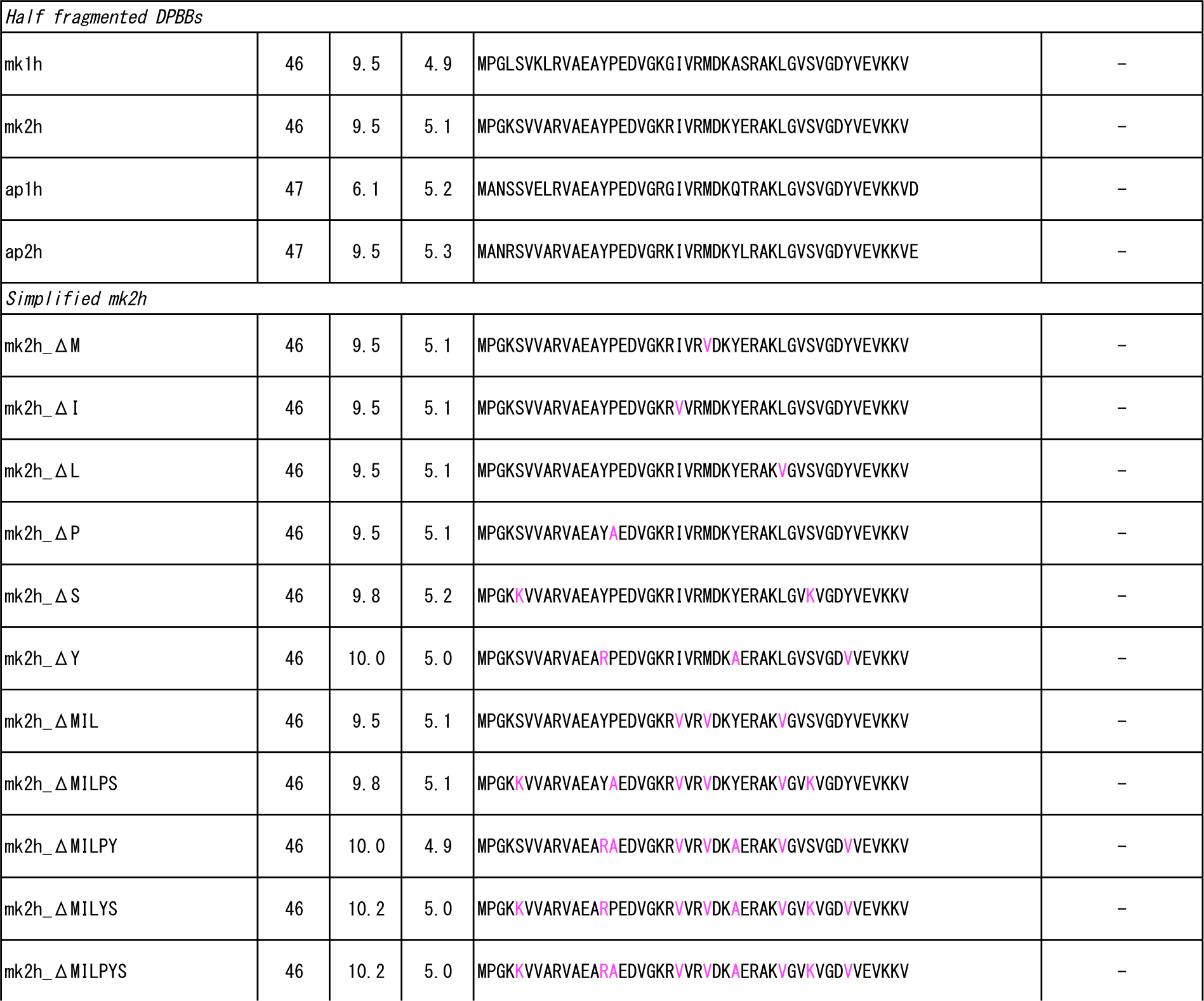
Sequences and internal identities of the natural and designed DPBB domains.

**Table S2.**
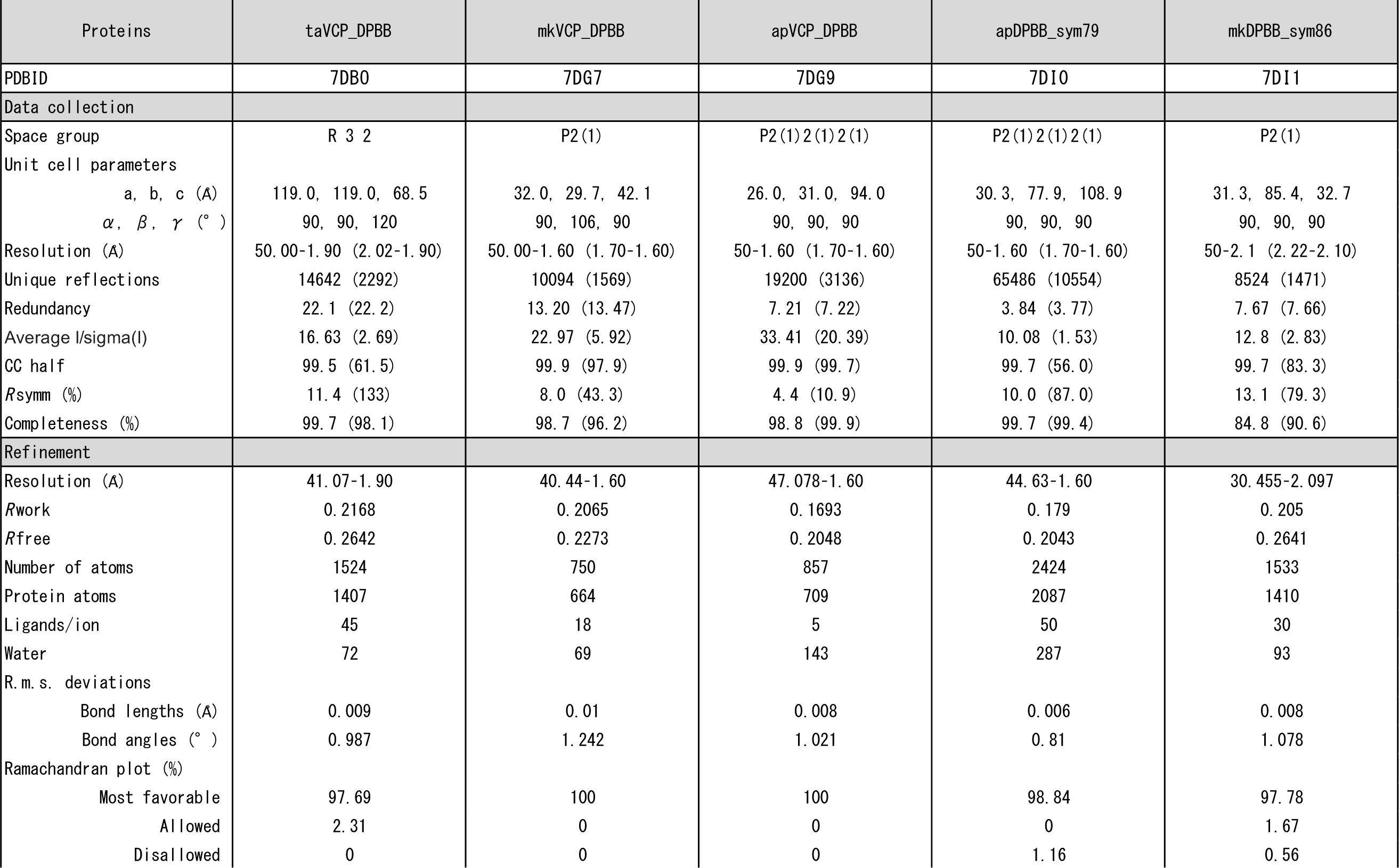
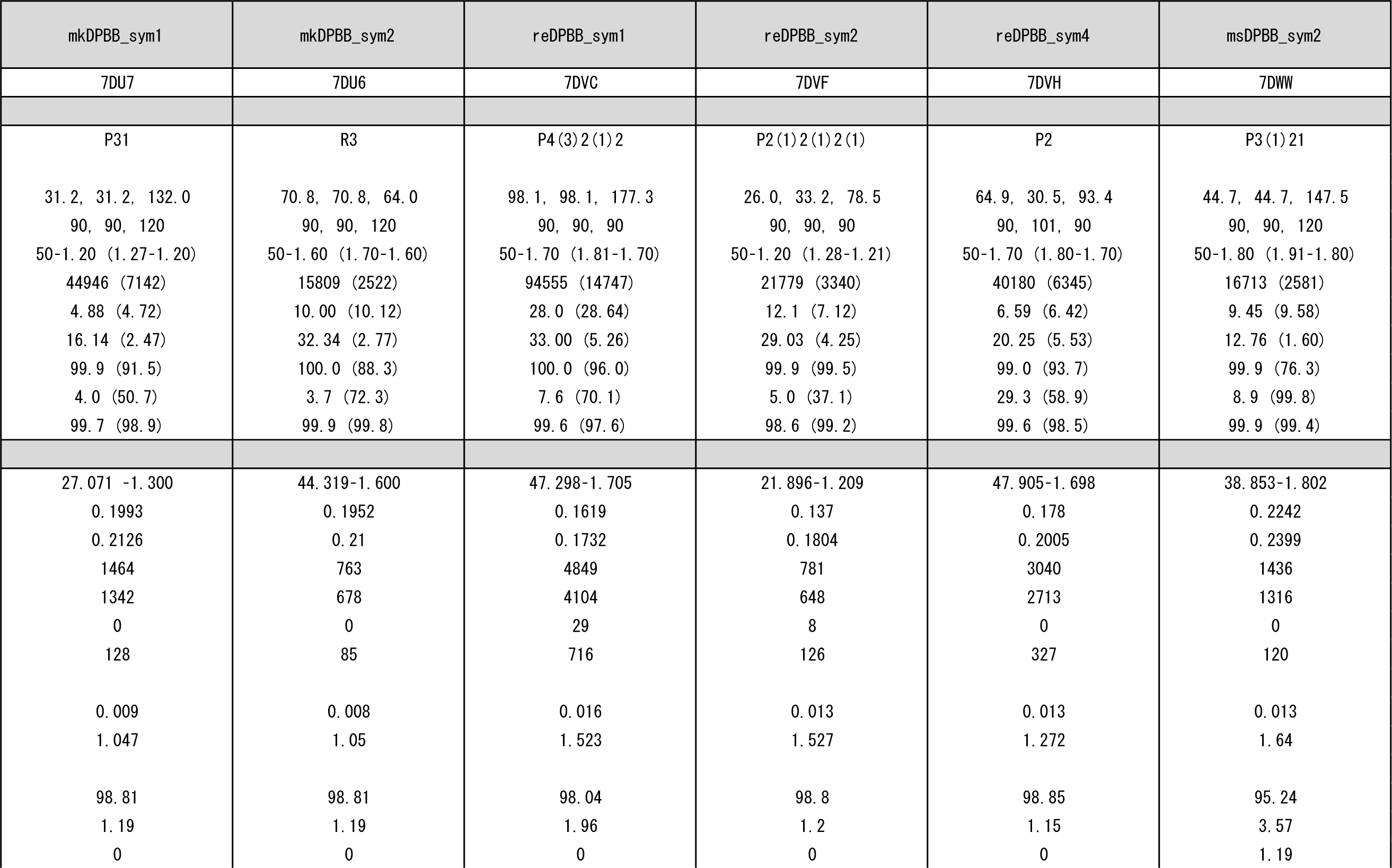
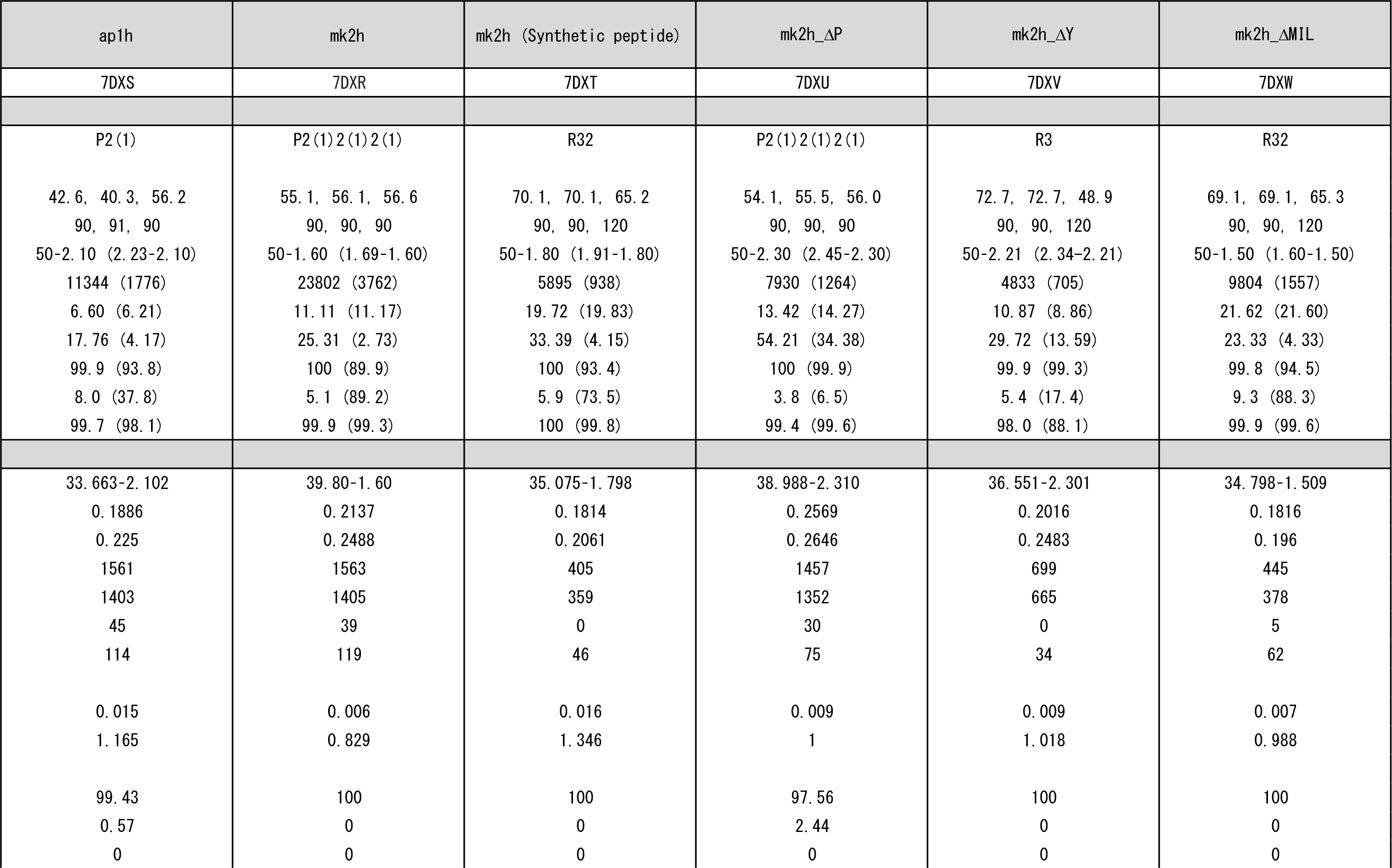
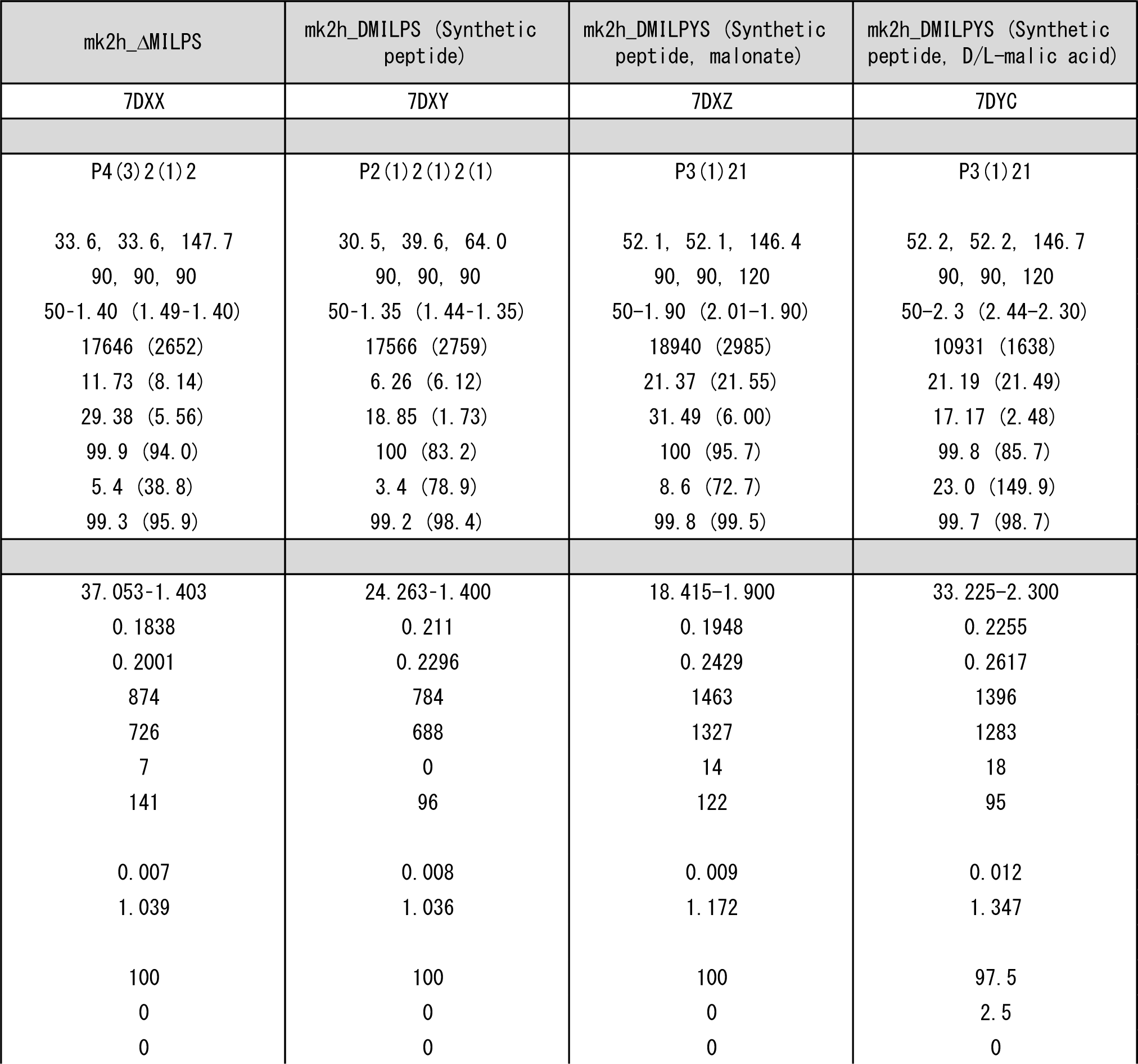
Data collection and refinement statistics

**Table S3.**
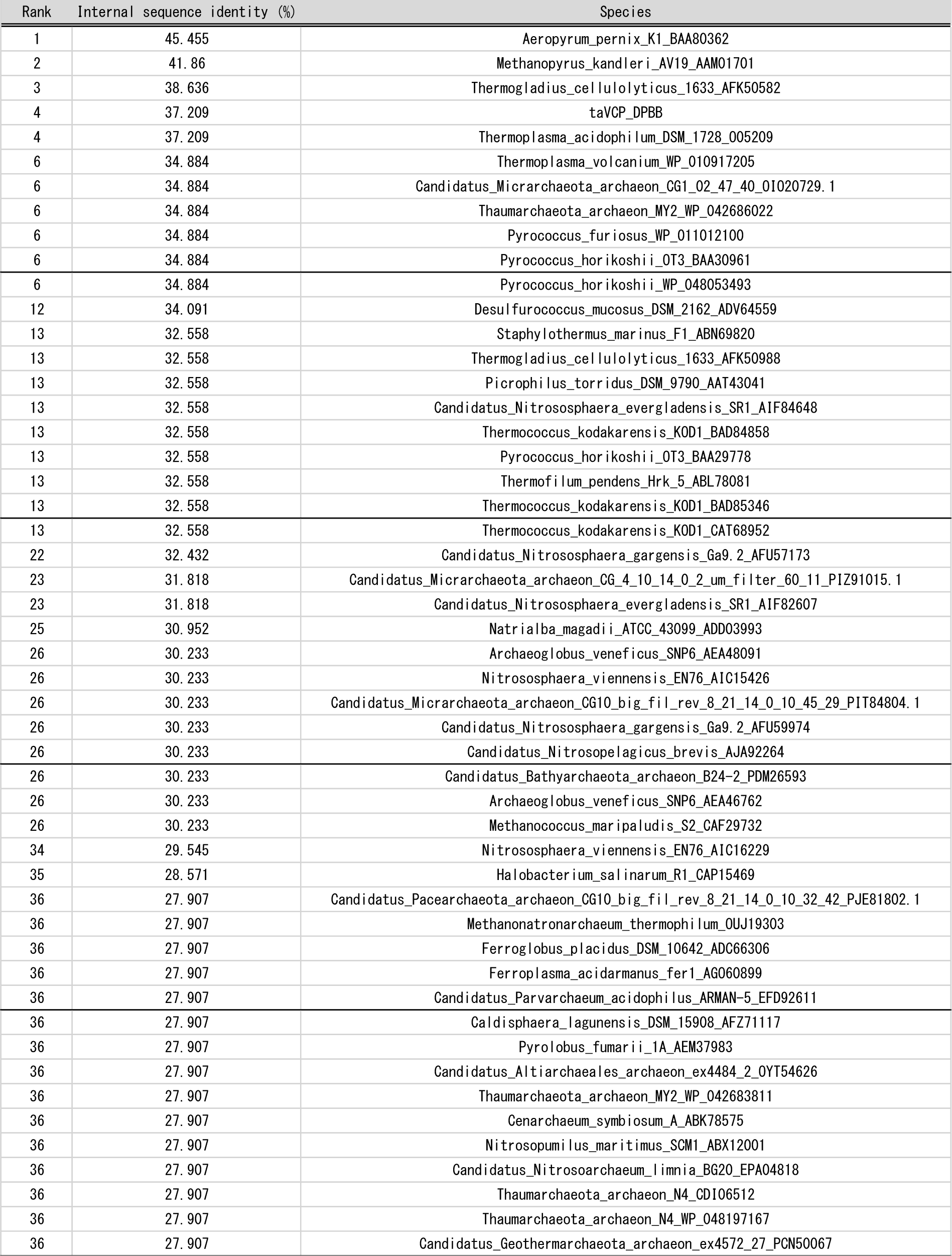
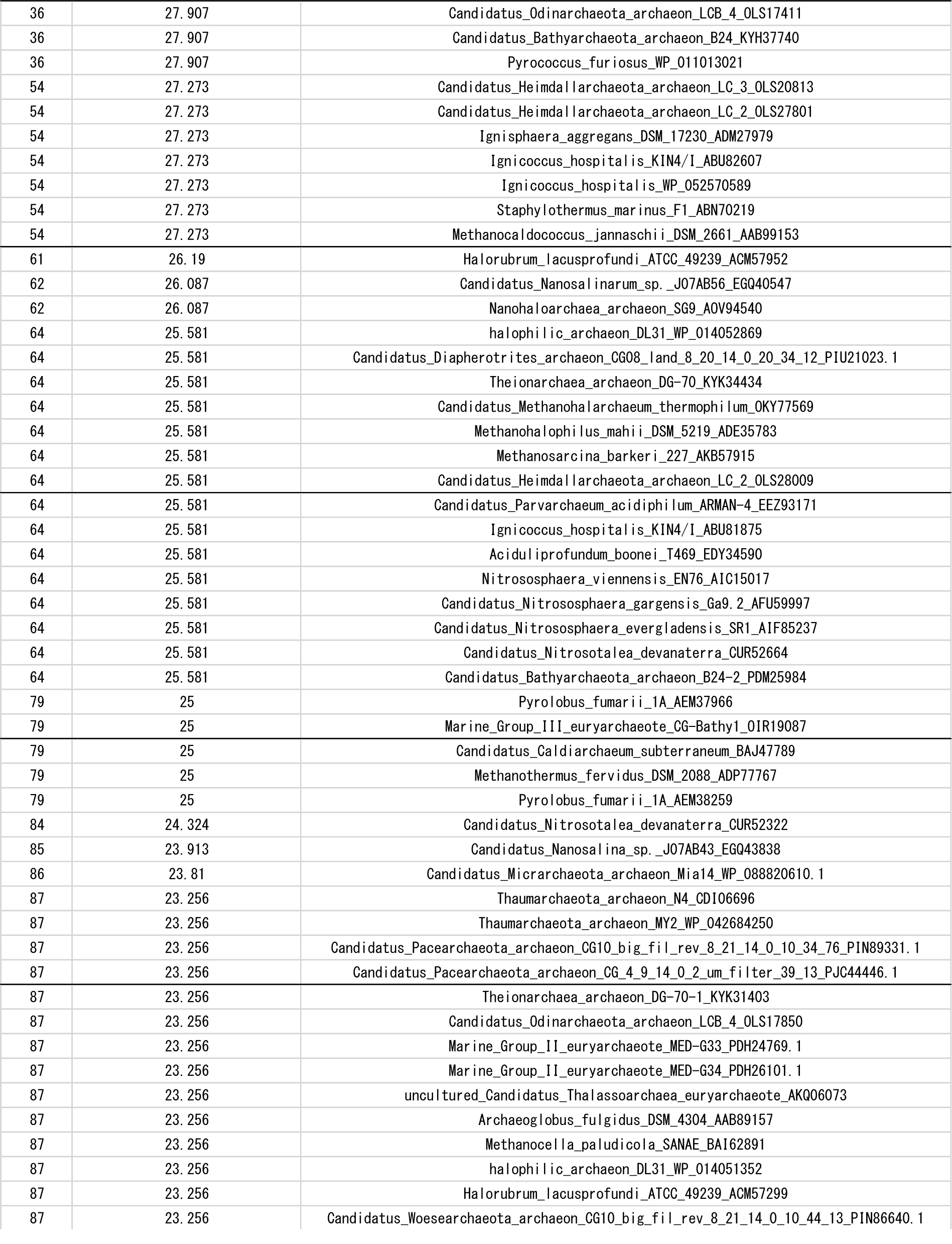
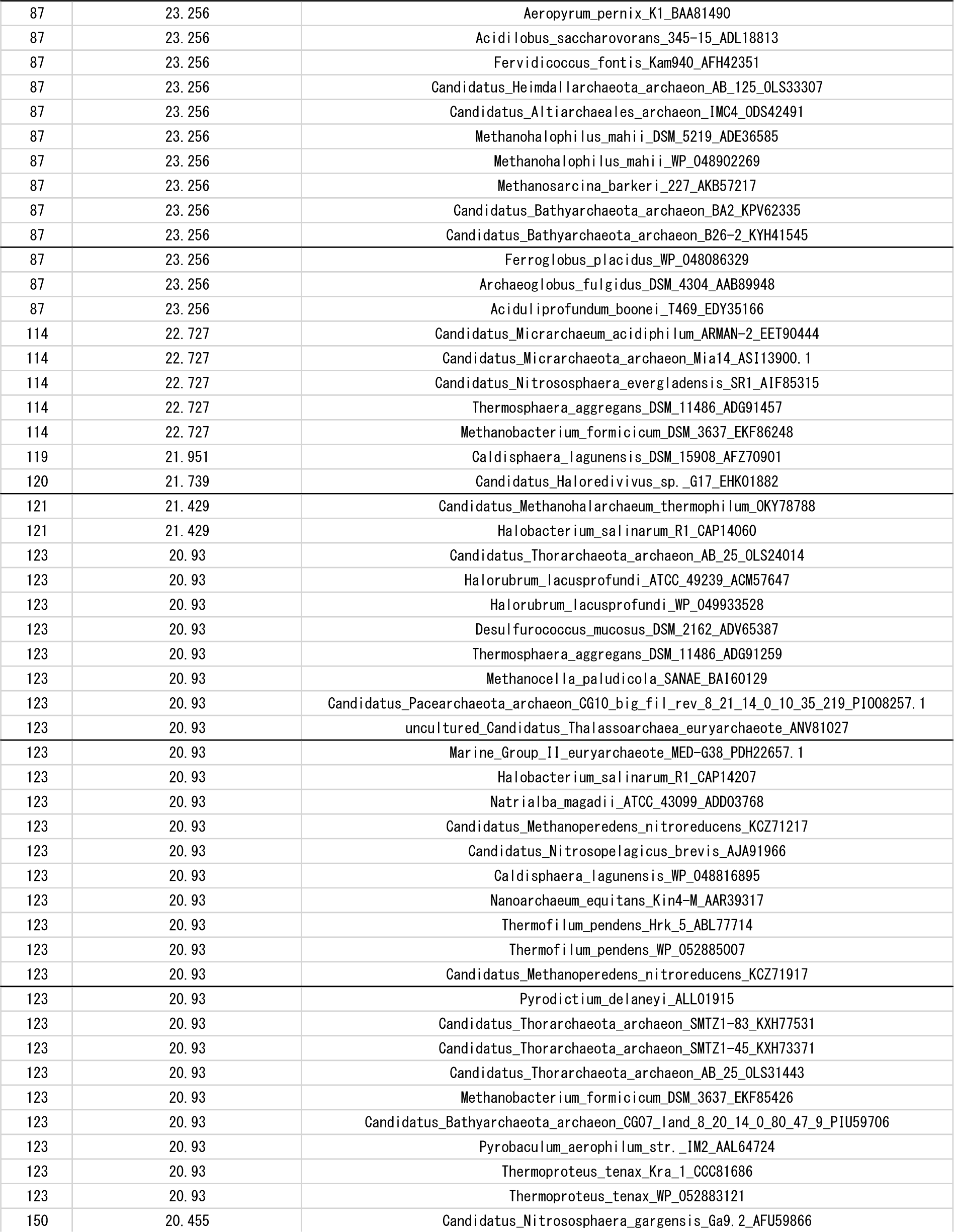
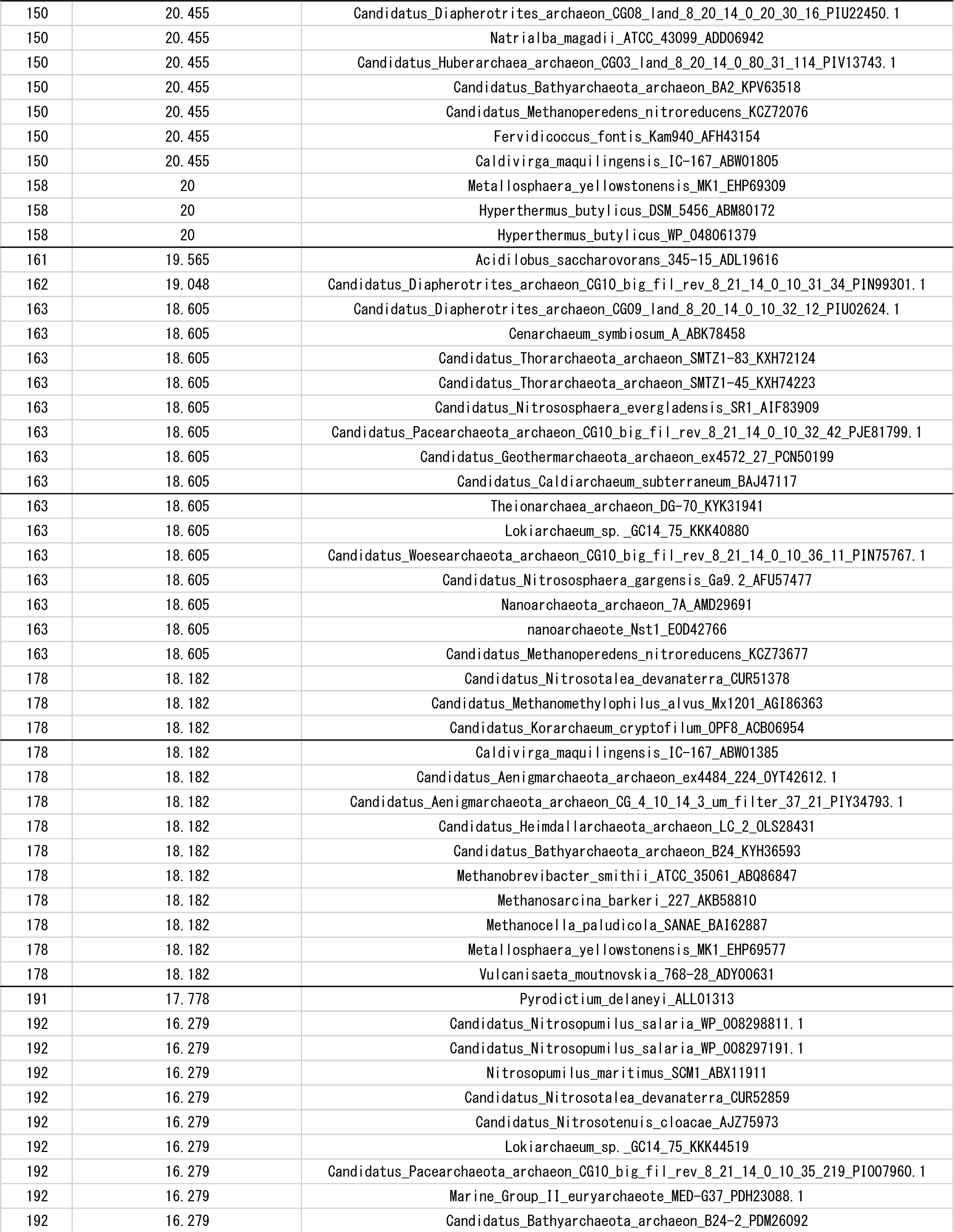
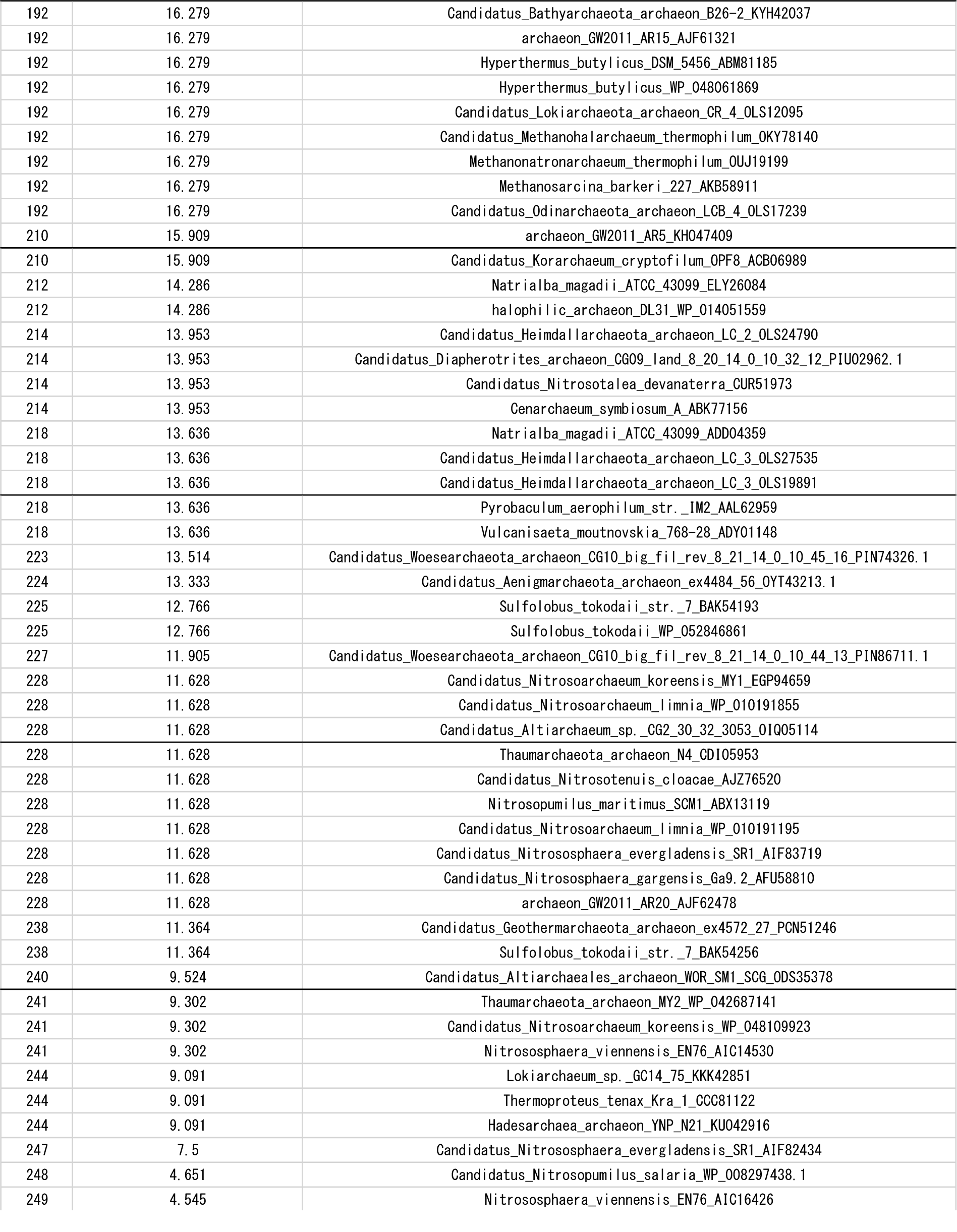
Ranking of VCP_DPBBs based on internal sequence identity.

**Table S4.**
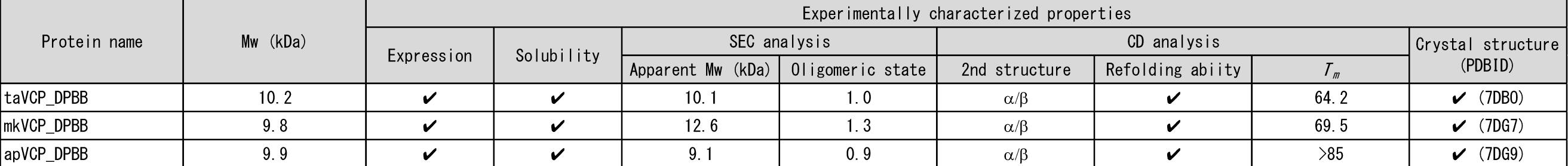
Experimental characterization of native DPBBs.

**Table S5.**
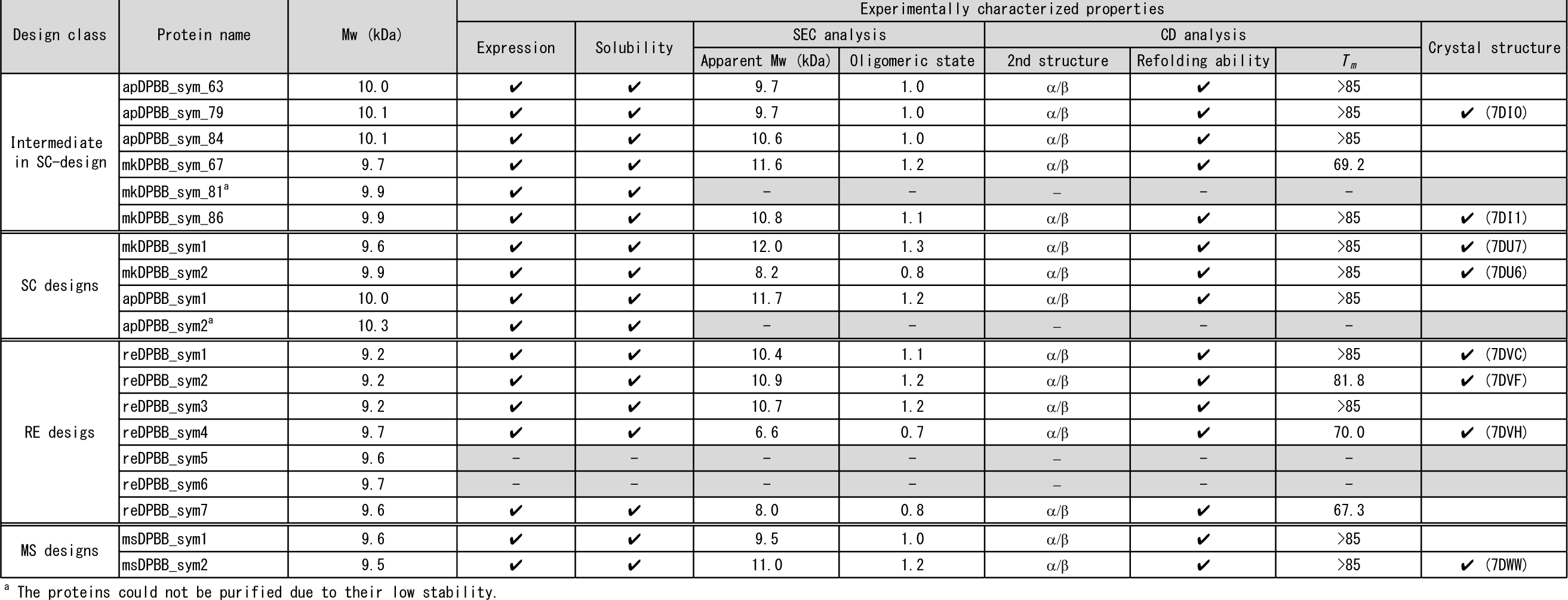
Experimental characterization of designed symmetric DPBBs.

**Table S6.**
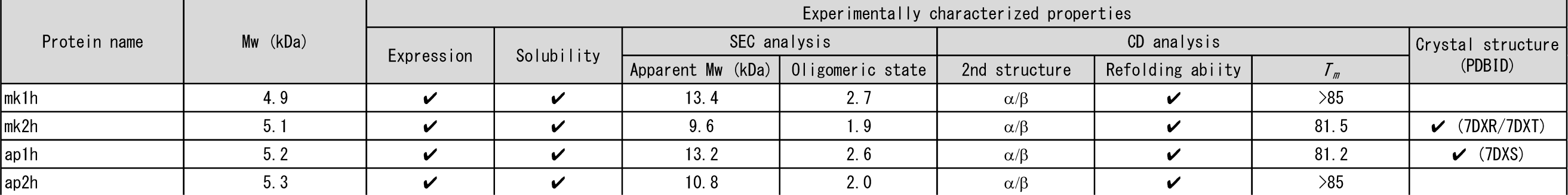
Experimental characterization of half-fragmented DPBBs

**Table S7.** Crystallization conditions for the synthetic mk2h peptide.

|  | Reservoir solution (RS) |
| --- | --- |
| 1 | 1000mM Potassium sodium tartrate, 100 mMimidazole/hydrochloric acid pH8.0, 200 mM Sodium chloride |
| 2 | 400mM Sodium phosphate monobasic/1600mM Potassium phosphate dibasic, 100mM Imidazole/hydrochloric acid pH8.0, 200mM Sodium chloride |
| 3 | 1000mM Sodium citrate tribasic, 100mM CHES/Sodium hydroxide pH9.5 |
| 4 | 1000mM Sodium citrate tribasic, 100mM Tris base/hydrochloric acid pH7.0, 200mM Sodium chloride |
| 5 | 800mM Sodium phosphate monobasic/1200mM Potassium phosphate dibasic, 100mM Sodium acetate/Acetic acid pH4.5 |
| 6 | 10% (w/v) PEG 3000, 100mM Sodium phosphate dibasic/Citric acid pH4.2, 200mM Sodium chloride |
| 7 | 20% (v/v) Jeffamine M-600, 100 mM HEPES/Sodium hydroxide pH7.5 |

**Table S8.** Quantification of peptide species by LC/MS.

| mk1h peptide |  |  |  |  |  |  |
| --- | --- | --- | --- | --- | --- | --- |
| Monoisotopic mass (Da) | Before SEC |  | Aggregation frac. |  | Dimer frac. |  |
|  | Sum Intensity | Relative Abundance (%) | Sum Intensity | Relative Abundance (%) | Sum Intensity | Relative Abundance (%) |
| 2914.6 | 2233357 | 16.7 | 1538848 | 19.3 | 1666535 | 8.8 |
| 3428.9 | 285446 | 2.1 | 50208 | 0.6 | 362748 | 1.9 |
| 4004.1 | 359518 | 2.7 | 673725 | 8.5 | 11170 | 0.1 |
| 4174.3 | 110288 | 0.8 | 388998 | 4.9 | ND<sup>a</sup> |  |
| 4316.4 | 419224 | 3.1 | 1242401 | 15.6 | 23622 | 0.1 |
| 4403.4 | 181392 | 1.4 | 519933 | 6.5 | ND<sup>a</sup> |  |
| 4408.5 | 51985 | 0.4 | 357992 | 4.5 | ND<sup>a</sup> |  |
| 4415.5 | 2939688 | 22.0 | 7965074 | 100.0 | 271045 | 1.4 |
| 4431.5 | 287595 | 2.2 | 697384 | 8.8 | 46841 | 0.2 |
| 4443.4 | 97978 | 0.7 | 358958 | 4.5 | 10669 | 0.1 |
| 4514.5 | 266774 | 2.0 | 950075 | 11.9 | 21476 | 0.1 |
| 4559.5 | 1436130 | 10.7 | 3926160 | 49.3 | 218946 | 1.2 |
| 4571.5 | 184029 | 1.4 | 471198 | 5.9 | 97616 | 0.5 |
| 4601.6 | 323717 | 2.4 | 455629 | 5.7 | 19369 | 0.1 |
| **4658.5 (mk1h FL)** | **13366797** | **100.0** | **1608175** | **20.2** | **18906107** | **100.0** |
| 4674.5 (mk1h FL met_OX) | 558788 | 4.2 | 262902 | 3.3 | 719772 | 3.8 |
| mk2h peptide |  |  |  |  |  |  |
| Monoisotopic mass (Da) | Before SEC |  | Aggregation frac. |  | Dimer frac. |  |
|  | Sum Intensity | Relative Abundance (%) | Sum Intensity | Relative Abundance (%) | Sum Intensity | Relative Abundance (%) |
| 1010.6 | 299575 | 2.7 | 370555 | 2.6 | 198979 | 1.2 |
| 3575.0 | 382652 | 3.5 | 1076537 | 7.5 | 13218 | 0.1 |
| 3600.9 | 601648 | 5.5 | 28685 | 0.2 | 70043 | 0.4 |
| 3662.0 | 621514 | 5.6 | 72742 | 0.5 | 814803 | 5.0 |
| 4321.4 | 216421 | 2.0 | 337394 | 2.4 | 4246 | 0.0 |
| 4477.5 | 277006 | 2.5 | 600300 | 4.2 | ND<sup>a</sup> |  |
| 4592.5 | 311937 | 2.8 | 993232 | 6.9 | ND<sup>a</sup> |  |
| 4620.5 | 150073 | 1.4 | 387705 | 2.7 | 35378 | 0.2 |
| 4633.6 | 225325 | 2.0 | 604106 | 4.2 | ND<sup>a</sup> |  |
| 4658.6 | 306598 | 2.8 | 17624 | 0.1 | 10857 | 0.1 |
| 4720.6 | 210070 | 1.9 | 589808 | 4.1 | 10318 | 0.1 |
| 4748.6 | 6001828 | 54.5 | 14336914 | 100.0 | 436117 | 2.7 |
| 4764.6 | 442905 | 4.0 | 688135 | 4.8 | 104955 | 0.6 |
| 4819.6 | 241081 | 2.2 | 301508 | 2.1 | 180023 | 1.1 |
| **4835.6 (mk2h FL)** | **11002630** | **100.0** | **850392** | **5.9** | **16360720** | **100.0** |
| 4851.6 (mk2h FL met_OX) | 524838 | 4.8 | 150537 | 1.0 | 1079166 | 6.6 |
| 4862.6 | 130898 | 1.2 | 285408 | 2.0 | ND<sup>a</sup> |  |

<sup>a</sup> The peptide could not be detected by LC/MS.

**Table S9.**
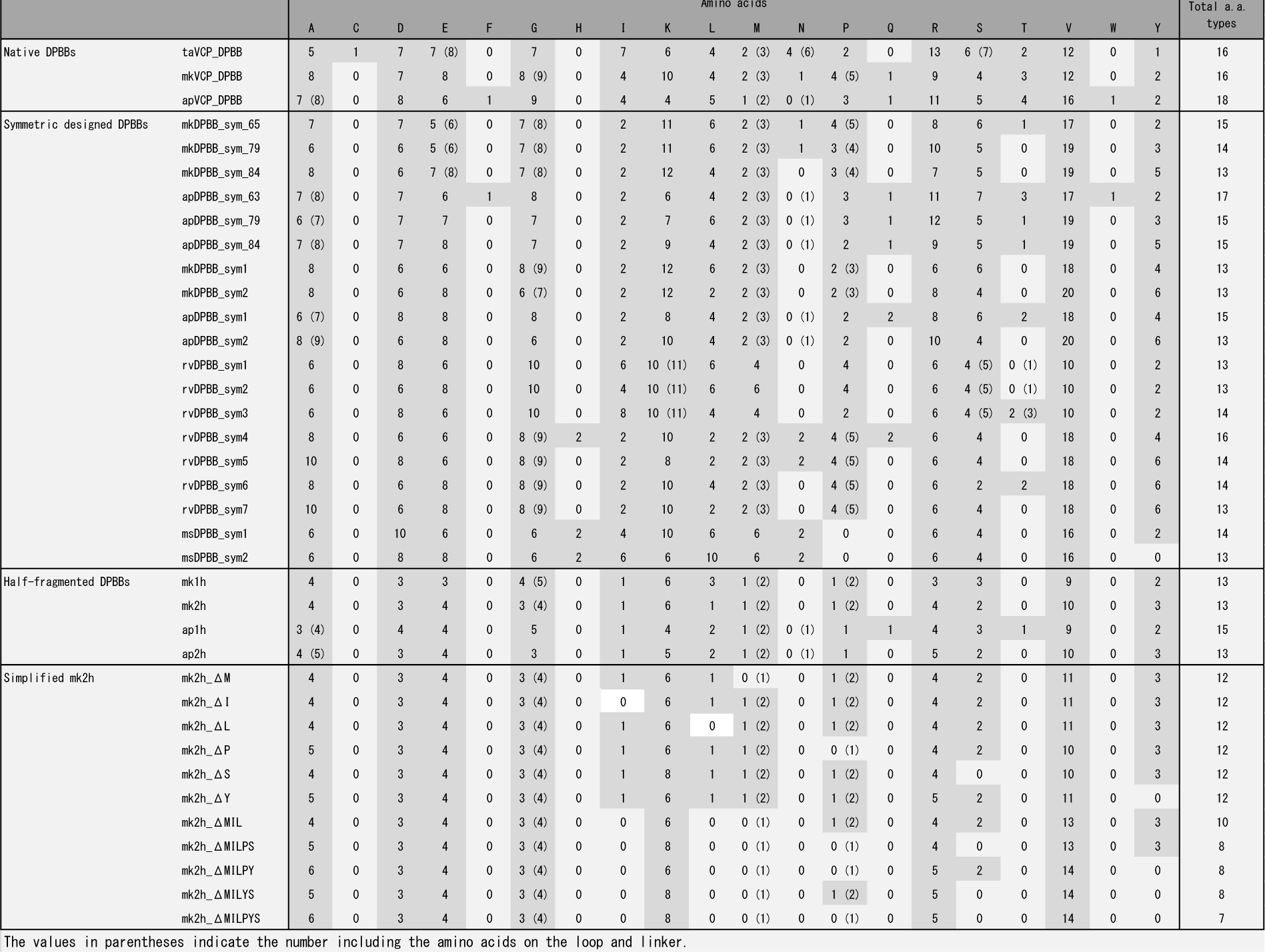
Amino acid usage in the designed DPBBs.

**Table S10.** Experimental characterization of simplified DPBBs

| Protein name | Mw (kDa) | Expression | Solubility | Experimentally characterized properties |  |  |  |  | Crystal structure (PDBID) |
| --- | --- | --- | --- | --- | --- | --- | --- | --- | --- |
|  |  |  |  | SEC analysis |  | CD analysis |  |  |  |
|  |  |  |  | Apparent Mw (kDa) | Oligomeric state | 2nd structure | Refolding ability | T<sub>m</sub> |  |
| mk2h_ΔM | 5.1 | ✓ | ✓ | 9.5 | 1.9 | α/β | ✓ | 71.3 |  |
| mk2h_ΔI | 5.1 | ✓ | ✓ | 9.6 | 1.9 | α/β | ✓ | 74.4 |  |
| mk2h_ΔL | 5.1 | ✓ | ✓ | 9.6 | 1.9 | α/β | ✓ | 76.2 |  |
| mk2h_ΔP | 5.1 | ✓ | ✓ | 9.7 | 1.9 | α/β | ✓ | 71.8 | ✓ (7DXU) |
| mk2h_ΔS | 5.2 | ✓ | ✓ | 13.1 | 2.5 | α/β | ✓ | 78.5 |  |
| mk2h_ΔY | 5.0 | ✓ | ✓ | 12.8 | 2.6 | α/β |  | 50.9 | ✓ (7DXV) |
| mk2h_ΔMIL | 5.1 | ✓ | ✓ | 10.4 | 2.1 | α/β |  | 54.6 | ✓ (7DXW) |
| mk2h_ΔMILPS | 5.1 | ✓ | ✓ | 10.8 | 2.1 | α/β |  | 50.5 | ✓ (7DXX/7DXY) |
| mk2h_ΔMILPY | 4.9 | ✓ | ✓ | 19.7 | 4.0 | random-coil | – | – | – |
| mk2h_ΔMILYS | 5.0 | ✓ | ✓ | 20.6 | 4.1 | random-coil | – | – | – |
| mk2h_ΔMILPYS | 5.0 | ✓ | ✓ | 20.3 | 4.1 | random-coil | – | – | ✓ (7DXZ/7DYC) |

**Table S11.**
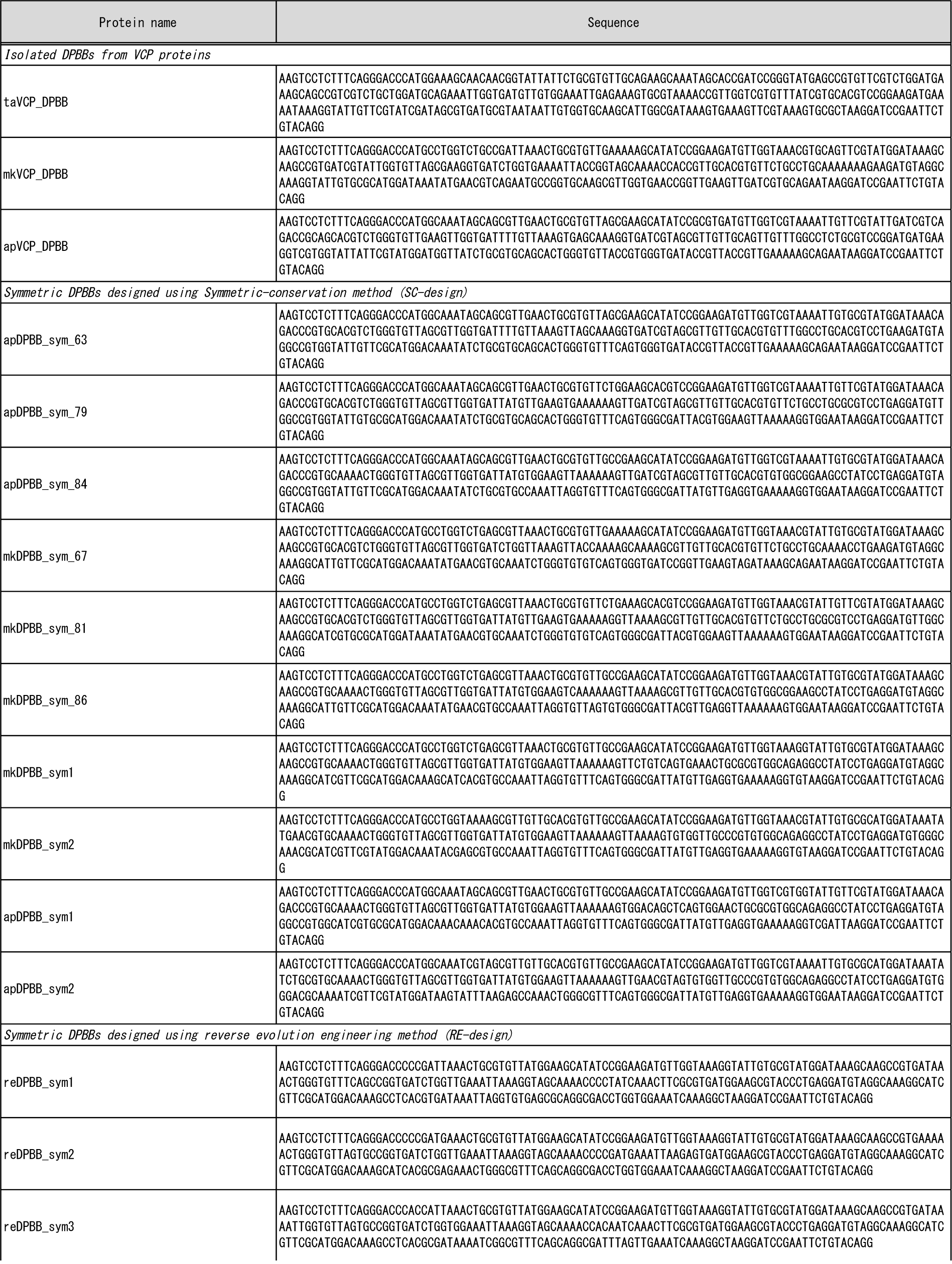
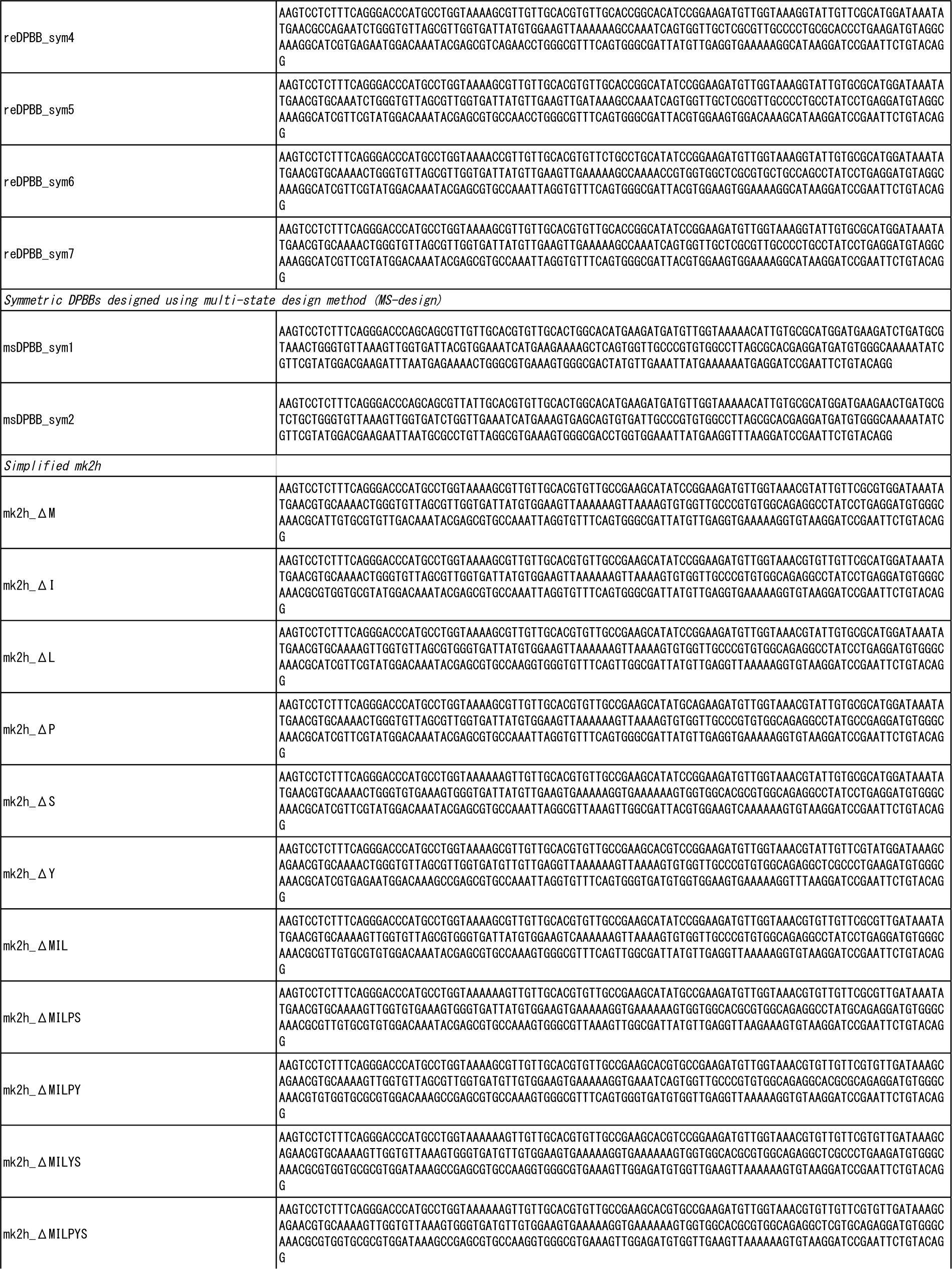
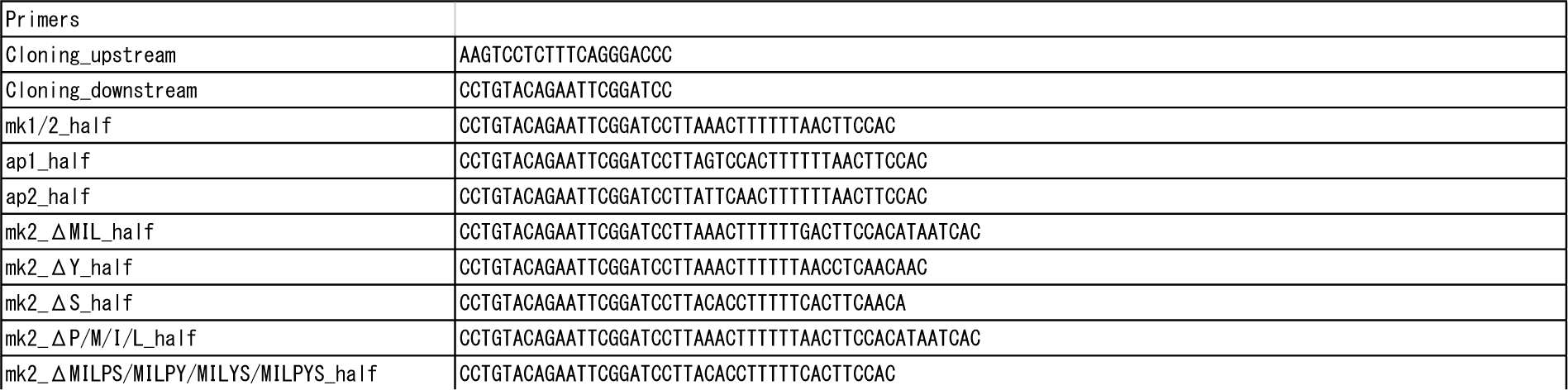
Gene and primer sequences.

**Table S12.**
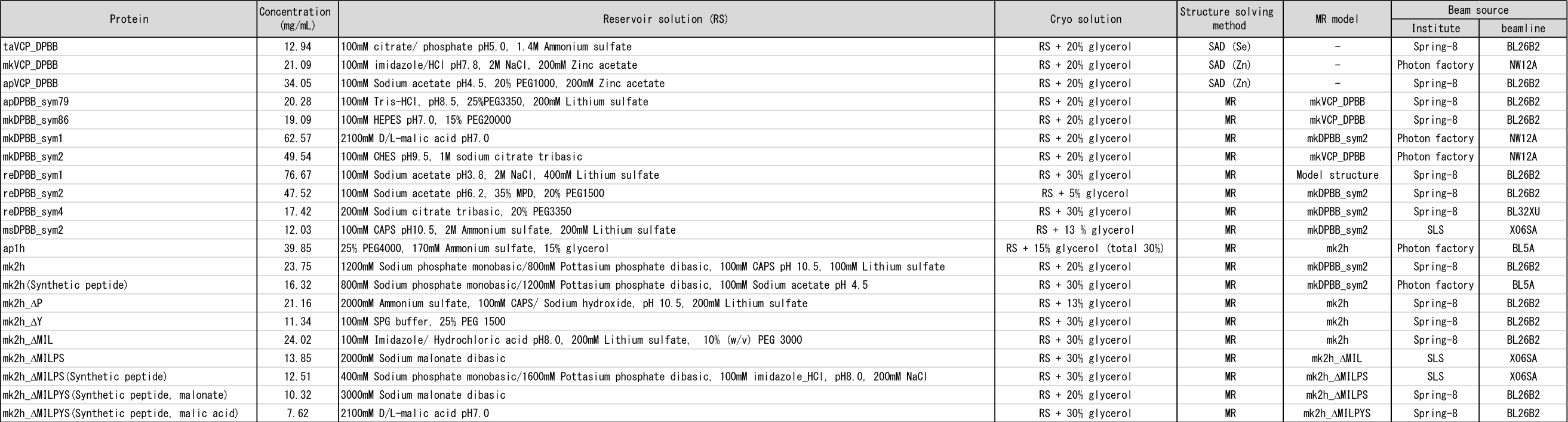
Summary of crystallization methods.

## Notes

### Competing Interest Statement

The authors have declared no competing interest.

